# Statistical Analysis of Spatial Expression Pattern for Spatially Resolved Transcriptomic Studies

**DOI:** 10.1101/810903

**Authors:** Shiquan Sun, Jiaqiang Zhu, Xiang Zhou

## Abstract

Recent development of various spatially resolved transcriptomic techniques has enabled gene expression profiling on complex tissues with spatial localization information. Identifying genes that display spatial expression pattern in these studies is an important first step towards characterizing the spatial transcriptomic landscape. Detecting spatially expressed genes requires the development of statistical methods that can properly model spatial count data, provide effective type I error control, have sufficient statistical power, and are computationally efficient. Here, we developed such a method, SPARK. SPARK directly models count data generated from various spatial resolved transcriptomic techniques through generalized linear spatial models. With a new efficient penalized quasi-likelihood based algorithm, SPARK is scalable to data sets with tens of thousands of genes measured on tens of thousands of samples. Importantly, SPARK relies on newly developed statistical formulas for hypothesis testing, producing well-calibrated *p*-values and yielding high statistical power. We illustrate the benefits of SPARK through extensive simulations and in-depth analysis of four published spatially resolved transcriptomic data sets. In the real data applications, SPARK is up to ten times more powerful than existing approaches. The high power of SPARK allows us to identify new genes and pathways that reveal new biology in the data that otherwise cannot be revealed by existing approaches.

## INTRODUCTION

Recent emergence of various spatially resolved transcriptomic technologies has enabled gene expression profiling with spatial localization information in tissues or cell cultures. Some of these techniques, such as MERFISH^1^ and seqFISH^2^, are based on single-molecular fluorescence *in situ* hybridization (smFISH)^3^. These smFISH based techniques can measure expression level for hundreds of genes with subcellular spatial resolution in a single cell. Some of these techniques, such as TIVA^4^, LCM^5^, Tomo-Seq^6^ and spatial transcriptomics through spatial barcoding^7^, are based on the next generation DNA sequencing. These DNA sequencing-based techniques can measure expression level for tens of thousands of genes on spatially organized tissue regions, each of which potentially consists of a couple hundred single cells. Some of these techniques, such as targeted *in situ* sequencing (ISS)^8^ and FISSEQ^9^, are based on *in situ* RNA sequencing. These RNA sequencing-based techniques can measure expression levels for the entire transcriptome with spatial information at a single cell resolution. These different spatially resolved transcriptomic techniques altogether have made it possible to study the spatial organization of transcriptomic landscape across tissue sections or within single cells, catalyzing new discoveries in many areas of biology^10, 11^.

In spatially resolved transcriptomic studies, identifying genes that display spatial expression pattern, which we simply refer to as SE analysis, is an important first step towards characterizing the spatial transcriptomic landscape. However, identifying SE genes is challenging both from a statistical perspective and from a computational perspective. From a statistical perspective, identifying SE genes requires the development of spatial statistical methods that can directly model raw count data generated from both smFISH based techniques and sequencing based techniques. Unfortunately, count based SE analysis methods are currently lacking. The only two existing approaches for SE analysis, SpatialDE^12^ and Trendsceek^13^, both transform count data into normalized data before analysis. However, analyzing normalized expression data can be suboptimal as this approach fails to account for the mean-variance relationship existed in raw counts, leading to a potential loss of power^14^. Indeed, similar loss of power has been well documented for methods that can only analyze normalized data in many other omics sequencing studies^15, 16^. Besides direct modeling of count data, identifying SE genes also requires the development of statistical methods that can produce well calibrated *p*-values to ensure proper control of type I error. However, some existing methods for SE analysis, such as SpatialDE^12^, rely on asymptotic normality and minimal *p*-value combination rule for constructing their hypothesis tests. Subsequently, these methods may fail to control for type I error at small *p*-values that are essential for detecting SE genes at the transcriptome-wide significance level. Failure of type I error control can lead to excessive false positives and/or substantial loss of power. From a computational perspective, while some spatial methods such as SpatialDE are based on linear models and are computationally efficient, some other spatial methods, in particular Trendsceek^13^, are built without a data generative model and compute non-parametric test statistics through computationally expensive permutation strategies that are not scalable to spatial transcriptomics data which are becoming increasingly large. Consequently, analyzing even moderate sized spatial transcriptomics data with hundreds of genes across hundreds of spatial locations can be a daunting task for these methods.

Here, we present a new method that address the above statistical and computational challenges. We refer to our method as Spatial PAttern Recognition via Kernels (SPARK). SPARK builds upon a generalized linear spatial model (GLSM)^17, 18^ with a variety of spatial kernels to accommodate count data generated from smFISH based or sequencing based spatial transcriptomics studies. With a newly developed penalized quasi-likelihood (PQL) algorithm^19, 20^, SPARK is scalable to analyzing tens of thousands genes across tens of thousands samples. Importantly, SPARK relies on a mixture of Chi-square distributions to serve as the exact test statistics distribution and further takes advantage of a recently developed Cauchy combination rule^21, 22^ to combine information across multiple spatial kernels for calibrated *p*-value calculation. As a result, SPARK properly controls for type I error at the transcriptome-wide level and is more powerful for identifying SE genes than existing approaches. We illustrate the benefits of SPARK through extensive simulations and applications to four published spatial transcriptomics studies. In the analysis of the real data sets, we show how SPARK can be used to identify new SE genes that reveal the importance of neuronal migration in the formation of the olfactory system as well as reveal the importance of immune system and cytoskeleton in tumor progression and metastasis.

## RESULTS

### Simulations

We provide an overview of SPARK in Materials and Methods, with technical details provided in Supplementary Text and a method schematic shown in Fig. 1A. Unlike Trendsceek, SPARK has an underlying data generative model which can be viewed as an extension of SpatialDE. However, unlike SpatialDE, SPARK models count data directly and relies on a proper statistical procedure to obtain calibrated *p*-values. A more detailed description of these methods is provided in Supplementary Text. We performed two sets of simulations to evaluate the performance of SPARK and compared it with two existing approaches, SpatialDE and Trendsceek. Simulation details are provided in Materials and Methods. Briefly, in the first set of simulations, for each scenario, we simulated 10,000 genes on 260 spatial locations (a.k.a. spots) in the mouse olfactory bulb data using parameters inferred from the real data. We examined both type I error control under the null hypothesis and power for identifying SE genes under common alternatives. In the null simulations, all genes are non-SE genes with expression levels randomly distributed across spatial locations without any spatial patterns (Fig. 1B). In the alternative simulations, 9,000 genes are non-SE genes, while 1,000 genes are SE genes whose expression levels display one of the three observed spatial patterns in the data (named as spatial pattern I, II and III; Fig. 1C). In the simulations, we varied noise variance to be either low, moderate or high, and varied the SE strength for SE genes to be either weak, moderate or strong.

**Figure 1:**
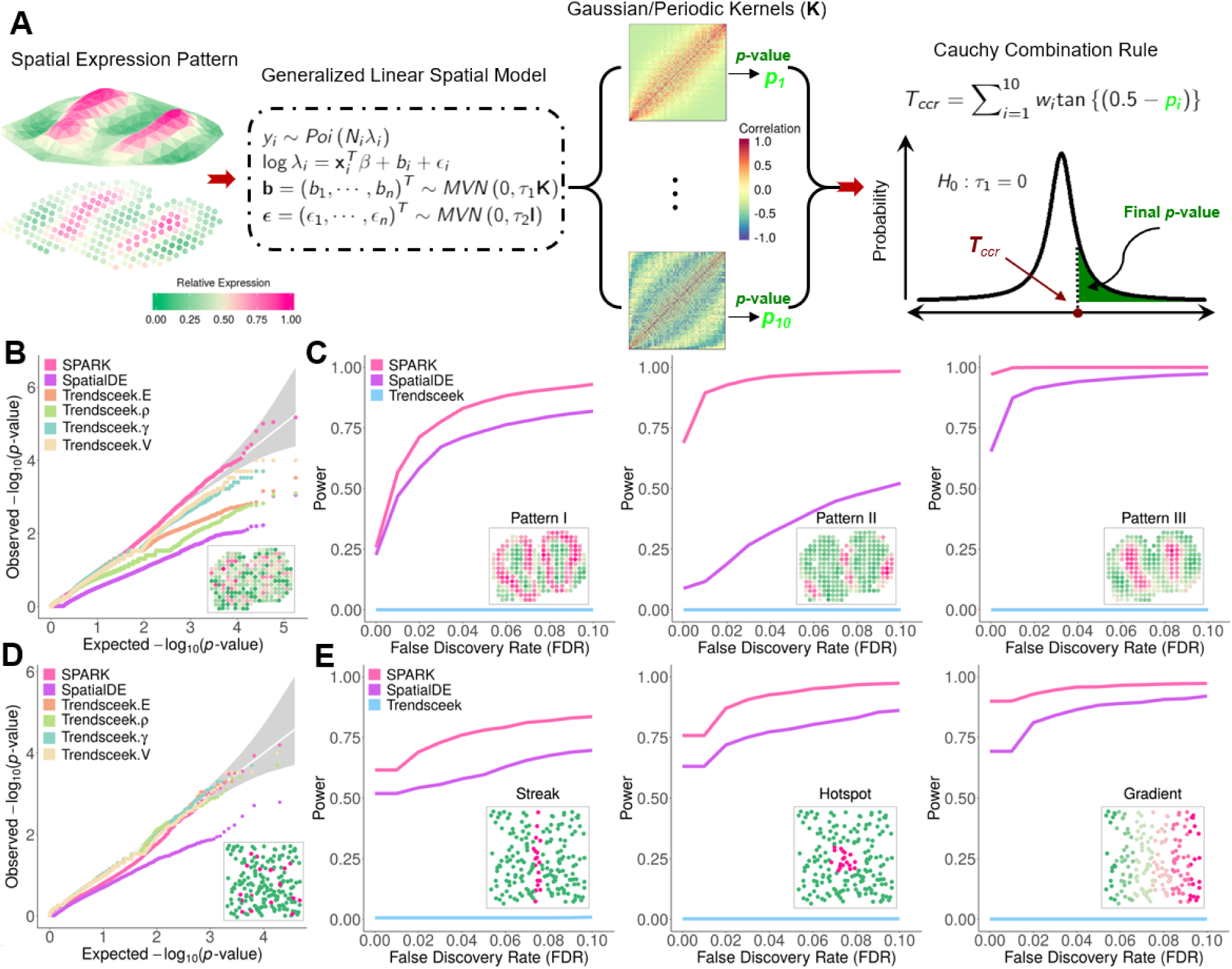
Method schematic of SPARK and simulation results. (**A**) Method schematic of SPARK. SPARK examines one gene at a time and models the gene expression measurements on spatial locations using the generalized linear spatial model (GLSM). To detect whether the gene shows spatial expression pattern, SPARK relies on a series of spatial kernels for pattern recognition and outputs a *p*-value for each spatial kernel using the Satterthwaite method that enables exact *p*-value computation. All these p-values from different spatial kernels are subsequently combined into a final SPARK *p*-value through the Cauchy combination rule. (**B**) Quantile-quantile plot of the observed −log10 *p*-values from different methods against the expected −log10 *p*-values under the null for the first set of null simulations based on the mouse olfactory bulb data. *p*-values are combined across ten simulation replicates. Simulations are performed under moderate noise (*τ*_2_ = 0.35). Compared methods include SPARK (pink), SpatialDE (purple), Trendsceek.E (light salmon) which is the Emark test of Trendsceek, Trendsceek. *ρ* (yellow-green) which is the Markcorr test of Trendsceek, Trendsceek. *γ* (light green) which is the Markvario test of Trendsceek, and Trendsceek.V (wheat) which is the Vmark test of Trendsceek. *p*-values from SPARK and some of the Trendsceek methods (e.g. Markvario and Vmark) are well calibrated. In contrast, *p*-values from SpatialDE, and to a lesser extent from the Emark and Markcorr tests of Trendsceek, are overly conservative and distributed below the expected diagonal line. Representative expression pattern for a null gene that does not show a spatial expression pattern is embedded inside the panel. (**C**) Power plots show the proportion of true positives (y-axis) detected by different methods at a range of false discovery rates (FDR; x-axis) for the first set of alternative simulations based on the mouse olfactory bulb data. Representative genes displaying each of the three spatial expression patterns I-III are embedded inside the panels. The proportion of true positives is averaged across ten simulation replicates. Simulations are performed under moderate noise (*τ*_2_ = 0.35) and moderate SE strength (threefold). Trendsceek (sky-blue) is the combined test of Trendsceek. (**D**) Quantile-quantile plot of the observed −log10 *p*-values from different methods against the expected −log10 *p*-values under the null for the second set of null simulations based on the SeqFISH data. *p*-values are combined across ten simulation replicates. Simulations are performed under moderate sample size (*n* = 200). *p*-values from SPARK and Trendsceek are well calibrated. In contrast, *p*-values from SpatialDE are overly conservative and distributed below the expected diagonal line. Representative expression pattern for a null gene that does not show a spatial expression pattern is embedded inside the panel. (**E**) Power plots show the proportion of true positives (y-axis) detected by different methods at a range of false discovery rates (FDR; x-axis) for the first set of alternative simulations based on the SeqFISH data. Representative genes displaying each of the three spatial expression patterns are embedded inside the panels. The proportion of true positives is averaged across ten simulation replicates. Simulations were performed under moderate fraction of marked cells (20%) and moderate SE strength (2 fold) for the hotspot and streak patterns, or under moderate SE strength (40% cells displaying expression gradient) for the linear gradient pattern. In all these simulations, SPARK properly controls for type I error and is more powerful than the other two methods for detecting genes with spatial expression patterns.

In the null simulations, we found that SPARK produces well-calibrated *p*-values at transcriptome-wide significance levels (Fig. 1B). Some Trendsceek test statistics (e.g. markvario and Vmark) also produce reasonably calibrated *p*-values while others (e.g. Emark statistics and markcorr statistics) yield slightly conservative *p*-values. In contrast, SpatialDE produces overly conservative *p*-values (Fig. 1B). The failure of SpatialDE in type I error control presumably is due to its use of an asymptotic Chi-square distribution in place of an exact distribution for *p*-value computation and/or its use of the *ad hoc* minimal *p*-value combination rule. The *p*-value calibration results for different methods are consistent across simulation settings and across a range of noise variance levels (Fig. S1). Because some methods fail to control for type I error, in the alternative simulations, we measured power based on false discovery rate (FDR) to ensure fair comparison among methods. In the alternative simulations, we found that SPARK is more powerful than the other two methods across a range of FDR cutoffs (Fig. 1C) and across a range of parameter settings (Figs. S2 and S3). The power performance of SPARK is followed by SpatialDE, while Trendsceek does not fare well in any of the alternative simulations. For example, in the easy setting where the noise variance is moderate and the spatial expression pattern strength is moderate, SPARK identified 860, 968, 1000 SE genes at an FDR of 5% for spatial patterns I-III, respectively (Fig. 1C). The power of SPARK is 16.5%, 168.1% and 5.5% higher than that of SpatialDE (which identified 738, 361 and 948 SE genes), for the three spatial patterns, respectively. In contrast, Trendsceek was only capable of identifying less than three SE genes (with the detailed number varying depending on the spatial pattern and the random seed Trendsceek used). In the more challenging setting where the noise variance is high and the spatial expression pattern strength is moderate, SPARK identified 540, 872, 982 SE genes at an FDR of 5% for spatial patterns I-III, respectively (Fig. S3). The power of SPARK is 38.8%, 685.6% and 36.6% higher than that of SpatialDE (which identified 389, 111 and 719 SE genes), for the three spatial patterns, respectively. In contrast, Trendsceek was only capable of identifying less than two SE genes.

Because of the extremely poor performance of Trendsceek in the first set of simulations, to rule out the possibility that our first simulations were somehow biased against Trendsceek, we compared different methods on a second set of simulations performed fully based on the original Trendsceek paper^13^. Simulation details are provided in Materials and Methods. Briefly, we first randomly simulated the spatial locations for a fixed number of cells through a spatial Poisson process. We then generated 1,000 genes in the simulated data, which were all non-SE genes in the null simulations and consisted of 100 SE genes and 900 non-SE genes in the alternative simulations. For non-SE genes, the expression measurements from the real data were randomly assigned to the simulated cells regardless of their spatial locations (Fig. 1D). For SE genes, the expression measurements from the real data were assigned to the simulated cells to display three distinct spatial patterns (Fig.1E): cells in a focal area showed higher expression measurements than the remaining cells (Hotspot pattern), cells in a streak area showed higher expression measurements than the remaining cells (Streak pattern), or cells tend to show gradually reduced expression measurements when they are further away from the streak (Gradient pattern). In the simulations, we varied the number of cells (*n* = 100, 200 or 500), the SE strength (low, moderate or high; measured by the fold change between cells inside and outside the focal/streak area in the first two spatial patterns and by the fraction of cells displaying expression gradient for the third spatial pattern), as well as the fraction of cells in the focal/streak area for the first two spatial patterns. We applied all three methods to the simulated data. The comparison results are largely consistent with the results obtained from the first set of simulations. Specifically, under the null, both SPARK and Trendsceek produce well-calibrated *p*-values, while SpatialDE does not (Fig. 1D). Under the alternative, SPARK is more powerful than the other two methods across a range of FDR cutoffs (Fig. 1E) in almost all parameter settings (Figs. S4-S7). The power performance of SPARK is followed by SpatialDE, while Trendsceek does not fare well, even though the power of Trendsceek is largely consistent with its performance shown in the original Trendsceek paper^13^. For example, when the SE strength is moderate and the cell number equals to 200, SPARK identified 94, 78 and 96 SE genes at an FDR of 5% for the Hotspot, Streak and Gradient patterns, respectively (Fig. 1E). The power of SPARK is 20.5%, 31.7% and 9% higher than that of SpatialDE (which identified 78, 60 and 88 SE genes) for the three patterns, respectively. In contrast, Trendsceek was only capable of identifying less than two SE genes, consistent with original study^13^. As expected, the power of all methods increases with increasing SE strength and increasing sample size. For example, when SE strength is high and the cell number is large (*n* = 500), consistent with^13^, Trendsceek detected 9, 7 and 12 SE genes for the Hotspot, Streak and Gradient patterns (Figs. S7 and S8), respectively, again consistent with^13^. However, in such setting, both SPARK and SpatialDE reach 100% power and can detect all SE genes. Overall, the two sets of simulations suggest that SPARK produces well-calibrated *p*-values while being more powerful than the other two methods in detecting SE genes.

### Olfactory Bulb Data

We applied SPARK to analyze four published data, including two data obtained through spatial transcriptomics sequencing and two data through smFISH (details in Materials and Methods). The first data we examined is a mouse olfactory bulb data^7^, consisting of gene expression measurements for 11,274 genes on 260 spots. Consistent with simulations, both SPARK and Trendsceek produce calibrated *p*-values under permuted null, while SpatialDE does not (Fig.2A). SPARK also identified more SE genes compared to SpatialDE and Trendsceek across a range of FDRs (Figs. 2B and S9). For example, at an FDR of 5%, SPARK identified 772 SE genes, which is ∼10-fold more than that detected by SpatialDE (which identified 67, among which 62 are overlapped with SPARK; Figs. 2B and 2E); Trendsceek was unable to detect any SE genes in the data, even though we tried ten different random seeds for the method.

**Figure 2:**
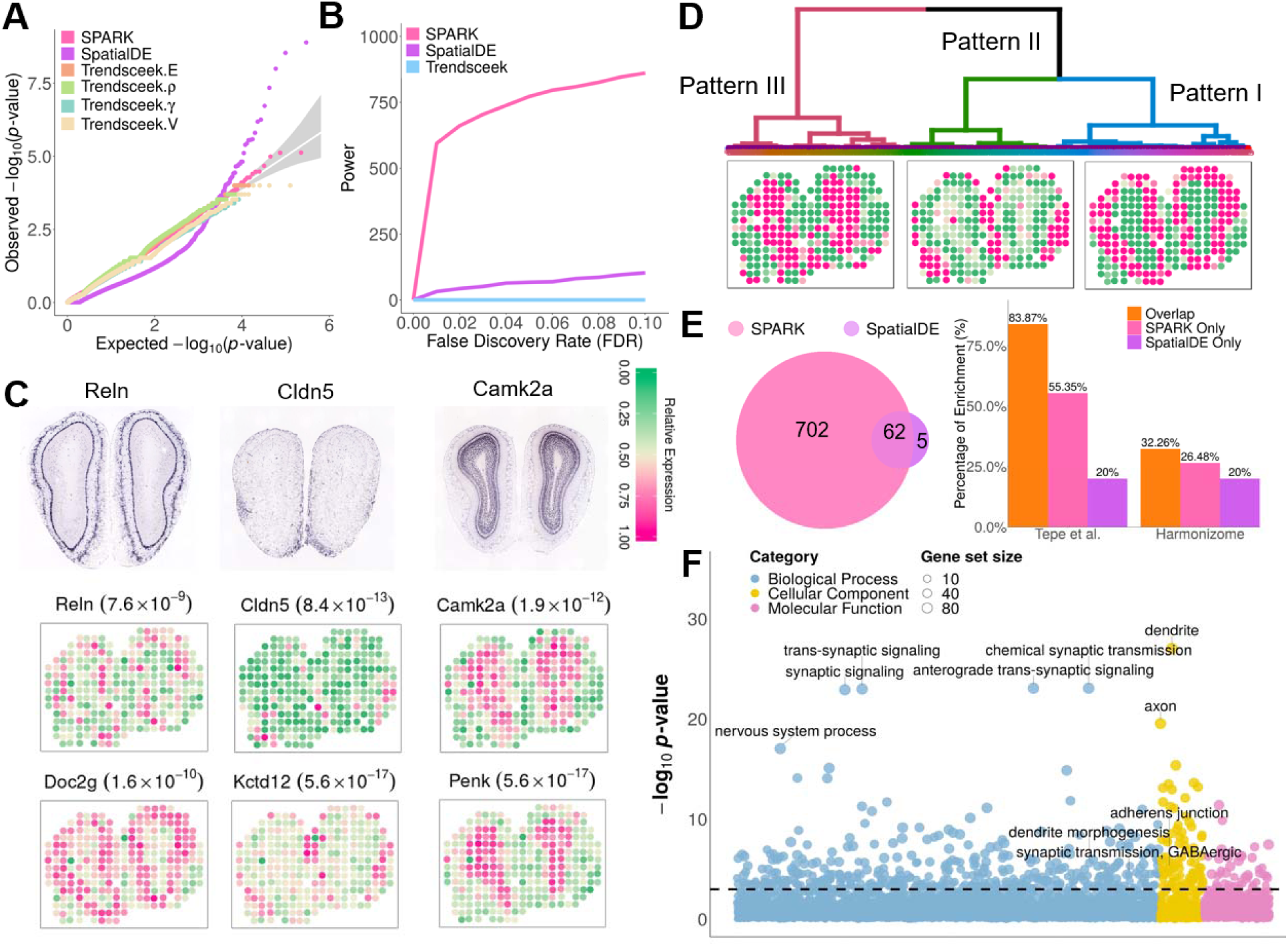
Analyzing the mouse olfactory bulb data. (**A**) Quantile-quantile plot of the observed −log10 *p*-values from different methods are plotted against the expected −log10 *p*-values under the null in the permuted data. *p*-values are combined across ten permutation replicates. Compared methods include SPARK (pink), SpatialDE (purple), Trendsceek.E (light salmon) which is the Emark test of Trendsceek, Trendsceek.*ρ* (yellow-green) which is the Markcorr test of Trendsceek, Trendsceek.*γ* (light green) which is the Markvario test of Trendsceek, and Trendsceek.V (wheat) which is the Vmark test of Trendsceek. *p*-values from SPARK and various Trendsceek methods are well calibrated. In contrast, *p*-values from SpatialDE are overly conservative at large values and are overly anti-conservative at small values. (**B**) Power plot shows the number of genes with spatial expression pattern (y-axis) identified by different methods at a range of false discovery rates (FDRs; x-axis). Across a range of FDRs, SPARK detected more genes with spatial expression pattern than SpatialDE, while Trendsceek (sky-blue) which is the combined test of Trendsceek, detected almost none. (**C**) *In situ* hybridization of three representative genes (*Reln*, *Cldn5*, and *Camk2a*) obtained from the database of the Allen Brain Atlas. *Reln* is spatially expressed in the mitral layer and glomeruli layer. *Cldn5* is spatially expressed in the nerve layer. *Camk2a* is spatially expressed in the granular layer. Spatial expression pattern for the same three genes (*Reln*, *Cldn5*, and *Camk2a*) in the spatial transcriptomics data, along with their *p*-values from SPARK (inside parenthesis). Color represents relative gene expression level (purple: high; green: low). These genes are only identified by SPARK, but not by the other two methods. Spatial expression patterns for three additional known marker genes (*Doc2g*, *Kctd12*, and *Penk*) in the spatial transcriptomics data, along with their *p*-values from SPARK (inside parenthesis). These genes are previously known molecular markers for different layers in the mouse olfactory bulb: *Doc2g* for mitral layer; *Kctd12* for nerve layer; and *Penk* for granular layer. (**D**) Three distinct spatial expression patterns summarized based on the 772 SE genes that are identified by SPARK, along with dendrogram displaying the clustering of these three main patterns. (**E**) Venn diagram shows the overlap between SE genes identified by SPARK and SpatialDE. Bar plot shows the percentage of SE genes identified by SPARK (orange/pink) or SpatialDE (orange/purple) that are also validated in two gene lists, one from a literature (left) and the other from the Harmonizome database (right). The orange bar represents the percentage of SE genes identified by both SPARK and SpatialDE that are in either of the gene lists; the pink bar represents the percentage of unique SE genes identified by SPARK that are in either of the gene lists; the purple bar represents the percentage of unique SE genes identified by SpatialDE that are in either of the gene lists. In both gene lists, SE genes identified only by SPARK show a higher percentage of overlap with existing gene lists than SE genes identified only by SpatialDE. (**F**) Bubble plot shows – log10 *p*-values for pathway enrichment analysis on SE genes obtained by SPARK. Gene sets are colored by three categories: GO biological process (blue), GO molecular function (purple), and GO cellular component (yellow).

We carefully examined the SE genes and found that most SE genes only detected by SpatialDE tend to have close to zero expression levels (Fig. S10) and appear to locate on either one or two spots (Fig. S11), suggesting potentially false signals. In contrast, the SE genes detected only by SPARK generally have comparable expression levels to the SE genes detected by both methods (Fig. S10). To assess the quality of the SE genes identified by SPARK, we performed clustering on the 772 SE genes and obtained three major spatial expression patterns (dendrogram in Fig. 2D; UMAP visualization in Fig. S12): one representing the mitral cell layer (Pattern I); one representing the glomerular layer (Pattern II); and one representing the granular cell layer (Pattern III); all clearly visualized via three previously known marker genes for the three layers, *Doc2g*, *Kctd12* and *Penk*^7^ (Fig. 2C). For each spatial pattern, we ranked genes only detected by SPARK based on their *p*-values and obtained 20 genes with increasing *p*-values from the ranked list as representative examples (Figs. S13-S15). Almost all these genes show clear spatial expression pattern, cross validated by *in situ* hybridization data provided by the Allen Brain Atlas (Fig. 2C), confirming the higher power of SPARK.

We provide three additional lines of evidence to validate the SE genes detected by SPARK. First, we examined the highlighted marker genes in the olfactory system presented in the original study. The list of highlighted marker genes, while is not necessarily the complete list of all marker genes, at least represents the likely best set of genes one can obtain that are both biologically important for the data and are detectable in the data. Importantly, SPARK detected 8 of 10 such highlighted mitral cell layer (MCL) enriched genes; while SpatialDE only detected 3 and Trendsceek detected none (Fig. S16). Second, we obtained a list of 2,030 cell type specific marker genes identified in a recent single cell RNAseq study in the olfactory bulb^23^. Reassuringly, 55% of the unique SE genes identified by SPARK are in the marker list, while only 20% of the unique SE genes identified by SpatialDE are in the same list (Fig. 2E). Third, we obtained a list of 3,262 genes that are related to the olfactory system from the Harmonizome database^24^. Again, 26% of the unique SE genes identified by SPARK are in the Harmonizome list, while only 20% of the unique SE genes identified by SpatialDE are in the same list (Fig. 2E). These three additional validation analyses provide convergence support for the higher power of SPARK.

Finally, we performed functional enrichment analyses of SE genes identified by SPARK and SpatialDE (details in Materials and Methods). A total of 1,023 GO terms (Fig. 2F) and 79 KEGG pathways were enriched in the SE genes identified by SPARK at an FDR of 5%, while only 87 GO terms (overlap = 64; Fig. S17A) and 2 KEGG pathways (overlap = 2; Fig. S17B) were enriched in the SE genes identified by SpatialDE (Table S1; Fig. S17C). Many enriched GO terms or KEGG pathways identified only by SPARK are directly related to the synaptic organization and olfactory bulb development. For example, olfactory lobe development is a highly enriched GO term detected only by SPARK (GO:0021988; SPARK: *p*-value = 5.81×10^−3^; SpatialDE: *p*-value = 1.21×10^−1^). Oxytocin signaling pathway is a highly enriched KEGG pathway detected only by SPARK (KEGG: mmu04921; SPARK: *p*-value = 1.59×10^−9^; SpatialDE: *p*-value = 2.15×10^−1^) and is known to modify olfactory response^26^. The newly identified GO term and KEGG pathway enrichment highlights the benefits of running SE analysis with SPARK.

A further enrichment analysis using SE genes in Patterns I-III separately provide additional biological insights. SPARK identified a total of 489, 714, and 684 enriched GO terms for Patterns I-III, respectively; while SpatialDE only identified 171 (overlap = 96), 275 (overlap = 177), and 22 (overlap = 22; Fig. S18, Table S1). For example, in Pattern I, the enriched GO term of glutamatergic synaptic transmission is only detected by SPARK (GO:0035249; SPARK: *p*-value = 1.06×10^−6^; SpatialDE: *p*-value = 2.14×10^−3^; Fig. S19; Table S1), and supports the functional role of the synaptic organization in the mitral cell layer^27^. One representative gene in this GO term is *Reln*, which is only identified by SPARK (Fig. 2C). *Reln* encodes the protein Reelin expressed in mitral cells and promotes tangential to radial migration transition^28^. In Pattern II, the enriched GO term of cell junction assembly is only detected by SPARK (GO:0034329; SPARK: *p*-value = 1.22×10^−9^; SpatialDE: *p*-value = 1.48×10^−2^; Fig. S20; Table S1), and supports the critical role of cell junction and synaptic connection in the nerve layer^29^. One representative gene in this GO term is *Cldn5*, which is only identified by SPARK (Fig. 2C). *Cldn5* is known to be enriched in the olfactory nerve layer and is critical for cell-cell adhesion^30^. In Pattern III, the enriched GO term of dendrite morphogenesis is only detected by SPARK (GO:0048813; SPARK: *p*-value = 6.53×10^−13^; SpatialDE: *p*-value = 8.39×10^−3^; Fig. S21; Table S1), and supports the importance of dendritic morphogenesis in the development of granular layer^31^. One representative gene in this GO term is *Camk2a*, which is again only identified by SPARK (Fig. 2C). *Camk2a* is enriched in the granular cell layer and encodes a protein that belongs to the Ca(2+)/calmodulin-dependent protein kinases subfamily which is crucial for several key aspects of synaptic and dendritic plasticity^32^. Overall, these new GO terms/KEGG pathways and SE genes that are only identified by SPARK reveal new biology in the data that otherwise cannot be discovered by existing methods.

### Breast Cancer Data

The second data we examined is a human breast cancer biopsy study^7^, which contains 5,262 genes measured on 250 spots. Consistent with simulations, both SPARK and Trendsceek produce calibrated *p*-values under permuted null, while SpatialDE does not (Fig. 3A); SPARK identified more SE genes compared to SpatialDE and Trendsceek across a range of FDRs (Figs. 3B and S22). For example, at an FDR of 5%, SPARK identified 290 SE genes, which is ∼3-fold more than that detected by SpatialDE (which identified 115, among which 85 are overlapped with SPARK; Figs. 3B and 3D). In contrast, Trendsceek only identified at most 15 SE genes across ten different random seeds. Consistent with the olfactory bulb study, we also found that the SE genes only detected by SpatialDE tend to have low expression levels, suggesting of potential false positives. In contrast, the SE genes detected only by SPARK generally have comparable expression levels to the SE genes detected by both methods (Fig. S23). To assess the quality of the SE genes identified only by SPARK, we obtained 20 genes with increasing *p*-values from the ranked list as representative examples (Fig. S24). Again, most of these genes show clear spatial expression pattern, confirming the higher power of SPARK.

**Figure 3:**
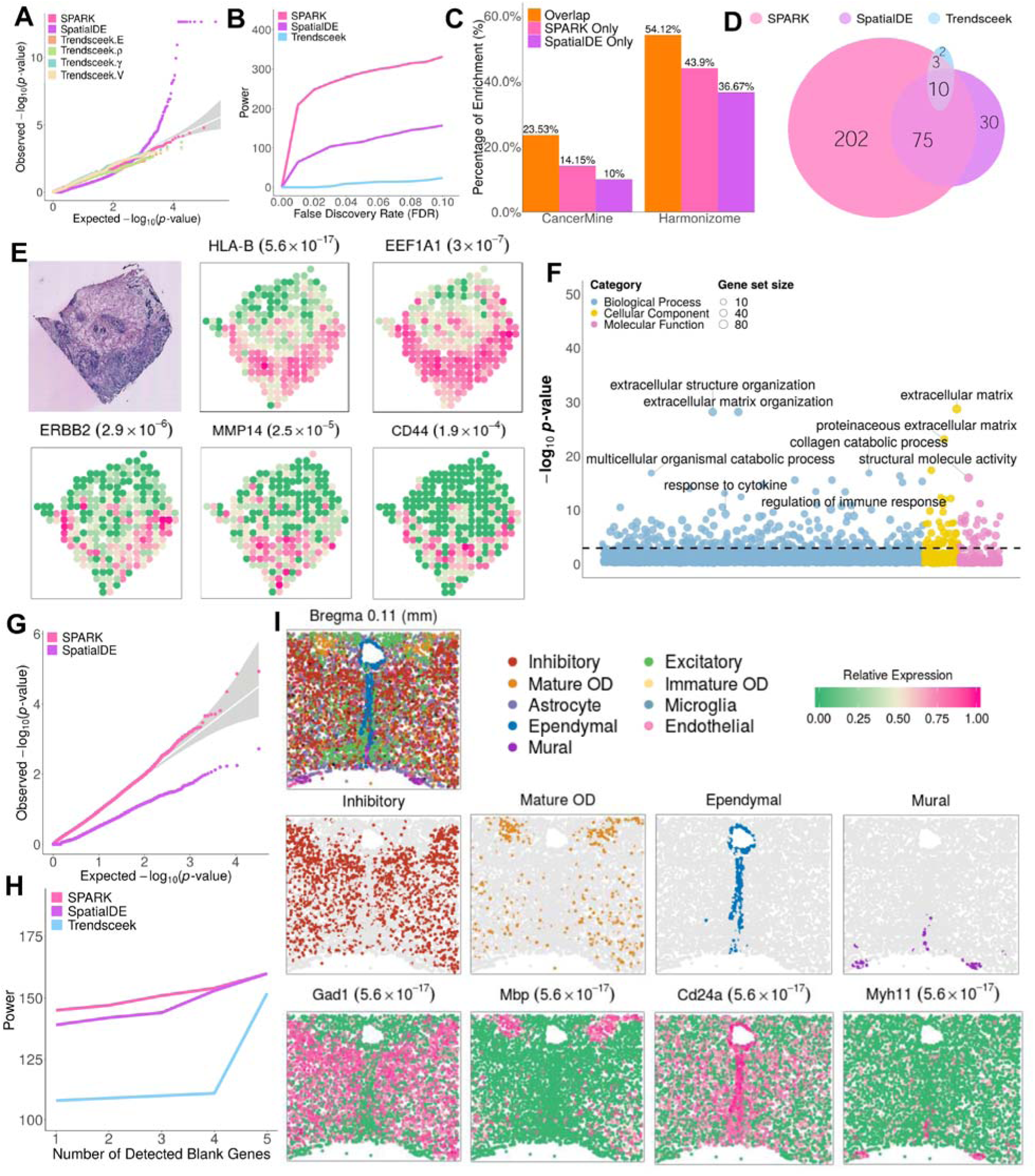
Analyzing the human breast cancer data and the mouse hypothalamus data. (**A**) Quantile-quantile plot of the observed −log10 *p*-values from different methods are plotted against the expected −log10 *p*-values under the null in the permuted data. *p*-values are combined across ten permutation replicates. Compared methods include SPARK (pink), SpatialDE (purple), Trendsceek.E (light salmon) which is the Emark test of Trendsceek, Trendsceek.*ρ* (yellow-green) which is the Markcorr test of Trendsceek, Trendsceek. *γ* (light green) which is the Markvario test of Trendsceek, and Trendsceek.V (wheat) which is the Vmark test of Trendsceek. *p*-values from SPARK and various Trendsceek methods are approximately well calibrated. In contrast, *p*-values from SpatialDE are overly conservative at large values and are overly anti-conservative at small values. (**B**) Power plot shows the number of genes with spatial expression pattern (y-axis) identified by different methods at a range of false discovery rates (FDRs; x-axis). Across a range of FDRs, SPARK detected more genes with spatial expression pattern than SpatialDE, while Trendsceek (sky-blue) which is the combined test of Trendsceek, detected only a few. (**C**) Bar plot shows the percentage of SE genes identified by SPARK (orange/pink) or SpatialDE (orange/purple) that are also validated in two gene lists, one from the CancerMine database (left) and the other from the Harmonizome database (right). The orange bar represents the percentage of SE genes identified by both SPARK and SpatialDE that are in either of the gene lists; the pink bar represents the percentage of unique SE genes identified by SPARK that are in either of the gene lists; the purple bar represents the percentage of unique SE genes identified by SpatialDE that are in either of the gene lists. In both gene lists, SE genes identified only by SPARK show a higher percentage of overlap with existing gene lists than SE genes identified only by SpatialDE. (**D**) Venn diagram shows the overlap between SE genes identified by SPARK and SpatialDE. (**E**) Spatial expression pattern for five genes (*HLA-B*, *EEF1A1*, *ERBB2*, MMP14, and *CD44*) that are only identified by SPARK but not by the other two methods. The *p*-values for the five genes from SPARK are shown inside parenthesis. Color represents relative gene expression level (purple: high; green: low). For reference, the hematoxylin and eosin (H&E) staining on an adjacent section is shown in the top left panel. The dark staining in the H&E panel represents potential tumors. The H&E panel is reproduced based on the reference^7^. These five genes are previously known molecular markers associated with tumor induced immune response (*HLA-B*), growth factor (ERBB2), or metastasis (*EEF1A1, MMP14* and *CD44*). (**F**) Bubble plot shows –log10 *p*-values for pathway enrichment analysis on SE genes obtained by SPARK. Gene sets are colored by categories: GO biological process (blue), GO molecular function (purple), and GO cellular component (yellow). (**G**) Quantile-quantile plot of the observed −log10 *p*-values from different methods are plotted against the expected −log10 *p*-values under the null in the permuted data. *p*-values are combined across one hundred permutation replicates. Compared methods include SPARK (pink) and SpatialDE (purple). Results for Trendsceek are not included here due to computational issue. (**H**) Power plot shows the number of genes with spatial expression pattern (y-axis) identified by different methods vs the number of blank control genes identified at the same threshold (x-axis). SPARK detected more genes with spatial expression pattern than SpatialDE and Trendsceek (sky-blue) across various numbers of false discoveries. Color represents relative gene expression level (purple: high; green: low). (**I**) Spatial distribution of all major cell classes on the 1.8-mm by 1.8-mm imaged slice from a single female mouse (Bregma +0.11). Cells are colored by cell classes shown in the legend, where the cell class information are obtained from the reference^42^. Spatial distribution of four main cell classes. The spatial distributions of the remaining five cell classes are shown in a Supplementary Figure. The cell classes are represented by colored dots while the background of all other cells is shown as gray dots. Spatial expression pattern for four representative genes (*Gad1*, *Mbp*, *Cd24a,* and *Myh11*) that are identified by all three methods. The *p*-values for the four genes from SPARK are shown inside parenthesis.

We provide three additional lines of evidence to validate the SE genes detected by SPARK. First, we examined the 14 cancer relevant genes highlighted in the original study. Importantly, SPARK detected 10 of them while SpatialDE detected 7 and Trendsceek detected two (Fig. S25). Both SpatialDE and Trendsceek missed three of these previously well-known cancer relevant genes (*SCGB2A2*, *KRT17* and *MMP14*). Second, we collected a list of 1,144 genes that are previously known to be relevant to breast cancer through literature based on the CancerMine database^33^. 14% of SE genes uniquely identified by SPARK are in the list while only 10% by SpatialDE are in the list (Fig. 3C). For example, the well-known proto-oncogene *ERBB2* gene has tens of thousands of previous literature support on breast cancer but it can only be identified by SPARK (Fig. 3E). Third, we collected a list of 3,538 genes that are relevant to breast cancer based on the Harmonizome database^24^. Again, 44% of SE genes uniquely identified by SPARK are in the list while only 37% by SpatialDE are in the list (Fig. 3C). Overall, these three additional lines of evidence provide convergent support for the higher power of SPARK.

We performed functional enrichment analysis with GO term and KEGG pathways. At an FDR of 5%, SPARK identified 542 GO terms and 20 KEGG pathways (Fig. 3F; Table S2) while SpatialDE identified 266 GO terms (overlap = 191) and 3 KEGG pathways (overlap = 3; Fig. S26; Table S2). Many enriched gene sets discovered only by SPARK are related to extracellular matrix organization and immune responses (Figs. S26A-S26C; Table S2). For example, the GO term of response to cytokine is only identified by SPARK (GO:0034097; SPARK: *p*-value = 5.58×10^−10^; SpatialDE: not enriched); cytokines are released in response to immunity and can function to inhibit cancer development^34^. One representative gene in this GO term is *HLA-B*, which is a member of the human leukocyte antigen (HLA) complex^35^. *HLA-B* and five other HLA members are only detected by SPARK and are all expressed in the areas of ductal cancer (Figs. 3E and S24), suggesting a potential tumor-associated local immune response. As another example, the GO term of *immune effector process* is only identified by SPARK (GO:0002252; SPARK: *p*-value = 1.03×10^−3^; SpatialDE: not enriched); the number of immune effector cells plays an important role of cancer immunotherapy to block the tumor immune evasion and to restore immune surveillance^36^. One representative gene related to this GO term is *EEF1A1*, which is only detected by SPARK and is previously known to be upregulated in breast cancer samples^37^. *EEF1A1* is highly expressed in the cancer area with moderate expression in the rest areas (Fig. 3E) and such spatial expression pattern is consistent with the previous hypothesis that *EEF1A1* promotes tumor cell motility and subsequently metastasis through its influence in the cytoskeleton organization^38^. As last example, the GO term of *cell-substrate adherens junction* is only identified by SPARK (GO:0005924; SPARK: *p*-value = 1.61×10^−6^; SpatialDE: 5.97×10^−2^); the adhesion protein junctional adhesion molecule-A regulates epithelial cell morphology and migration, and its over-expression has recently been linked with increased risk of metastasis in breast cancer patients^39^. The example gene related to this GO term is *CD44* (Fig. 3E), which encodes a cell-surface glycoprotein involved in tumors metastasis. The interaction between *CD44* and matrix metalloproteinases (MMP) members such as *MMP2* and *MMP14* has been discovered in many cancer cell types^40, 41,^. It has been hypothesized that MMP-induced *CD44* cleave is associated with enhanced cell migration, thus facilitating metastasis. The coordinated expression of *MMP2*, *MMP14* and *CD44* in the cancer area of the studied sample (Figs. 3E and S24) highlights the importance of extracellular matrix in the process of metastasis. These new GO terms/KEGG pathways and SE genes that are only identified by SPARK again reveal new biology in the data that otherwise cannot be discovered by existing methods.

### Hypothalamus Data

The third data we examined is a MERFISH data collected on the preoptic area of the mouse hypothalamus^42^. The data contains 160 genes measured on 4,975 single cells with known spatial locations (Fig. 3I), and 155 out of these 160 genes were selected in the original study as they are makers of distinct cell populations or relevant to various neuronal functions of the hypothalamus. Besides these likely true positive genes, a total of 5 blank control genes that were also included in the original study to serve as negative controls. In the analysis, consistent with simulations, we found that SPARK produces calibrated *p*-values under permuted null, while SpatialDE does not (Fig. 3G). Note that we did not apply Trendsceek to the permuted null here due to computational reasons: it takes Trendsceek over 48 hours to analyze even one gene in this data. Also consistent with simulations, the QQ-plot of *p*-values from different methods suggest that both SpatialDE and SPARK are more powerful than Trendsceek (Fig. S27A). Because this data contains 5 negative control genes and 155 likely positive genes, we directly compared power of different methods based on the number of SE genes identified given a fixed number of negative control genes identified (Fig. 3H). The power comparison results again support a higher power of SPARK. For example, conditional on only one blank control gene being detected (i.e. one false positive), SPARK identified 145 SE genes, which is 6 more than that detected by SpatialDE (which identified 139, among which 138 are overlapped with SPARK; Figs. 3H and S27B). The performance of SPARK and SpatialDE is followed by Trendsceek, which identified 108 SE genes, among which 103 are overlapped with SPARK.

A careful examination suggests that almost all SE genes identified by SPARK show clear spatial expression pattern as one would expect. For example, we display 9 major cell classes in hypothalamus (Figs. 3I and S28A) along with 9 marker genes^42^ (Fig. S28B). Importantly, all four SE genes only identified by SPARK are closely related to the neuronal functions of the hypothalamus. Specifically, *Grpr* encodes a multiphase membrane protein that functions as a receptor for gastrin-releasing peptide. *Grpr* has been recently shown to mediate an antidepressant-like effect in mouse model and could be potentially served as a new therapeutic target of depression^43^. *Avpr1a* encodes a receptor for arginine vasopressin, which is a neurohypophysial hormone involved in the regulation of adrenocorticotropic hormone released from the pituitary^44^. A recent study has shown that the blocking AVPR1A may improve social communication in autism spectrum disorder^45^. *Chat* encodes an enzyme which catalyzes the biosynthesis of the neurotransmitter acetylcholine. Polymorphisms in *Chat* have been associated with Alzheimer’s disease and mild cognitive impairment^46^. *Nup62cl* itself is a protein coding gene related to structural constituent of nuclear pore. Previous studies have found that *Nup62cl* is co-expressed with *Ghrh*^42^, which is the gene coding the growth hormone-relating hormone (GHRH) secreted by the hypothalamus that further stimulates the synthesis and release of growth hormone (GH) in pituitary^47^. These important genes missed by other methods highlight the power of SPARK.

### Hippocampus Data

The final data we examined is from a mouse hippocampus study^48^. This is a small seqFISH data that contains 249 genes measured on 131 single cells with known spatial locations (Fig. S29A). These 249 genes include 214 genes that were selected in the original study as transcription factors and signaling pathway components and 35 remaining genes that are previously known cell identity markers. In the analysis, consistent with simulations, both SPARK and Trendsceek produce calibrated *p*-values under permuted null, while SpatialDE yields conservative *p*-values (Fig. S29B); SPARK again identified more SE genes compared to SpatialDE and Trendsceek across a range of FDRs (Figs. S29C and S29D). For example, at an FDR of 5%, SPARK identified 17 SE genes; while SpatialDE and Trendsceek identified 11 (all overlap with SPARK) and 4 (one overlap with SPARK) SE genes, respectively (Figs. S30-S32). The 11 SE genes identified by both SpatialDE and SPARK show clear spatial expression patterns (Fig. S31), so are the 6 SE genes identified only by SPARK (Fig. S32). The 3 SE genes only detected by Trendsceek tend to express uniformly highly in most cells and show less obvious spatial pattern (Fig. S33). The higher number and apparent spatial expression pattern of SE genes identified by SPARK support its higher power.

We carefully examined all six SE genes that are only identified by SPARK. Four of them are cell identity markers: *FoxO1* and *Slc17a8* for glutamatergic neurons; *igtp* for GABAergic neurons; and *opalin* for Oligodendrocytes^49^. All of them are closely related to neuronal functions in hippocampus. For example, the spatial expression pattern of *FoxO1* detected by SPARK is consistent with the previous observation that it is highly enriched in the ventral CA3 area of the hippocampus as well as in the amygdalohippocampal region^50, 51^. *FoxO1* is activated in hippocampal progenitor stem cells following cortisol exposure to prenatal stress and mediates the negative effect of stress on neurogenesis^52^. Besides these four marker genes, the remaining two genes are *pou4f1* and *gfi1*, both of which encode neural transcription factors and play important roles in the sensory nervous system development^53, 54^. These important genes that are missed by other methods again highlight the power of SPARK.

## DISCUSSION

We have presented a new computational method, SPARK, for identifying genes with spatial expression patterns in spatially resolved transcriptomic studies. Compared with existing approaches, SPARK is computationally scalable, produces well-calibrated *p*-values for type I error control, and is more powerful in identifying SE genes. We have illustrated the benefits of SPARK through extensive simulations and in-depth analysis of four real data sets.

Different from previous literatures in spatial statistics, SPARK incorporates a data generative model and relies on a model-based hypothesis test framework for spatial pattern detection. The data generative model in SPARK distinguishes it from previous spatial data exploratory tools that rely on variogram or semi-variogram to visualize spatial autocorrelation pattern^55, 56^. The model-based hypothesis test in SPARK also distinguishes it from previous simple spatial test statistics such as Moran’s I and Geary’s C^57, 58^ for detecting spatial autocorrelation patterns. However, presumably because Moran’s I relies on asymptotic normality, its *p*-values under permuted null were highly inflated in the real data we examined here (Fig. S34). In addition, Moran’s I effectively computes correlation among neighboring locations to detect the existence of spatial autocorrelation. Subsequently, these conventional test statistics are not specifically designed to detect spatial patterns other than autocorrelation. For example, studies have shown that Moran’s I (and Geary’s C) are not well powered to detect spatial periodicity patterns^57, 58^. In contrast, by incorporating multiple spatial kernel functions, SPARK can accommodate a range of spatial patterns commonly observed in spatial transcriptomics studies. Indeed, in our analysis, we also found that Moran’s I test was unable to identify most SE genes in pattern I from the mouse olfactory bulb data, including the well-known genes *Doc2g* and *Reln*, whose spatial patterns do not reflect simple autocorrelation. Subsequently, the power of Moran’s I test was lower than SPARK across all four real data sets (Fig. S35).

We have primarily focused on modeling count data with SPARK. Modeling count data directly allows us to account for the mean-variance dependency observed in the spatial data (Fig. S36), resulting in an appreciable power gain. Such power gain is especially apparent in data with low count reads such as the first two spatial transcriptomics data we examined here. However, we acknowledge that the power gain brought by count modeling may be small in data with high count reads such as the MERFISH and seqFISH data, since a normal distribution can often approximate high counts as well as an over-dispersed Poisson distribution. Subsequently, it could be beneficial to provide a Gaussian version of SPARK. Here, we have developed such a Gaussian version of SPARK and implemented it in the SPARK software package. Modeling and algorithm details are provided in the Supplementary Text. The Gaussian version of SPARK allows for robust modeling and scalable computation and can be particularly beneficial for data with high counts. Because we rely on novel statistical techniques to obtain p-values, the Gaussian version of SPARK also produces well-calibrated *p*-values in all permuted data (Fig. S37), much more so than the *p*-values from SpatialDE. While the power of the Gaussian version of SPARK is inferior to the Poisson version of SPARK for data with low counts (Figs. S38A-S38B), its power is somewhat comparable with the Poisson version of SPARK for data with high counts, even though its power remains higher than that of SpatialDE (Figs. S38C-S38D). We hope that by providing both the Poisson and Gaussian versions of SPARK, practitioners can make their own choice in selecting the appropriate model for applied data analysis.

We have primarily focused on aggregating *p*-values obtained from ten different kernels. Aggregating *p*-values across different kernels ensures stable performance across a range of possible scenarios. However, we fully acknowledge that some kernels may work preferentially well for certain data sets (Fig. S39), for detecting certain spatial patterns, and/or for identifying certain SE genes. Subsequently, it could be beneficial to estimate the weights of the ten kernels for each gene separately. In addition, because many SE genes may share similar spatial expression pattern, it could be beneficial to exploit such common information across multiple genes to further improve the power of SE analysis. For example, we could infer for all genes in the same gene set a common set of weights, with which to combine *p*-values from different kernels. How to obtain these kernel weights and how to combine *p*-values in a weighted fashion are important topics for future research.

There are several potential extensions for SPARK. We have primarily focused on *de novo* detection of genes with spatial expression patterns without knowing what specific spatial patterns to look for in the data *a priori*. If we have prior knowledge on the structure of the tissue, we can incorporate such structural information into the kernel functions to facilitate the detection of genes that are specifically expressed in the known structures. We have primarily focused on analyzing spatial transcriptomic data collected on a two-dimensional space of a tissue/culture layout. SPARK is flexible and can be easily extended to analyse three-dimensional (3D) data sets such as STARmap where the depth of the sample location in the tissue can be recorded^59^ or even higher dimensional data sets where other coordinates (e.g. time) are also recorded. We have primarily relied on simple spatial pattern plots and hierarchical clustering for downstream analysis to visualize and categorize identified SE genes from SPARK. Using model based downstream analysis approaches such as the hidden Markov random field model^60^ may provide additional accuracy in the categorization of spatial patterns inferred from identified SE genes. We have primarily focused on using an over-dispersed Poisson model to model count data. Several recent studies have shown that over-dispersed Poisson models are well suited for modeling the data generating process underlying, for example, the unique molecular identifier (UMI) based sequencing studies^61, 62^. However, exploring the use of zero inflated models for data types with inflated zero counts could be a useful future extension. We have primarily focused on analyzing one gene at a time. Future extension of SPARK towards joint modelling of multiple correlated genes in a hierarchical Bayesian framework may further increase power, as it allows for information sharing on the common spatial patterns inferred across genes. Finally, SPARK is computationally efficient. It takes less than an hour to analyze each real data set examined here (Table S3) and can easily handle tens of thousands of genes measured on tens of thousands of spatial locations (Fig. S40). However, extending SPARK to analyze the larger data collected from emerging techniques such as Slide-seq^63^ will likely require new algorithmic development or better computing environment other than standard desktop PCs.

## MATERIALS AND METHODS

### SPARK: Model and Algorithm

We consider modeling gene expression data collected by various high-throughput spatial sequencing techniques such as smFISH and spatial transcriptomics technology. These spatial techniques simultaneously measure gene expression levels of *m* different genes on *n* different spatial locations on a tissue of interest (which we simply refer to as samples). The gene expression measurements are often obtained in the form of counts: they are collected either as the number of barcoded mRNA for any given transcript in a single cell through smFISH based techniques or as the number of sequencing reads mapped to any given gene through sequencing based spatial techniques. The ru, varies across different spatial sequencing techniques and often ranges from a couple hundred (in the case of smFISH) to the whole transcriptome (in the case of spatial transcriptomics technology). The sample composition varies across different spatial sequencing techniques and can consist of either a single cell (in the case of smFISH) or a small set of approximately homogenous single cells residing in a small region of the sampled location known as a spot (in the case of spatial transcriptomics technology). The sampled locations have known spatial coordinates that are recorded during the experiment. These sampled locations can either be considered as random (in the case of smFISH; as expressions are measured on single cells that are randomly scattered across the tissue/culture space) or are pre-determined by researches (in the case of spatial transcriptomics technology) before the experiment. We denote ***s****_i_* = (*s*_*i*1_, *s*_*i*2_) as the spatial coordinates (i.e. location index) for *i*’th sample, with *i* ∈ (1, …, *n*). These spatial coordinates vary continuously over a two-dimensional space *R*^2^, or ***s****_i_* ∈ *R*^2^. While we only focus on the cases where samples are collected on a two-dimensional space of a tissue/culture layout, our model and method are general, capable of handling three-dimensional cases where the depth of the sample location in the tissue can be recorded or handling even higher dimensional cases where other coordinates (e.g. time) are also recorded.

Our primary goal is to detect genes whose expression level displays spatial pattern with respect to the sample locations. We simply refer to these genes as SE genes (genes with spatial expression pattern), in parallel to DE genes (differentially expressed genes) used in other settings. To identify SE genes, we examine one gene at a time and model its expression level across sampled locations using a generalized linear spatial model (GLSM)^64, 65^. GLSM, also known as the generalized linear geostatistical model or the spatial generalized linear mixed model, is a generalized linear mixed model that directly models non-Gaussian spatial data and uses random effects to capture the underlying stationary spatial process. GSLM has been commonly used for interpolating and prediction of spatial data, with applications in spatial disease mapping and spatial epidemiologic studies^66, 67^. However, different from all these previous GLSM development, we instead focus on developing a hypothesis testing framework for GLSM. Here, for the gene of focus, we denote *y_i_*(***s****_i_*) as the gene expression measurement in terms of counts for the *i*’th sample. We denote ***x****_i_*(***s****_i_*) as a *k*-vector of covariates that include a scalar of one for the intercept and *k*-1 observed explanatory variables for the *i*’th sample. These explanatory variables could contain batch information, cell cycle information, or other information that are important to adjust for during the analysis. We denote *N_i_*(***s****_i_*) as the normalization factor for *i*’th sample. Here, we set *N_i_*(***s****_i_*) as the summation of the total number of counts across all genes for the sample as our main interest is in analyzing the relative gene expression level. Other choices of *N_i_*(***s****_i_*) are possible; for example, *N_i_*(***s****_i_*) can be set to one if the main interest is in the absolute gene expression level. We consider modeling the observed expression count data with an over-dispersed Poisson distribution

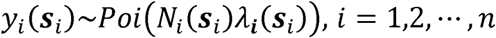

where *λ_i_*(***s****_i_*) is an unknown Poisson rate parameter that represents the underlying gene expression level for the *i*’th sample. In the spatial setting, *λ_i_*(***s****_i_*) can also be viewed as the unobserved spatial random process occurred at location ***s****_i_*. We model the log scale of the latent variable λ*_i_*(***s****_i_*) as a linear combination of three terms,

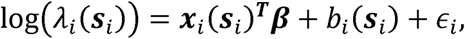

where ***β*** is a *k*-vector of coefficients that include an intercept representing the mean log-expression of the gene across spatial locations together with *k*-1 coefficients for the corresponding explanatory variables; *∊_i_* is the residual error that is independently and identically distributed from *N*(0, *τ*_2_) with variance *τ*_2_; and *b_i_*(***s****_i_*) is a zero-mean, stationary Gaussian process modeling the spatial correlation pattern among spatial locations

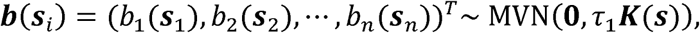

where the covariance ***K***(***s***) is a kernel function of the spatial locations ***s*** = (***s***_1_, …, ***s***_2_)^*T*^, with *ij*’th element being ***K***(***s**_i_, **s**_j_*); *τ*_1_ is a scaling factor of the covariance kernel; and MVN denotes a multivariate normal distribution. We will discuss the choice of the kernel function in more details below. In the above model, the covariance for the latent variables log(*λ_i_*(***s****_i_*)) is **Σ** = *τ*_1_***K***(***s***) + *τ*_2_***I***, where ***λ***(***s****_i_*) = (*λ*_1_(***s***_1_), *λ*_2_(***s***_2_), …, *λ_n_*(***s****_n_*))*^T^* and ***I*** is an *n*-dimensional identity matrix. In spatial statistics, *τ*_1_ is commonly referred to as the partial sill which effectively measures the expression variance in log(*λ_i_*(***s****_i_*)) captured by spatial patterns or spatial location information; *τ*_2_ is commonly referred to as the nugget which effectively measures the expression variance in log(*λ_i_*(***s****_i_*)) due to random noise independent of spatial locations.

In the GLSM defined above, testing whether a gene shows spatial expression pattern can be translated into testing the null hypothesis ***H*_0_**: *τ*_1_=0. The statistical power of such hypothesis test will inevitably depend on how the spatial kernel function ***K***(***s***) matches the true underlying spatial pattern displayed by the gene of interest. For example, a periodic kernel will be particularly useful to detect expression pattern that is periodic across the location space, while a Gaussian kernel will be particularly useful to detect expression pattern that is clustered in focal areas. The true underlying spatial pattern for any gene is unfortunately unknown and may vary across genes. To ensure robust identification of SE genes across various spatial patterns, we consider using a total of ten different spatial kernels, including five periodic kernels with different periodicity parameters and five Gaussian kernels with different smoothness parameters. The detailed construction of these kernels is described in Supplementary Text. These ten kernels cover a range of possible spatial patterns that are observed in common biological data sets (Fig. S41) and are used as default kernels in our software implementation for all analysis results presented here. However, we note that our method and software implemented can easily handle many other kernel functions or incorporate different number of kernel functions as the users see fit.

We fit the above GLSM and test the null hypothesis using the ten kernels one at a time. Parameter estimation and hypothesis testing in GLSM is notoriously difficult, as the GLSM likelihood consists of an *n*-dimensional integral that cannot be solved analytically. To overcome the high dimensional integral and enable scalable estimation and inference with GLSM, we develop an approximate inference algorithm based on the penalized quasi-likelihood (PQL) approach^20, 68^. The algorithmic details are provided in the Supplementary Text. With parameter estimates from the PQL-based algorithm, we computed a *p*-value for each of the ten kernels using the Satterthwaite Method^69^ based on score statistics, which follow a mixture of chi-square distributions. Afterwards, we combined these ten *p*-values through the recently developed Cauchy *p*-value combination rule^21^. To apply the Cauchy combination rule, we converted each of the ten *p*-values into a Cauchy statistic, aggregated the ten Cauchy statistics through summation, and converted the summation back to a single *p*-value based on the standard Cauchy distribution. The Cauchy rule takes advantage of the fact that combination of Cauchy random variables also follows a Cauchy distribution regardless whether these random variables are correlated or not^21, 22^. Therefore, the Cauchy combination rule allows us to combine multiple potentially correlated *p*-values into a single *p*-value without loss of type I error control. After obtaining *m p*-values across *m* genes, we controlled for false discovery rate (FDR) using the *Benjamini–Yekutieli* (BY) procedure, which is effective under arbitrary dependence across genes^70^.

We refer to the above method as the Poisson version of SPARK (Spatial PAttern Recognition via Kernels) and is the main method used in the present study. Besides the Poisson version, we have also developed a Gaussian version of SPARK for modeling normalized spatial data (Supplementary Text). Both versions of SPARK are implemented in the same R package with multiple threads computing capability, and with underlying efficient C/C++ code linked through Rcpp. The software SPARK, together with all analysis code used in the present study for reproducing the results presented in the manuscript, are freely available at www.xzlab.org/software.html.

### Simulation Designs

We performed two sets of simulations. In the first set of simulations, we simulated gene expression data on 260 spatial locations (i.e. spots) collected in the mouse olfactory bulb study using parameters inferred from the corresponding real data (detail of the study is described in the next section). For null simulations, we simulated 10,000 non-SE genes on these locations to examine type I error control. For power simulations, we simulated 1,000 SE genes and 9,000 non-SE genes on these locations to examine method power. In either null or power simulations, for each gene in turn, we first simulated the non-spatial residual errors on the spots independently based on a normal distribution with mean zero and variance being either 0.2, 0.35 or 0.6, which are equivalent to approximately the first quartile, median and third quartile of the non-spatial variance estimates in the real data, respectively. For non-SE genes, we set the intercept to be −10.2, which is shared across all spots and corresponds to the median of the intercept estimates in the mouse olfactory data. For SE genes, we first categorized spots into two groups – a group of spots with low expression levels and a group of spots with high expression levels -- based on the three spatial patterns illustrated in Fig. 1C. We then set the intercept in the low expression group to be −10.2 and set the intercept in the high expression group to be either two-fold, three-fold or four-fold higher than the lower one on rate parameter scale; thus the intercept in the high expression group is −9.5, −9.1, and −8.8, for the three cases, respectively. The difference in the intercept between the two groups of spots thus introduces spatial differential expression pattern. Finally, for each gene in turn, regardless whether it is SE or non-SE, we summed the residual errors and the intercept to a spot-specific latent variable log*λ_i_*. We then simulated the gene expression count data based on a Poisson distribution with the rate being a product of the latent variable *λ_i_* and the total read counts (*N_i_*) that is obtained from the real data. That is, *y_i_*∼*Poi*(*N_i_λ_i_*) for the *i*’th spot. With the above procedure, in each of the spatial pattern illustrated in Fig. 1C, we first simulated data using a baseline parameter setting where the noise variance is set to be 0.35 and the intercept is set to be either −10.2 for non-SE genes or −10.2/−9.1 (representing a three-fold change) for SE genes. Afterwards, we varied one parameter at a time to examine the influence of different parameters on the performance of different methods. We performed 10 replicates for each scenario and combined results across all 10 replicates.

In the second set of simulations, we simulated count data on spatially distributed cells following the Trendsceek paper. Specifically, we first randomly simulated the spatial locations for a fixed number of cells (*n* = 100, 200 or 500) through a random-point-pattern Poisson process. We generated 1,000 genes in the simulated data, which were all non-SE genes in the null simulations and consisted of 100 SE genes and 900 non-SE genes in the power simulations. For non-SE genes, the expression measurements from the real data were randomly assigned to the simulated cells regardless of their spatial locations (Fig. 1E). For SE genes, the expression measurements from the real data were assigned to the simulated cells to display three distinct spatial patterns (Hotspot, Streak and Gradient patterns; Fig. 1E). Specifically, for the first two spatial patterns, we created either a circle (for Hotspot pattern) or a band (for Streak pattern) in the middle of the panel and marked cells residing in these areas. The size of the circle and the size of the band were designed so that the marked cells inside these areas represent a fixed proportion of all cells, with the proportion set to be either 10%, 20%, or 30%. The expression measurements of the non-marked cells were randomly assigned from the observed expression distribution of the gene in the seqFISH data. The expression measurements of the marked cells were randomly assigned from either the upper quantile (for 50 SE genes) or the lower quantile (for the other 50 SE genes) of the expression distribution of the gene in the seqFISH data. We set the quantile cutoff to be either 40%, 66% or 80%, representing low, moderate, or high SE signal strength, respectively. These three different quantile cutoffs correspond to an expected expression fold change between the marked cells and the non-marked cells of either 1.5, 2 or 2.5, respectively. For the Gradient pattern, the expression levels of a fraction of marked cells (=30%, 40%, or 50%) were set either in an increasing order (for 50 SE genes) or a decreasing order (for the other 50 SE genes) along the x-axis. To do so, we draw the expression measurements for the marked cells randomly from the observed expression distribution in the real data and assigned these drawn values in either increasing or decreasing order to the marked cells based on their x-axis coordinates. In contrast, the expression measurements of the non-marked cells were again randomly assigned from the observed expression distribution of the gene in the seqFISH data, regardless of their spatial location. In all these simulations, we varied the number of cells (*n* = 100, 200 or 500), the SE strength (low, moderate or high; measured by the quantile cutoff for the first two spatial patterns and by the fraction of cells displaying expression gradient for the third spatial pattern), as well as the fraction of cells in the focal/streak area for the first two spatial patterns.

### Clustering SE Genes Detected by SPARK

We summarized the spatial expression patterns detected by SPARK by dividing SE genes into different categories. To do so, we first applied variance-stabilizing transformation (VST) to the raw count data^12^ and obtained the relative gene expression levels through adjusting for the log-scale total read counts. We then used the hierarchical agglomerative clustering algorithm in the R package *amap* (v0.8-17) to cluster identified SE genes detected by SPARK into five groups. Afterwards, we summarized the gene expression patterns by using the expression level of the five cluster centers (Fig. S42). In the hierarchical clustering, we set the two optional parameters in the R function to be Euclidean distance and Ward’s criterion, respectively.

### Gene Sets and Functional Enrichment Analysis

For each of the first two real data sets, we obtained lists of genes that can be used to serve as unbiased validation for the SE genes identified by different methods. Specifically, for the olfactory bulb data, we obtained a gene list directly based on the three layers (mitral, glomerular and granule) of the main olfactory bulb listed in the Harmonizome database (https://amp.pharm.mssm.edu/Harmonizome/). For the breast cancer data, we obtained from the Harmonizome database a gene list that consists of breast cancer related genes from six different data sets (OMIM Gene-Disease Associations; PhosphoSitePlus Phosphosite-Disease Associations; DISEASES Text-mining Gene-Disease Association Evidence Scores; GAD Gene-Disease Associations; GWAS Catalog SNP-Phenotype Associations). For the breast cancer data, we also obtained from the CancerMine database (http://bionlp.bcgsc.ca/cancermine/) another gene list that consists of breast cancer related genes that are either cancer drivers, oncogenes, or tumor suppressors. We used these gene lists to validate the SE genes identified by different methods.

In addition, we performed the functional enrichment analysis of significant SE genes identified by SPARK and SpatialDE in Gene Ontology (GO) and Kyoto Encyclopedia of Genes and Genomes (KEGG). We performed all enrichment analyses using the R package *clusterProfiler*^71^ (v3.12.0). In the package, we used the default “BH” method for *p*-value multiple testing correction and we used the default number of permutations to be 1,000.

### Spatial Transcriptomics Data Sets

We downloaded two spatial transcriptomics data sets from the Spatial Transcriptomics Research (http://www.spatialtranscriptomicsresearch.org). These two data sets include a mouse olfactory bulb data and a human breast cancer data. These data consist of gene expression measurements in the form of read counts that are collected on a number of spatial locations known as spots. Following the SpatialDE paper, we used the *MOB Replicate 11* file for mouse olfactory bulb data, which contains 16,218 genes measured on 262 spots and *Breast Cancer Layer 2* file for the breast cancer data, which contains 14,789 genes measured on 251 spots. We filtered out genes that are expressed in less than 10% of the array spots and selected spots with at least 10 total read counts. With these filtering criteria, we analyzed a final set of 11,274 genes on 260 spots in the mouse olfactory bulb data and 5,262 genes on 250 spots for the breast cancer data. In the analysis, we performed permutations to construct an empirical null distribution of *p*-values for each method by permuting the spot coordinates 10 times. Afterwards, we examined type I error control of different methods based on the empirical null distribution of *p*-values.

### MERFISH Data Set

We obtained the MERFISH data set collected on the mouse preoptic region of the hypothalamus from Dryad^42, 72^. We used the slice at Bregma +0.11mm from animal 18 for analysis, as it contains all 160 genes measured on the largest number of single cells (5,665) across all nine cell classes. Among the 160 genes, 155 of them were pre-selected in the original study as either known markers for major cell classes or relevant to various neuronal functions of the hypothalamus (e.g. some are neuropeptides and some are neuro-modulator receptors). Most of these 155 genes are expected to have spatial expression pattern in the hypothalamus. The remaining 5 genes are blank control genes without spatial expression pattern in the hypothalamus and thus can serve as negative controls. The downloaded data contains normalized genes expression values, which were computed as read counts divided by either the cell volume (combinatorial smFISH) or arbitrary fluorescence units per µm^3^ (non-combinatorial, sequential FISH) and further scaled by 1,000. To obtain the raw count data, we thus rescaled the expression values by first multiplying 1,000, adjusted for cell volume, and then converted the rescaled value into integers by taking the ceiling over the rescaled data. After removing the ambiguous cells that were identified as putative doublets in the original data, we analyzed a final set of 160 genes on 4,975 cells. In the analysis, we permuted the location coordinates 100 times to construct an empirical null distribution, with which we examined type I error control of different methods.

### SeqFISH Data Set

We obtained the seqFISH data set collected on the mouse hippocampus from the supplementary file of the original paper^48^. Following the SpatialDE paper, we extracted the field 43 data set for analysis. The data are in the form of raw count data for 249 genes measured in 257 cells with known spatial location information. Among 249 measured genes, 214 were selected from a list of transcription factors and signaling pathway components, and the remaining 35 were selected from cell identity markers^48^. Following Trendsceek^13^ and the original study^48^, we filtered out cells with x- or y-axis values falling outside the range of 203 - 822 pixels in order to address border artifacts. After filtering, we analyzed a final set of 249 genes measured on 131 cells. In the analysis, we permuted the location coordinates 100 times to construct an empirical null distribution, with which we examined type I error control of different methods.

### Compared Methods

We compared SPARK with two existing methods for detecting genes with spatial expression patterns. Both of these methods are designed for normalized data. The first method is Trendsceek (R package *trendsceek*; v1.0.0; download date: 12/20/2018). We followed the same procedure described in the original Trendsceek paper^13^ to filter and normalize count data. Specifically, for the two spatial transcriptomics data, we excluded genes that are expressed in less than 3 spots and excluded spots that contain less than 5 read counts. We then performed log10-transformation on raw count data (by adding a pseudo-count of one to void log transformation of zero values). For the real data analysis, we focused on analyzing the top 500 most variable genes to ensure sufficient power as well as computational feasibility as described in the Trendsceek paper. For the permuted data, we analyzed all the genes to construct an empirical null distribution. For seqFISH data, we first removed boundary cells as described in the previous section. Afterwards, following the Trendsceek recommendation, for each gene in turn, we performed a one-sided winsorization procedure to remove outlier effects by setting the first four largest values to be the fifth largest value. We then applied log10-transformation on the count data (again adding a pseudo-count of one) to obtain normalized expression values. For MERFISH data, we performed log10-transformation on raw count data (again adding a pseudo-count of one) and included all genes for analysis. Besides filtering and normalization, Trendsceek relies on permutation to compute *p*-values. Here, we set the number of permutations to be the default of 10,000. In addition, because the results of Trendsceek depend on the seeds used in the software, we analyzed each data using ten different seeds and reported results based on the seed that yields the highest number of discoveries; thus the power estimates of Trendsceek are likely upward biased. One disadvantage of Trendsceek is its slow computation: it takes over 48 hours to analyze one single gene in the mouse hypothalamus data. Therefore, in that data, we only applied the Trendsceek to the real data but not to the permuted data. Following the Trendsceek paper, we used the *Benjamini-Hochberg* procedure implemented in Trendsceek software to obtain adjusted *p-* value (i.e. FDR). With the adjusted *p*-value, we declared an SE gene significant if at least one of the four adjusted *p*-value outputs from (the four tests of) Trendsceek is below the threshold of 0.05.

The second method we compared with is SpatialDE (python package; v.1.1.0; download date: 12/12/2018). For the mouse olfactory data and human breast cancer data, we directly used the analysis code provided by the SpatialDE authors on the Github (https://github.com/Teichlab/SpatialDE) to perform analysis.

For the mouse hippocampus data, we applied their analysis code to the border artifacts adjusted data set described above to avoid detection of border artifacts and ensure fair comparison across methods. For the mouse hypothalamus data, we also directly applied the MERFISH analysis code described in the SpatialDE paper. Following the SpatialDE paper, we declared an SE gene as significant if the output *q*-value (i.e. FDR) from SpatialDE is below the threshold of 0.05.

Finally, we also examined the performance of Moran’s I test in all four real data sets. We used the function *moran.test* implemented in the R package *spdep* (v1.1.2) for this analysis. The results on Moran’s I are presented in the Discussion.

## ACKNOWLEDGEMENTS

This study was supported by the National Institutes of Health (NIH) Grants R01HG009124 and R01GM126553, and the National Science Foundation (NSF) Grant DMS1712933. S.S. was supported by NIH Grant R01HD088558 (PI Tung), the National Natural Science Foundation of China (Grant No. 61902319), and the Natural Science Foundation of Shaanxi Province (Grant No. 2019JQ127). J.Z. was supported by NIH Grant U01HL137182 (PI Kang).

## AUTHORS’ CONTRIBUTIONS

XZ conceived the idea and provided funding support. SS and XZ designed the experiments. SS and JQ developed the method, implemented the software, performed simulations and analyzed real data. SS, JQ and XZ wrote the manuscript.

## AVAILABILITY OF DATA AND MATERIALS

All source code and data sets used in our experiments have been deposited at www.xzlab.org/software.html.

## CONSENT FOR PUBLICATION

Not applicable.

## REFERENCE

1. Chen, K.H., Boettiger, A.N., Moffitt, J.R., Wang, S. & Zhuang, X. RNA imaging. Spatially resolved, highly multiplexed RNA profiling in single cells. Science 348, aaa6090 (2015).

2. Lubeck, E., Coskun, A.F., Zhiyentayev, T., Ahmad, M. & Cai, L. Single-cell in situ RNA profiling by sequential hybridization. Nat Methods 11, 360–361 (2014).

3. Femino, A.M., Fogarty, K., Lifshitz, L.M., Carrington, W. & Singer, R.H. Visualization of single molecules of mRNA in situ. Method Enzymol 361, 245–304 (2003).

4. Lovatt, D. et al. Transcriptome in vivo analysis (TIVA) of spatially defined single cells in live tissue. Nat Methods 11, 190–196 (2014).

5. Simone, N.L., Bonner, R.F., Gillespie, J.W., Emmert-Buck, M.R. & Liotta, L.A. Laser-capture microdissection: opening the microscopic frontier to molecular analysis. Trends In Genetics 14, 272–276 (1998).

6. Junker, J.P. et al. Genome-wide RNA Tomography in the Zebrafish Embryo. Cell 159, 662–675 (2014).

7. Stahl, P.L. et al. Visualization and analysis of gene expression in tissue sections by spatial transcriptomics. Science 353, 78–82 (2016).

8. Ke, R.Q. et al. In situ sequencing for RNA analysis in preserved tissue and cells. Nat Methods 10, 857–860 (2013).

9. Lee, J.H. et al. Highly Multiplexed Subcellular RNA Sequencing in Situ. Science 343, 1360–1363 (2014).

10. Crosetto, N., Bienko, M. & van Oudenaarden, A. Spatially resolved transcriptomics and beyond. Nat Rev Genet 16, 57–66 (2015).

11. Fan, X. et al. Spatial transcriptomic survey of human embryonic cerebral cortex by single-cell RNA-seq analysis. Cell Res 28, 730–745 (2018).

12. Svensson, V., Teichmann, S.A. & Stegle, O. SpatialDE: identification of spatially variable genes. Nat Methods 15, 343–346 (2018).

13. Edsgard, D., Johnsson, P. & Sandberg, R. Identification of spatial expression trends in single-cell gene expression data. Nat Methods 15, 339–342 (2018).

14. Lun, A. Overcoming systematic errors caused by log-transformation of normalized single-cell RNA sequencing data. BioRxiv, 404962 (2019).

15. Lea, A.J., Alberts, S.C., Tung, J. & Zhou, X. A flexible, efficient binomial mixed model for identifying differential DNA methylation in bisulfite sequencing data. Plos Genet 11, e1005650 (2015).

16. Sun, S.Q. et al. Differential expression analysis for RNAseq using Poisson mixed models. Nucleic Acids Res 45, e106 (2017).

17. Li, Y., Tang, H.C. & Lin, X.H. Spatial Linear Mixed Models with Covariate Measurement Errors. Stat Sinica 19, 1077–1093 (2009).

18. Ben-Ahmed, K., Bouratbine, A. & El-Aroui, M.A. Generalized linear spatial models in epidemiology: A case study of zoonotic cutaneous leishmaniasis in Tunisia. J Appl Stat 37, 159–170 (2010).

19. Breslow, N.E. & Lin, X.H. Bias Correction In Generalized Linear Mixed Models with a Single-Component Of Dispersion. Biometrika 82, 81–91 (1995).

20. Sun, S.Q. et al. Heritability estimation and differential analysis of count data with generalized linear mixed models in genomic sequencing studies. Bioinformatics 35, 487–496 (2019).

21. Liu, Y.W. et al. ACAT: A Fast and Powerful p Value Combination Method for Rare-Variant Analysis in Sequencing Studies. Am J Hum Genet 104, 410–421 (2019).

22. Pillai, N.S. & Meng, X.L. An Unexpected Encounter with Cauchy And Levy. Ann Stat 44, 2089–2097 (2016).

23. Tepe, B. et al. Single-Cell RNA-Seq of Mouse Olfactory Bulb Reveals Cellular Heterogeneity and Activity-Dependent Molecular Census of Adult-Born Neurons. Cell Rep 25, 2689–2703 (2018).

24. Rouillard, A.D. et al. The harmonizome: a collection of processed datasets gathered to serve and mine knowledge about genes and proteins. Database-Oxford 2016, baw100 (2016).

25. Menalled, L.B., Sison, J.D., Dragatsis, I., Zeitlin, S. & Chesselet, M.F. Time course of early motor and neuropathological anomalies in a knock-in mouse model of Huntington’s disease with 140 CAG repeats. J Comp Neurol 465, 11–26 (2003).

26. Adan, R.A.H. et al. Rat Oxytocin Receptor In Brain, Pituitary, Mammary-Gland, And Uterus - Partial Sequence And Immunocytochemical Localization. Endocrinology 136, 4022–4028 (1995).

27. Didier, A. et al. A dendrodendritic reciprocal synapse provides a recurrent excitatory connection in the olfactory bulb. P Natl Acad Sci USA 98, 6441–6446 (2001).

28. Hack, I., Bancila, M., Loulier, K., Carroll, P. & Cremer, H. Reelin is a detachment signal in tangential chain-migration during postnatal neurogenesis. Nat Neurosci 5, 939–945 (2002).

29. Kosaka, T., Deans, M.R., Paul, D.L. & Kosaka, K. Neuronal gap junctions in the mouse main olfactory bulb: Morphological analyses on transgenic mice. Neuroscience 134, 757–769 (2005).

30. Morita, K., Sasaki, H., Furuse, M. & Tsukita, S. Endothelial claudin: Claudin-5/TMVCF constitutes tight junction strands in endothelial cells. J Cell Biol 147, 185–194 (1999).

31. Djurisic, M., Popovic, M., Carnevale, N. & Zecevic, D. Functional structure of the mitral cell dendritic tuft in the rat olfactory bulb. J Neurosci 28, 4057–4068 (2008).

32. Zou, D.J., Greer, C.A. & Firestein, S. Expression pattern of alpha CaMKII in the mouse main olfactory bulb. J Comp Neurol 443, 226–236 (2002).

33. Lever, J., Zhao, E.Y., Grewal, J., Jones, M.R. & Jones, S.J.M. CancerMine: a literature-mined resource for drivers, oncogenes and tumor suppressors in cancer. Nat Methods 16, 505–507 (2019).

34. Esquivel-Velazquez, M. et al. The Role of Cytokines in Breast Cancer Development and Progression. J Interf Cytok Res 35, 1–16 (2015).

35. Beck, S. & Trowsdale, J. The human major histocompatibility complex: Lessons from the DNA sequence. Annu Rev Genom Hum G 1, 117–137 (2000).

36. Rossi, J.F., Ceballos, P. & Lu, Z.Y. Immune precision medicine for cancer: a novel insight based on the efficiency of immune effector cells. Cancer Commun 39, 34 (2019).

37. Lin, C.Y., Beattie, A., Baradaran, B., Dray, E. & Duijf, P.H.G. Contradictory mRNA and protein misexpression of EEF1A1 in ductal breast carcinoma due to cell cycle regulation and cellular stress. Sci Rep-Uk 8, 13904 (2018).

38. Liu, H.Z. et al. Increased expression of elongation factor-1 alpha is significantly correlated with poor prognosis of human prostate cancer. Scand J Urol Nephrol 44, 277–283 (2010).

39. McSherry, E.A., Brennan, K., Hudson, L., Hill, A.D.K. & Hopkins, A.M. Breast cancer cell migration is regulated through junctional adhesion molecule-A-mediated activation of Rap1 GTPase. Breast Cancer Res 13 (2011).

40. Chetty, C. et al. MMP-9 induces CD44 cleavage and CD44 mediated cell migration in glioblastoma xenograft cells. Cell Signal 24, 549–559 (2012).

41. Redondo-Munoz, J. et al. alpha 4 beta 1 integrin and 190-kDa CD44v constitute a cell surface docking complex for gelatinase B/MMP-9 in chronic leukemic but not in normal B cells. Blood 112, 169–178 (2008).

42. Moffitt, J.R. et al. Molecular, spatial, and functional single-cell profiling of the hypothalamic preoptic region. Science 362, eaau5324 (2018).

43. Yao, L.H. et al. Hypothalamic gastrin-releasing peptide receptor mediates an antidepressant-like effect in a mouse model of stress. Am J Transl Res 8, 3097–3105 (2016).

44. Fabio, K. et al. Synthesis and evaluation of potent and selective human V1a receptor antagonists as potential ligands for PET or SPECT imaging. Bioorgan Med Chem 20, 1337–1345 (2012).

45. Tabak, B.A. et al. Null results of oxytocin and vasopressin administration across a range of social cognitive and behavioral paradigms: Evidence from a randomized controlled trial. Psychoneuroendocrinology 107, 124–132 (2019).

46. Ozturk, A., DeKosky, S.T. & Kamboh, M.I. Genetic variation in the choline acetyltransferase (CHAT) gene may be associated with the risk of Alzheimer’s disease. Neurobiol Aging 27, 1440–1444 (2006).

47. Kiaris, H., Schally, A.V. & Kalofoutis, A. Extrapituitary effects of the growth hormone-releasing hormone. Vitam Horm 70, 1–24 (2005).

48. Shah, S., Lubeck, E., Zhou, W. & Cai, L. In Situ Transcription Profiling of Single Cells Reveals Spatial Organization of Cells in the Mouse Hippocampus. Neuron 92, 342–357 (2016).

49. Tasic, B. et al. Adult mouse cortical cell taxonomy revealed by single cell transcriptomics. Nat Neurosci 19, 335–346 (2016).

50. Li, X.H., Polter, A. & Yang, S. FoxO transcription factors - Regulation in brain and behavioral manifestation. Biol Psychiat 63, 150–159 (2008).

51. Hoekman, M.F.M., Jacobs, F.M.J., Smidt, M.P. & Burbach, J.P.H. Spatial and temporal expression of FoxO transcription factors in the developing and adult murine brain. Gene Expr Patterns 6, 134–140 (2006).

52. Cattaneo, A. et al. FoxO1, A2M, and TGF-beta 1: three novel genes predicting depression in gene X environment interactions are identified using cross-species and cross-tissues transcriptomic and miRNomic analyses. Mol Psychiatr 23, 2192–2208 (2018).

53. Shrestha, B.R. et al. Sensory Neuron Diversity in the Inner Ear Is Shaped by Activity. Cell 174, 1229–1246 (2018).

54. Sun, Y.F. et al. A central role for Islet1 in sensory neuron development linking sensory and spinal gene regulatory programs. Nat Neurosci 11, 1283–1293 (2008).

55. Voss, S., Zimmermann, B. & Zimmermann, A. Detecting spatial structures in throughfall data: The effect of extent, sample size, sampling design, and variogram estimation method. J Hydrol 540, 527–537 (2016).

56. Lark, R.M., Heuvelink, G.B.M. & Bishop, T.F.A. Burgess, TM & Webster, R. 1980. Optimal interpolation and isarithmic mapping of soil properties. I. The semi-variogram and punctual kriging. Journal of Soil Science, 31, 315-331. Eur J Soil Sci 70, 7–10 (2019).

57. Li, H.F., Calder, C.A. & Cressie, N. Beyond Moran’s I: Testing for spatial dependence based on the spatial autoregressive model. Geogr Anal 39, 357–375 (2007).

58. Radeloff, V.C., Miller, T.F., He, H.S. & Mladenoff, D.J. Periodicity in spatial data and geostatistical models: autocorrelation between patches. Ecography 23, 81–91 (2000).

59. Wang, X. et al. Three-dimensional intact-tissue sequencing of single-cell transcriptional states. Science 361, eaat5691 (2018).

60. Zhu, Q., Shah, S., Dries, R., Cai, L. & Yuan, G.C. Identification of spatially associated subpopulations by combining scRNAseq and sequential fluorescence in situ hybridization data. Nat Biotechnol 36, 1183–1190 (2018).

61. Islam, S. et al. Quantitative single-cell RNA-seq with unique molecular identifiers. Nat Methods 11, 163–166 (2014).

62. Kivioja, T. et al. Counting absolute numbers of molecules using unique molecular identifiers. Nat Methods 9, 72–74 (2012).

63. Rodriques, S.G. et al. Slide-seq: A scalable technology for measuring genome-wide expression at high spatial resolution. Science 363, 1463–1467 (2019).

64. Diggle, P.J., Tawn, J.A. & Moyeed, R.A. Model-based geostatistics. J R Stat Soc C-Appl 47, 299–326 (1998).

65. Christensen, O.F. & Waagepetersen, R. Bayesian prediction of spatial count data using generalized linear mixed models. Biometrics 58, 280–286 (2002).

66. Rousset, F. & Ferdy, J.B. Testing environmental and genetic effects in the presence of spatial autocorrelation. Ecography 37, 781–790 (2014).

67. Vanhatalo, J., Pietilainen, V. & Vehtari, A. Approximate inference for disease mapping with sparse Gaussian processes. Statistics in medicine 29, 1580–1607 (2010).

68. Lin, X.H. & Breslow, N.E. Bias correction in generalized linear mixed models with multiple components of dispersion. J Am Stat Assoc 91, 1007–1016 (1996).

69. Satterthwaite, F.E. An Approximate Distribution Of Estimates Of Variance Components. Biometrics Bull 2, 110–114 (1946).

70. Benjamini, Y. & Yekutieli, D. The control of the false discovery rate in multiple testing under dependency. Ann Stat 29, 1165–1188 (2001).

71. Yu, G.C., Wang, L.G., Han, Y.Y. & He, Q.Y. clusterProfiler: an R Package for Comparing Biological Themes Among Gene Clusters. Omics 16, 284–287 (2012).

72. Moffitt, J.R., et al. in Dryad Digital Repository, Vol. 362. (ed. D.D. Repository) eaau5324 (2018).

