## supplemental text for "Statistical Analysis of Spatial Expression Pattern for Spatially Resolved Transcriptomic Studies"

**Supplementary Figure 1: Quantile-quantile plot of the observed -log10 *p*-values from different methods against the expected -log10 *p*-values in null simulations with different noise levels.** *p*-values are combined across ten simulation replicates. Simulations are performed under different nugget values that represent different noise levels: (**A**) $\tau_{2}$=0.2, (**B**) $\tau_{2}$=0.35, and (**C**)$\tau_{2}$=0.6. Compared methods include SPARK (pink), SpatialDE (purple), Trendsceek.E (light salmon) which is the Emark test of Trendsceek, Trendsceek.$\rho$ (yellow-green) which is the Markcorr test of Trendsceek, Trendsceek.$\gamma$ (light green) which is the Markvario test of Trendsceek, and Trendsceek.V (wheat) which is the Vmark test of Trendsceek. *p*-values from SPARK and some of the Trendsceek methods (e.g. Markvario and Vmark) are well calibrated. In contrast, *p*-values from SpatialDE, and to a lesser extent from the Emark and Markcorr tests of Trendsceek, are overly conservative and distributed below the expected diagonal line.

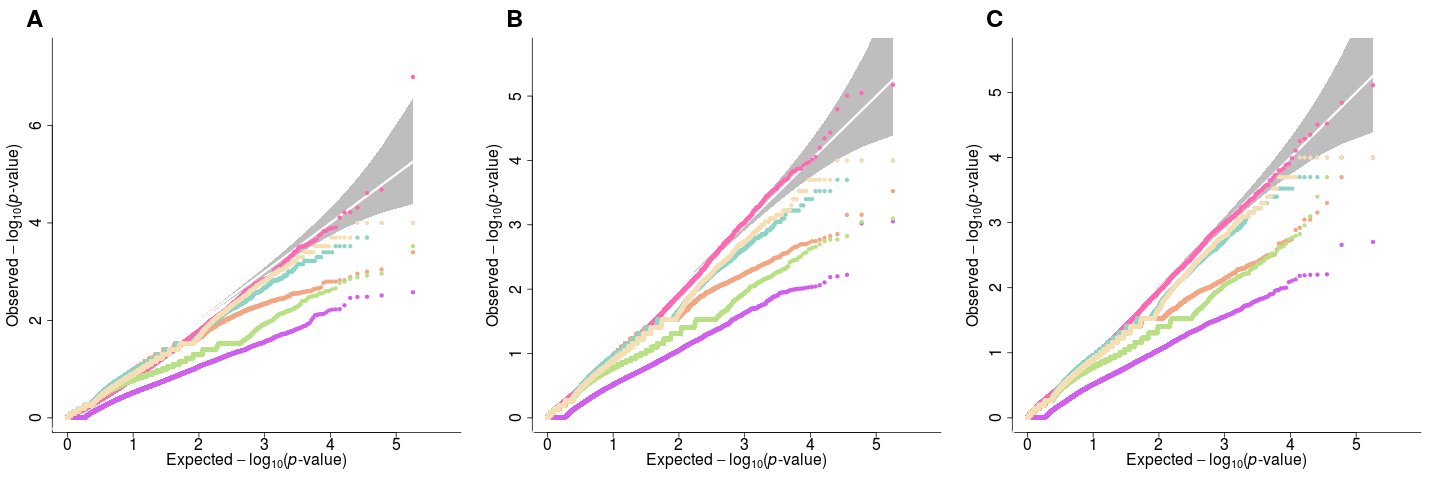

**A**

**B**

**C**

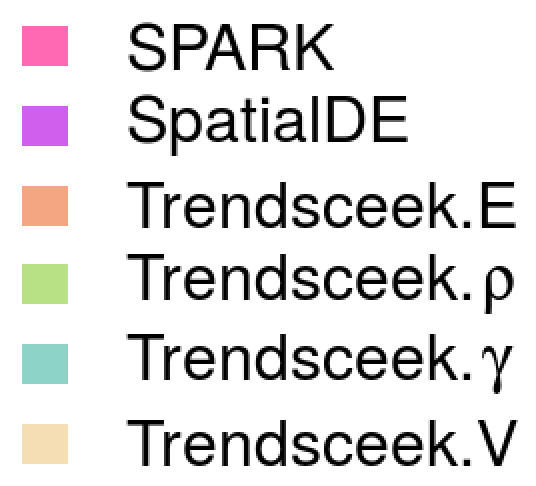

**Supplementary Figure 2: Power comparison of different methods in alternative simulations under different pattern strength.** Power plots show the proportion of true positives (y-axis) detected by different methods at a range of false discovery rates (FDR; x-axis) in the alternative simulations. The proportion of true positives is averaged across ten simulation replicates. Compared methods include SPARK (pink), SpatialDE (purple), Trendsceek (sky-blue) which is combined test of Trendsceek. Simulations are performed under different spatial expression pattern strength: weak pattern (two-fold) in (**A**-**C**); moderate pattern (three-fold) in (**D**-**F**); and strong pattern (four-fold) in (**G**-**I**). Simulations are also performed under three different spatial expression patterns I-III as illustrated in the main Figure 1C: pattern I in (**A**, **D**, and **G**); pattern II in (**B**, **E**, and **H**); and pattern III in (**C**, **F**, and **I**). The nugget level is set to be $\tau_{2}$ = 0.35 in all settings. Across simulations and across FDR cutoffs, SPARK is more powerful than the other two methods for detecting genes with spatial expression patterns.

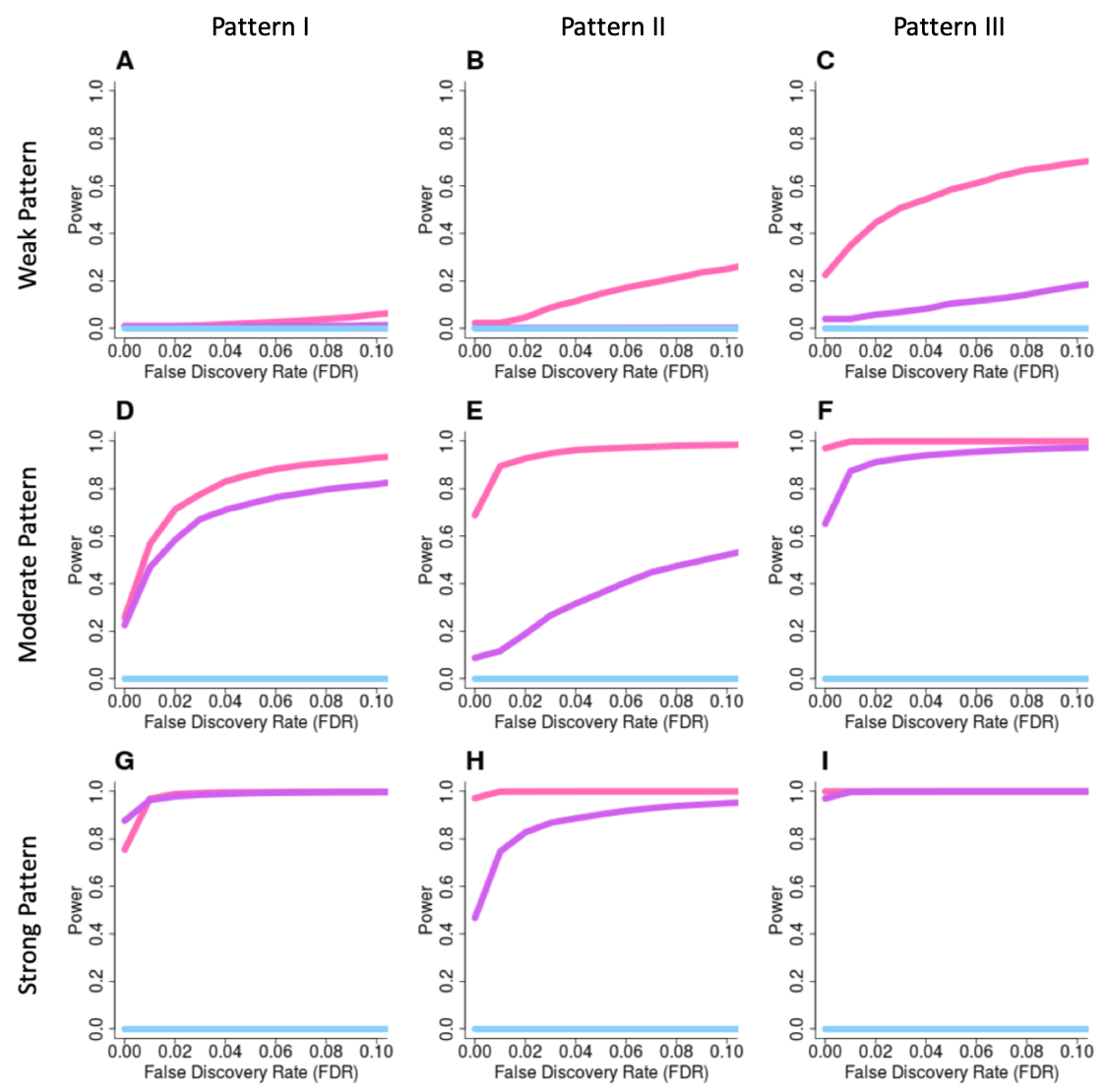

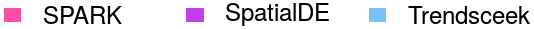

**Supplementary Figure 3: Power comparison of different methods in alternative simulations under different noise levels.** Power plots show the proportion of true positives (y-axis) detected by different methods at a range of false discovery rates (FDR; x-axis) in the alternative simulations. The proportion of true positives is averaged across ten simulation replicates. Compared methods include SPARK (pink), SpatialDE (purple), Trendsceek (sky-blue) which is combined test of Trendsceek. Simulations are performed under different noise levels characterized by different nugget values: low noise ($\tau_{2}$=0.2) in (**A**-**C**); moderate noise ($\tau_{2}$=0.35) in (**D**-**F**); and high noise ($\tau_{2}$=0.6) in (**G**-**I**). Simulations are also performed under three different spatial expression patterns I-III as illustrated in the main Figure 1C: pattern I in (**A**, **D**, and **G**); pattern II in (**B**, **E**, and **H**); and pattern III in (**C**, **F**, and **I**). The spatial expression pattern strength is set to be moderate (three-fold) in all settings. Across simulations and across FDR cutoffs, SPARK is more powerful than the other two methods for detecting genes with spatial expression patterns.

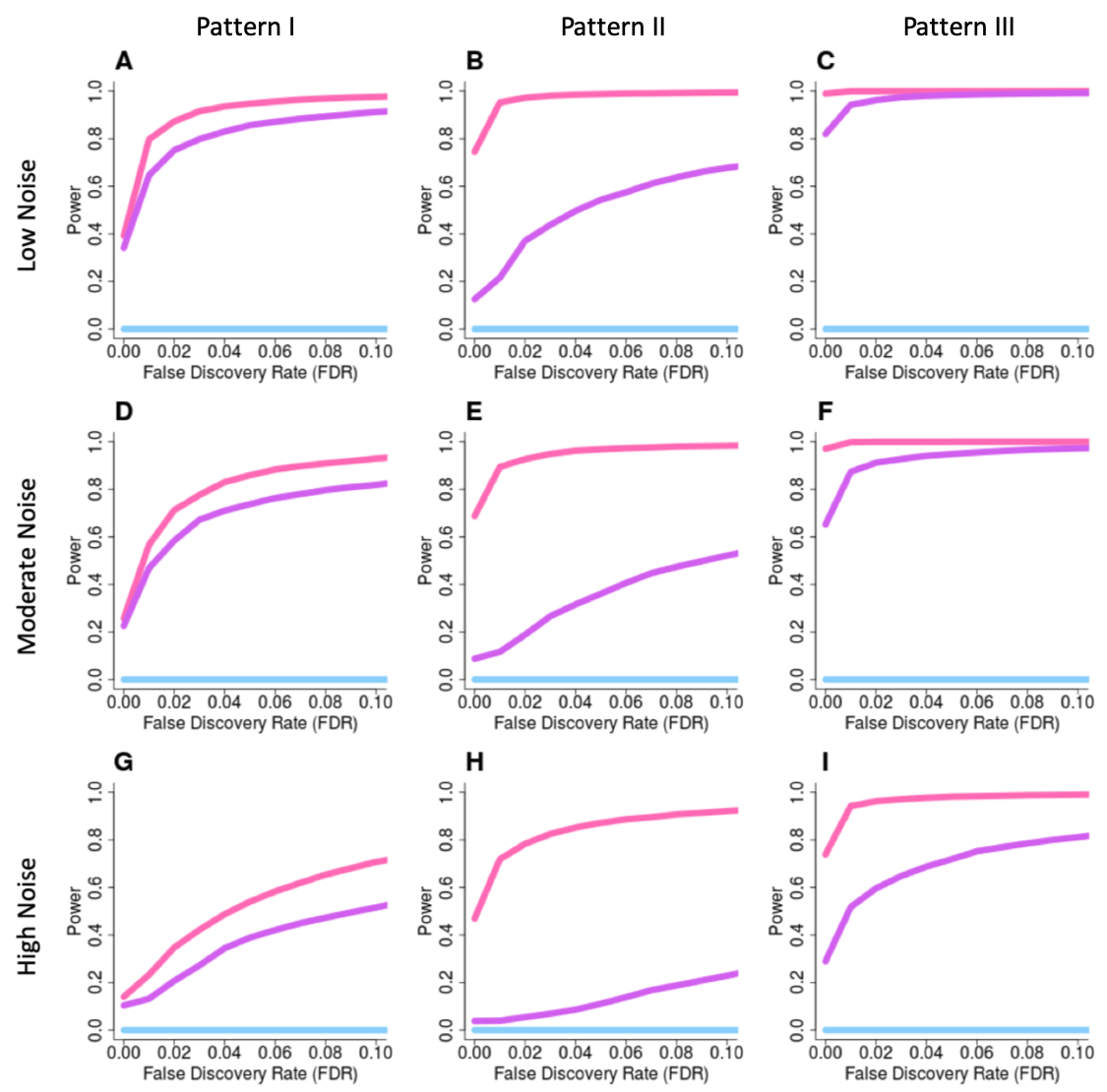

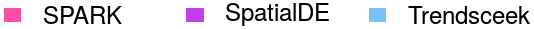

**Supplementary Figure 4: Power comparison of different methods in the alternative simulations under different sample sizes in the simulations based on the original Trendsceek paper.** Power plots show the proportion of true positives (y-axis) detected by different methods at a range of false discovery rates (FDR; x-axis) in the alternative simulations. The proportion of true positives is averaged across ten simulation replicates. Compared methods include SPARK (pink), SpatialDE (purple), Trendsceek (sky-blue) which is combined test of Trendsceek. Simulations are performed under different sample sizes: 100 in (**A** and **D**); 200 in (**B** and **E**); and 500 (**C** and **F**). Simulations were performed under moderate fraction of marked cells (20%) and moderate SE strength (2 fold) for the hotspot (**C**) and non-radial streak (**F**) patterns. When sample size is small or moderate, SPARK is more powerful than the other two methods for detecting genes with spatial expression patterns (first two columns). In the case where sample size equals to 500, both SPARK and SpatialDE reach 100% power while the power of Trendsceek remains low (last column). SS: sample size.

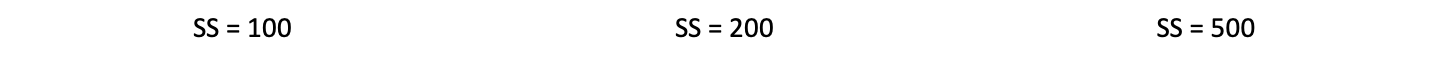

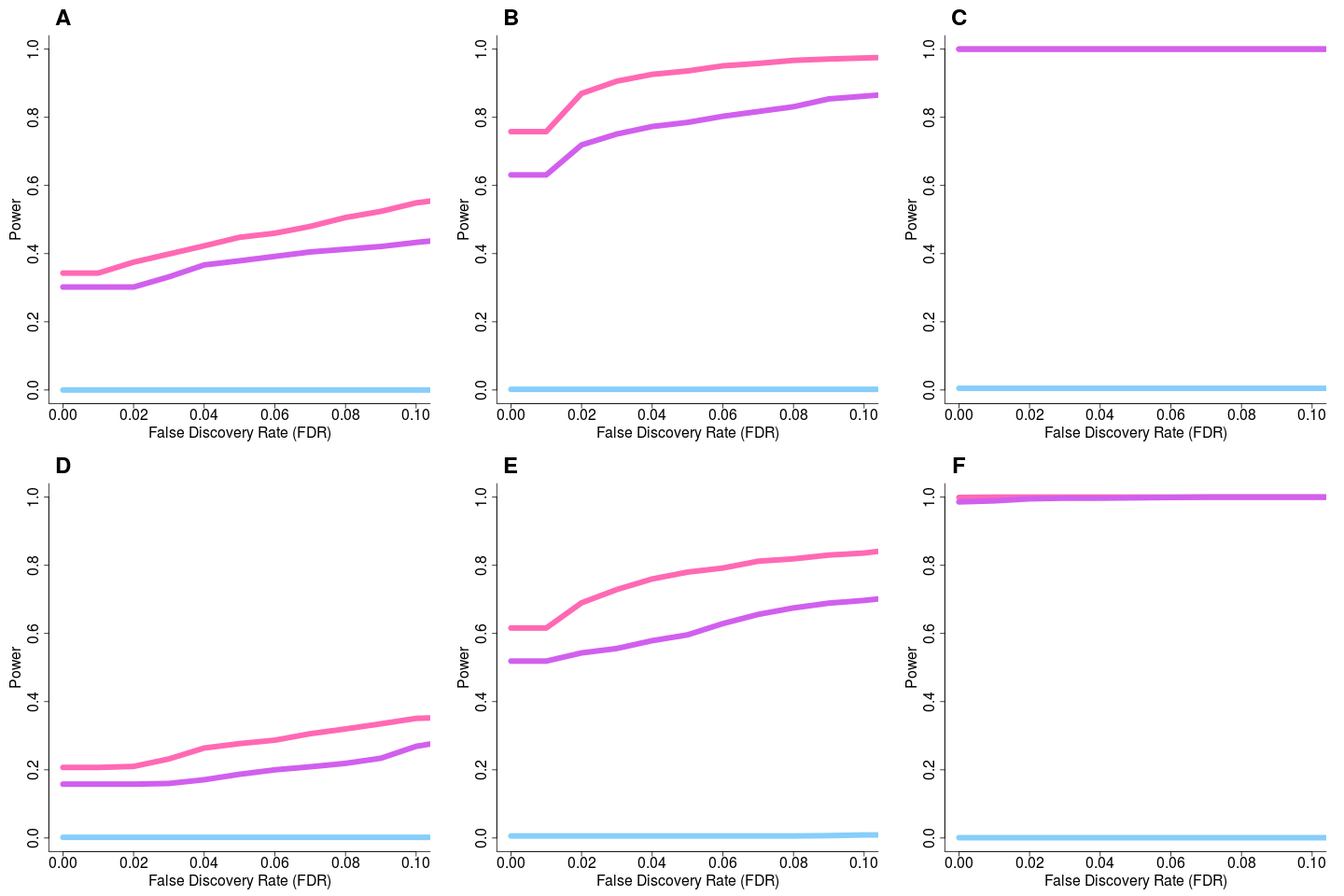

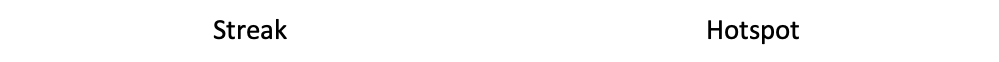

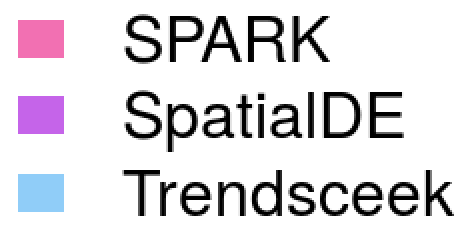

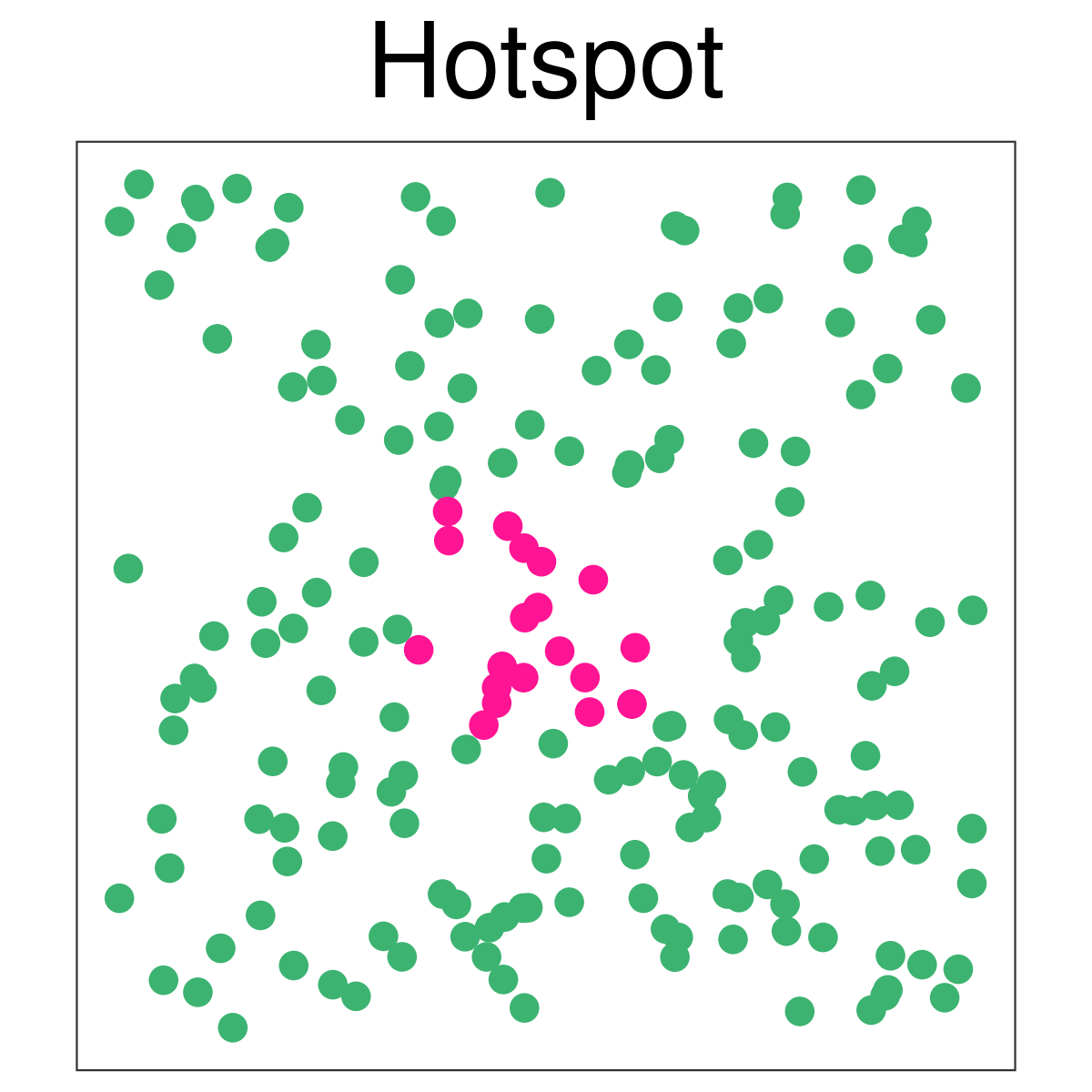

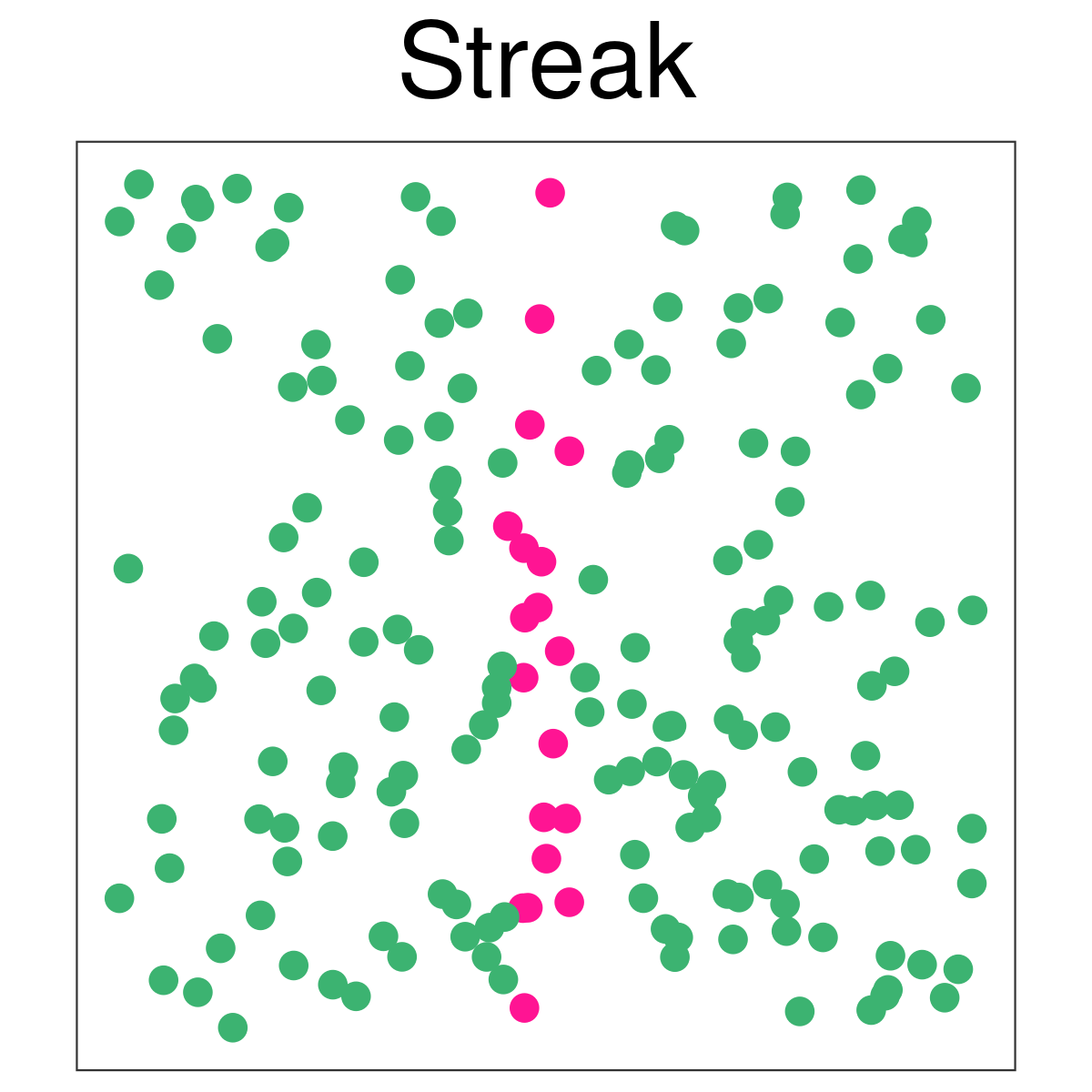

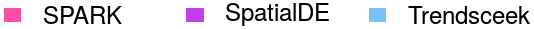

**Supplementary Figure 5: Power comparison of different methods in alternative simulations under different SE strength in the simulations based on the original Trendsceek paper.** Power plots show the proportion of true positives (y-axis) detected by different methods at a range of false discovery rates (FDR; x-axis) in the alternative simulations. The proportion of true positives is averaged across ten simulation replicates. Pattern strength (SE strength) is defined as the fold change between mean expression value in spiked cells and mean expression value in background cells. Compared methods include SPARK (pink), SpatialDE (purple), Trendsceek (sky-blue) which is combined test of Trendsceek. Simulations were performed for the hotspot pattern (first row) or the non-radial streak pattern (second row) under different SE strength: weak SE strength (1.5 fold change; first column), moderate SE strength (2 fold change; second column), and strong SE strength (2.5 fold change; third column). In all settings, the sample size is set to be 200 and the fraction of marked cells is set to be moderate (20%). When SE strength is weak, SPARK and SpatialDE have similar power and both are more powerful than Trendsceek (first column). When SE strength is moderate or strong, SPARK is more powerful than the other two methods for detecting genes with spatial expression patterns (last two columns).

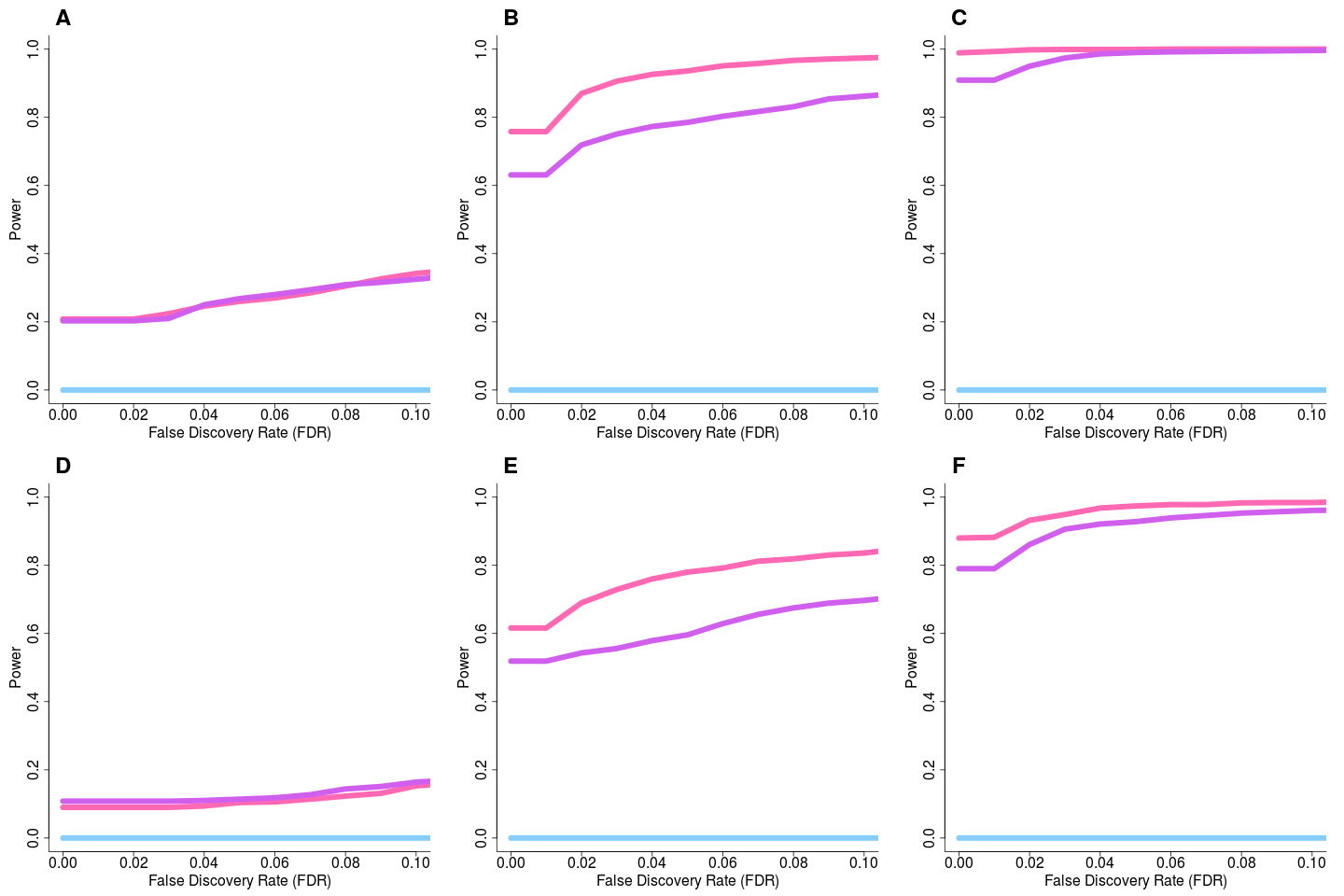

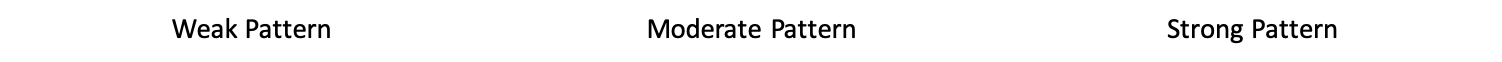

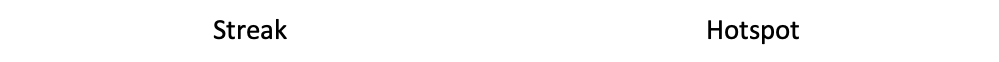

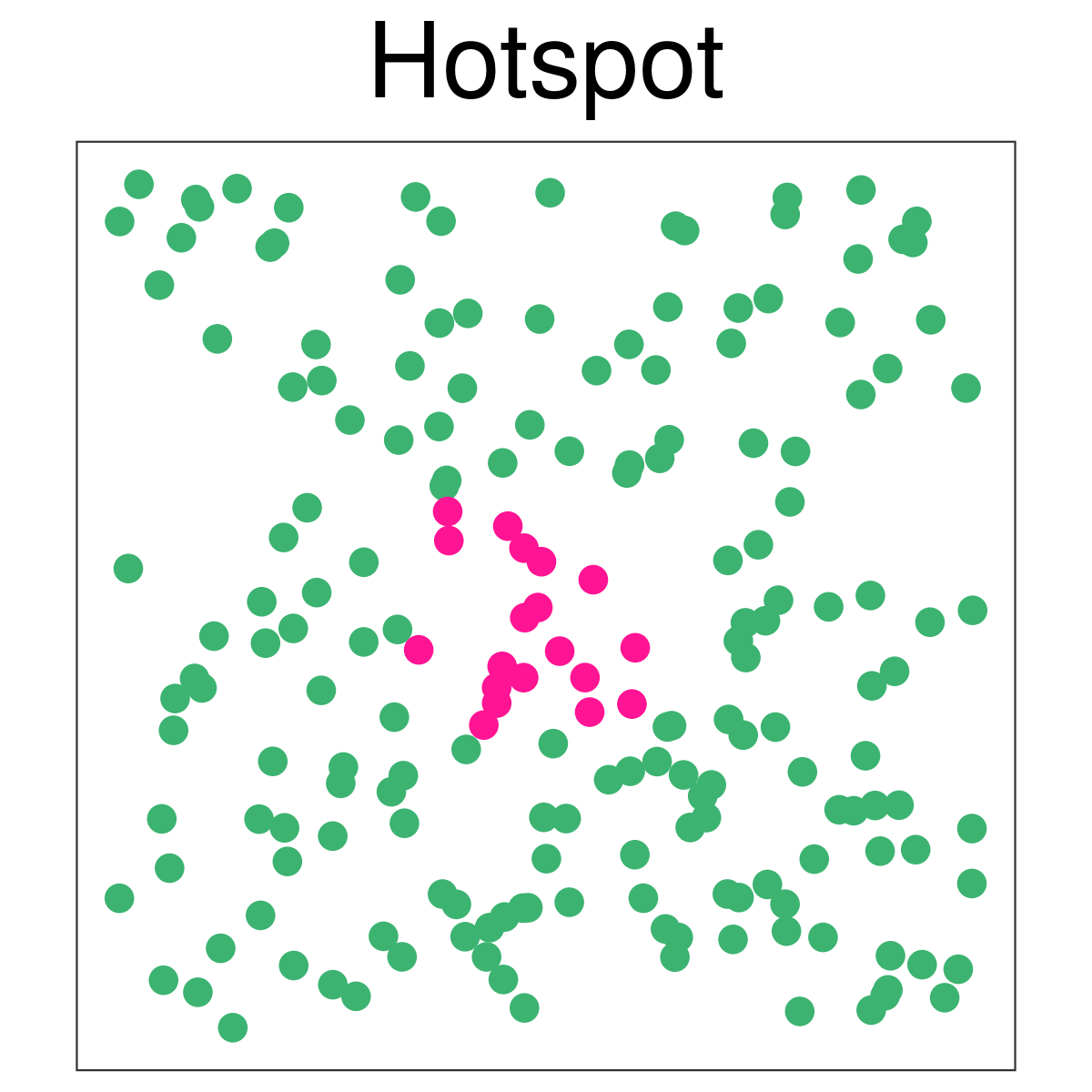

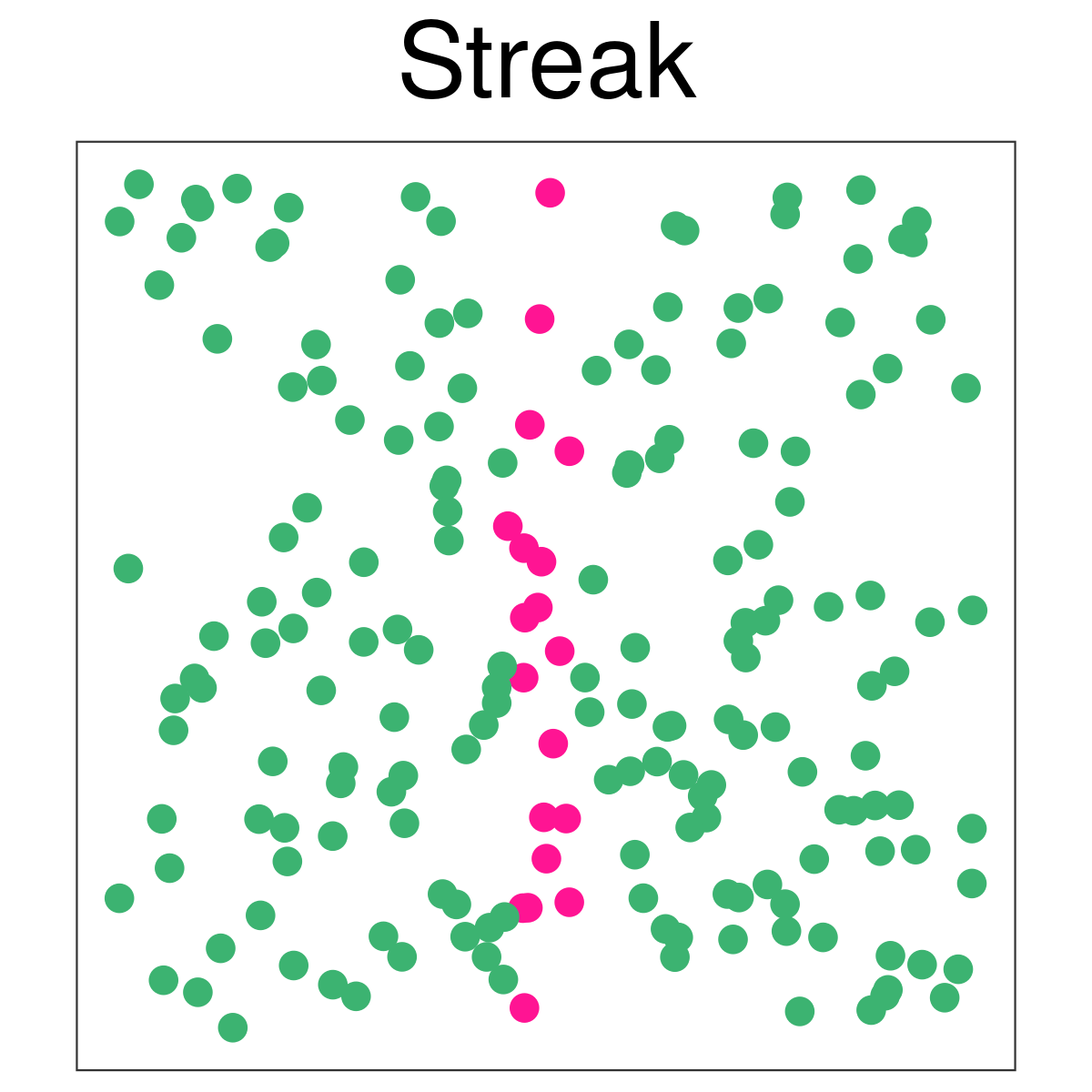

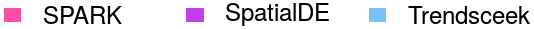

**Supplementary Figure 6: Power comparison of different methods in alternative simulations under different fraction of marked cells in the simulations based on the original Trendsceek paper.** Power plots show the proportion of true positives (y-axis) detected by different methods at a range of false discovery rates (FDR; x-axis) in the alternative simulations. The proportion of true positives is averaged across ten simulation replicates. Compared methods include SPARK (pink), SpatialDE (purple), Trendsceek (sky-blue) which is combined test of Trendsceek. Simulations were performed for the hotspot pattern (first row) or the non-radial streak pattern (second row) under different fraction of cells: 0.1 in (first column); 0.2 in (second column); and 0.3 (third column). In all settings, the sample size is set to be 200 and the SE strength is set to be moderate (2 fold change). When fraction of marked cells is small, SpatialDE is more powerful than SPARK, while both SPARK and SpatialDE are more powerful than Trendsceek (first column). In the case where fraction of spiked cells is moderate or large, SPARK is more powerful than the other two methods for detecting genes with spatial expression patterns (last two columns).

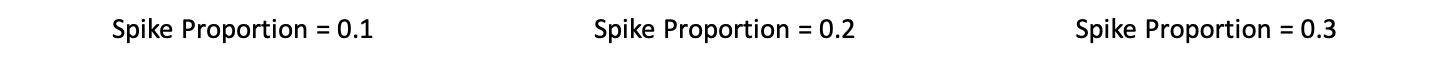

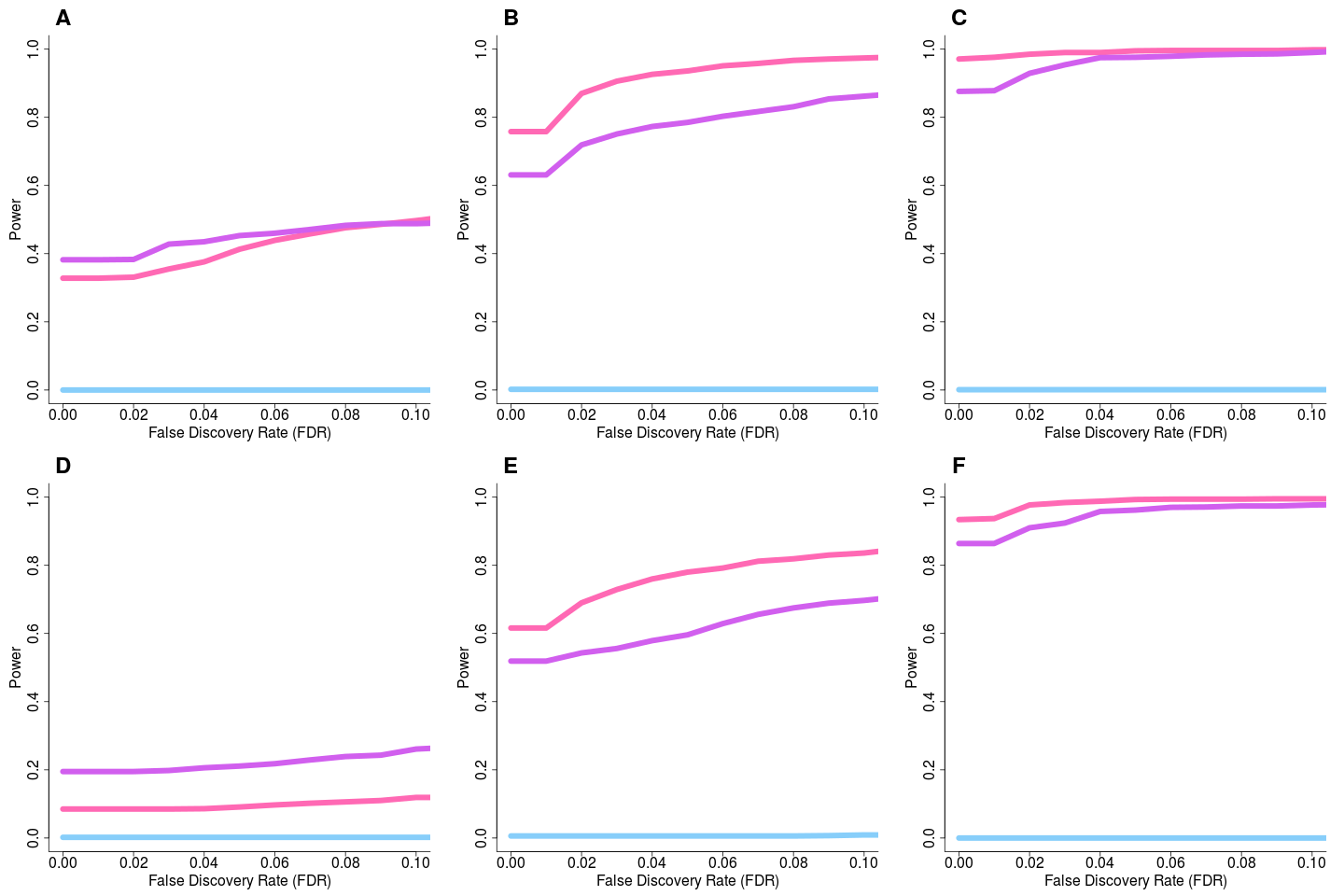

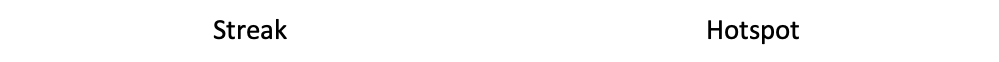

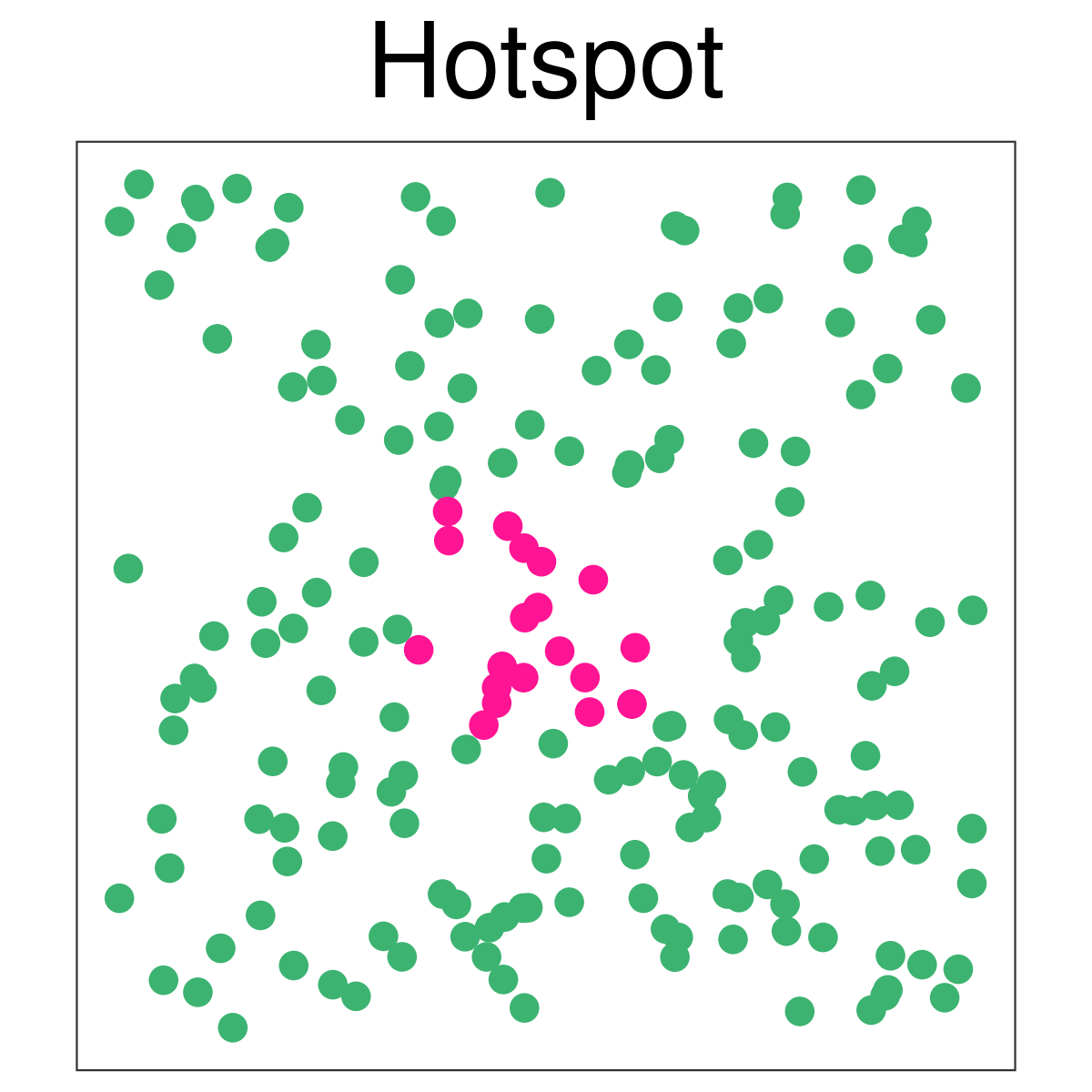

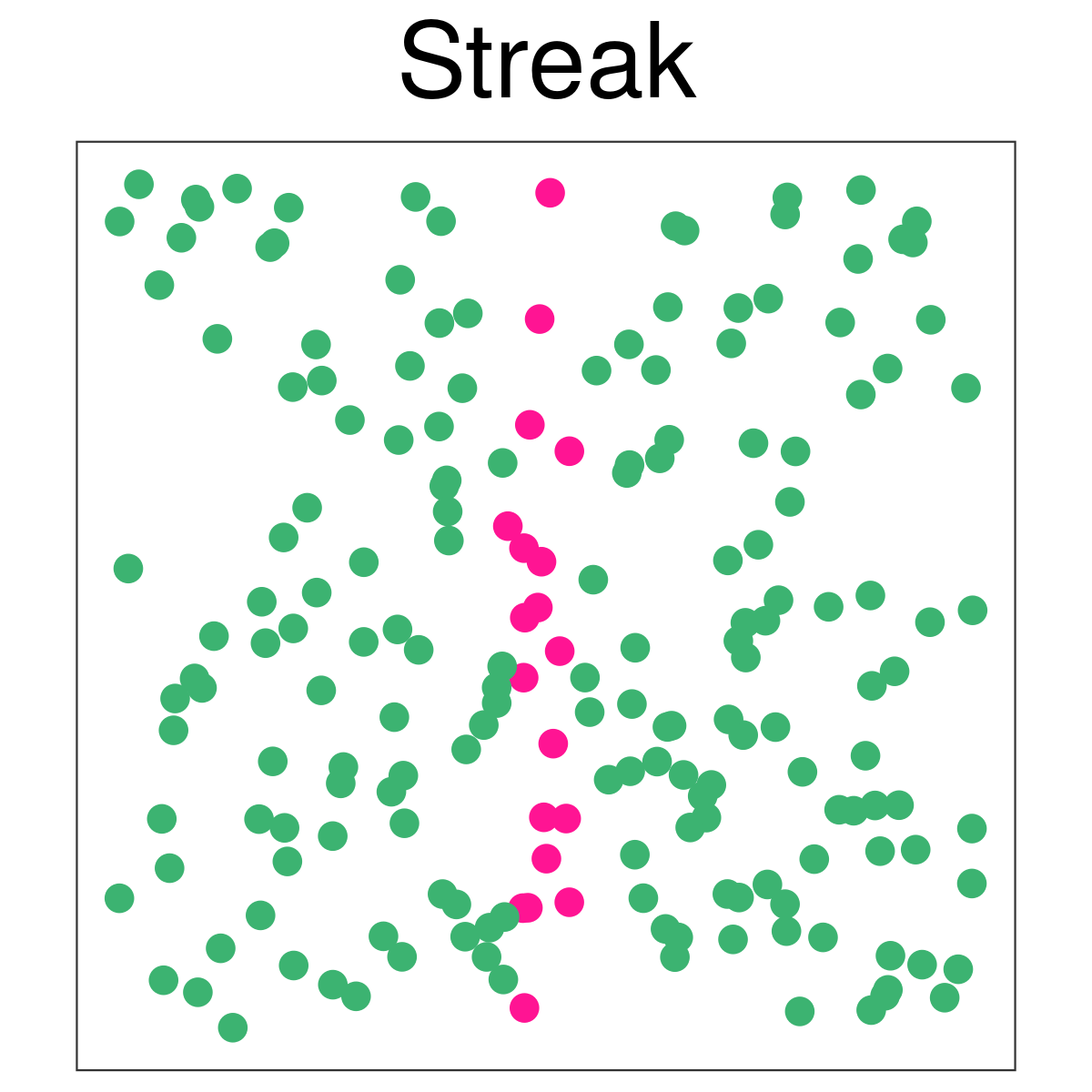

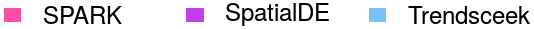

**Supplementary Figure 7: Power comparison of different methods in linear gradient simulations under different sample size and SE signal strength in the simulations based on the original Trendsceek paper.** Power plots show the proportion of true positives (y-axis) detected by different methods at a range of false discovery rates (FDR; x-axis) in the alternative simulations. The proportion of true positives is averaged across ten simulation replicates. Compared methods include SPARK (pink), SpatialDE (purple), Trendsceek (sky-blue) which is combined test of Trendsceek. Linear gradient pattern is illustrated in (I). Simulations were performed under different sample size: low (100; first column); moderate (200; second column); and high (500; third column). Simulations were also performed under three SE signal strength levels: weak SE strength (30% cells are marked cells that display expression gradient), moderate SE strength (40%), and strong SE strength (50%). When sample size is small or moderate, SPARK is more powerful than the other two methods for detecting genes with spatial expression patterns (first two columns). In the case where sample size is high, both SPARK and SpatialDE reach close to 100% power and are more powerful than Trendsceek (last column).

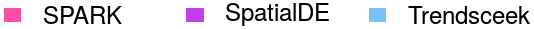

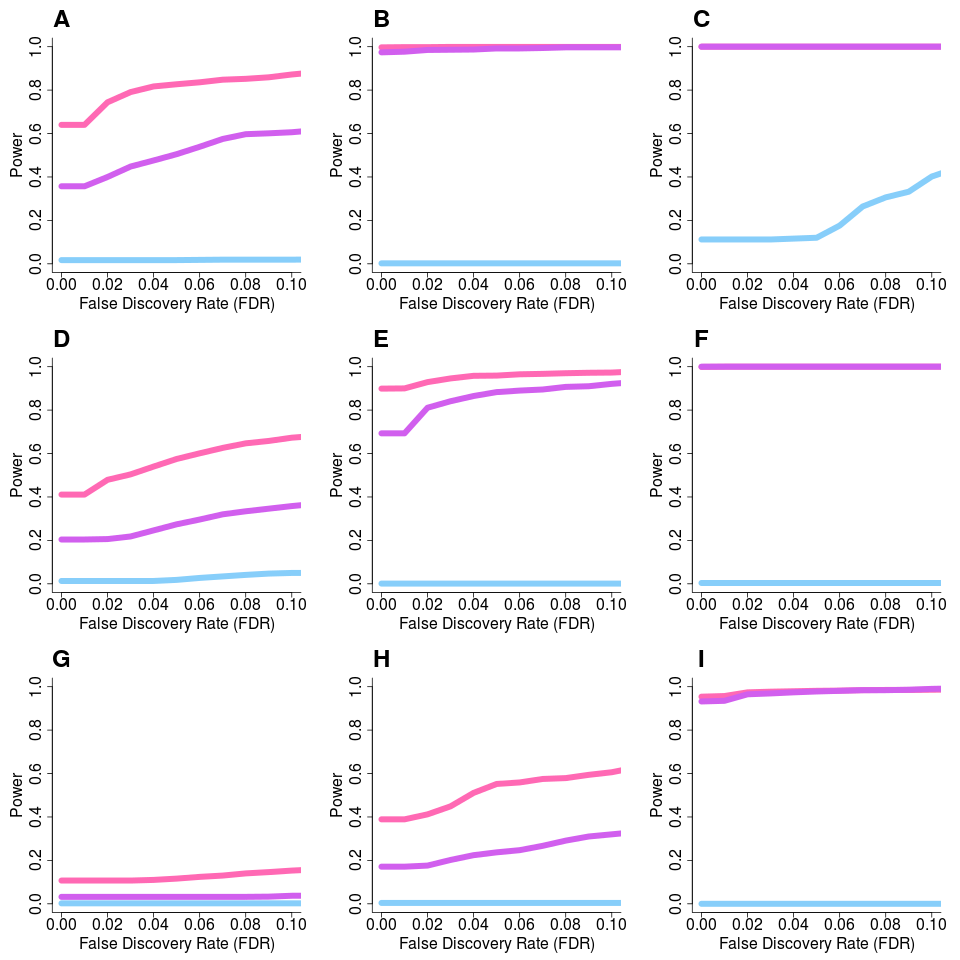

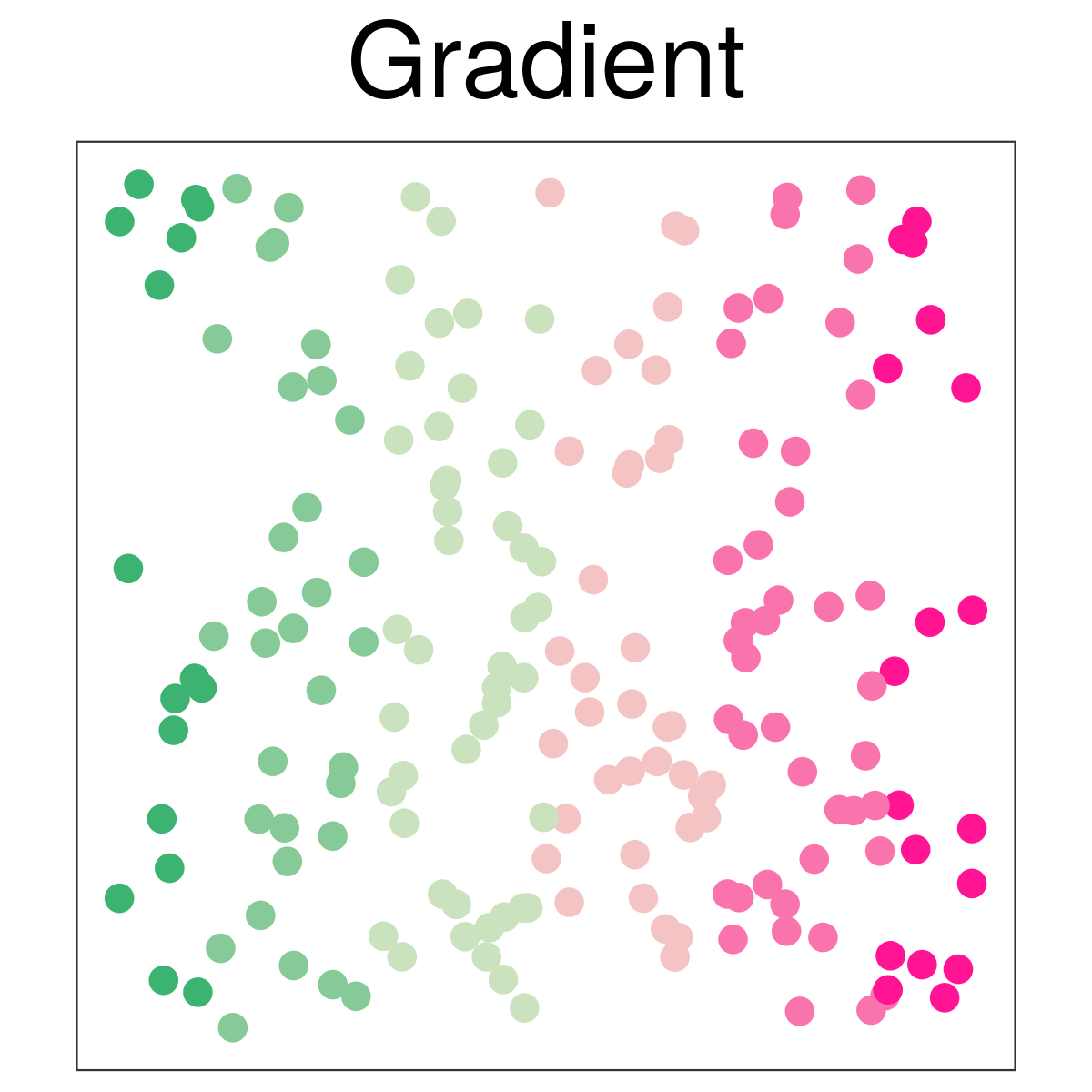

**Supplementary Figure 8: Power comparison of different methods in alternative simulations under different sample sizes when the SE signal strength is extremely strong in the simulations based on the original Trendsceek paper.** Power plots show the proportion of true positives (y-axis) detected by different methods at a range of false discovery rates (FDR; x-axis) in the alternative simulations. The proportion of true positives is averaged across ten simulation replicates. Compared methods include SPARK (pink), SpatialDE (purple), Trendsceek (sky-blue) which is combined test of Trendsceek. Simulations were performed for hotspot pattern (first row) and non-radial streak pattern (second row) under different sample size: low (100; first column); moderate (200; second column); and high (500; third column). In all settings, the fraction of spike cells is set to be 0.2 and the SE strength is set to be extremely high: 5 fold. Even under such high SE strength, Trendsceek only displays low power.

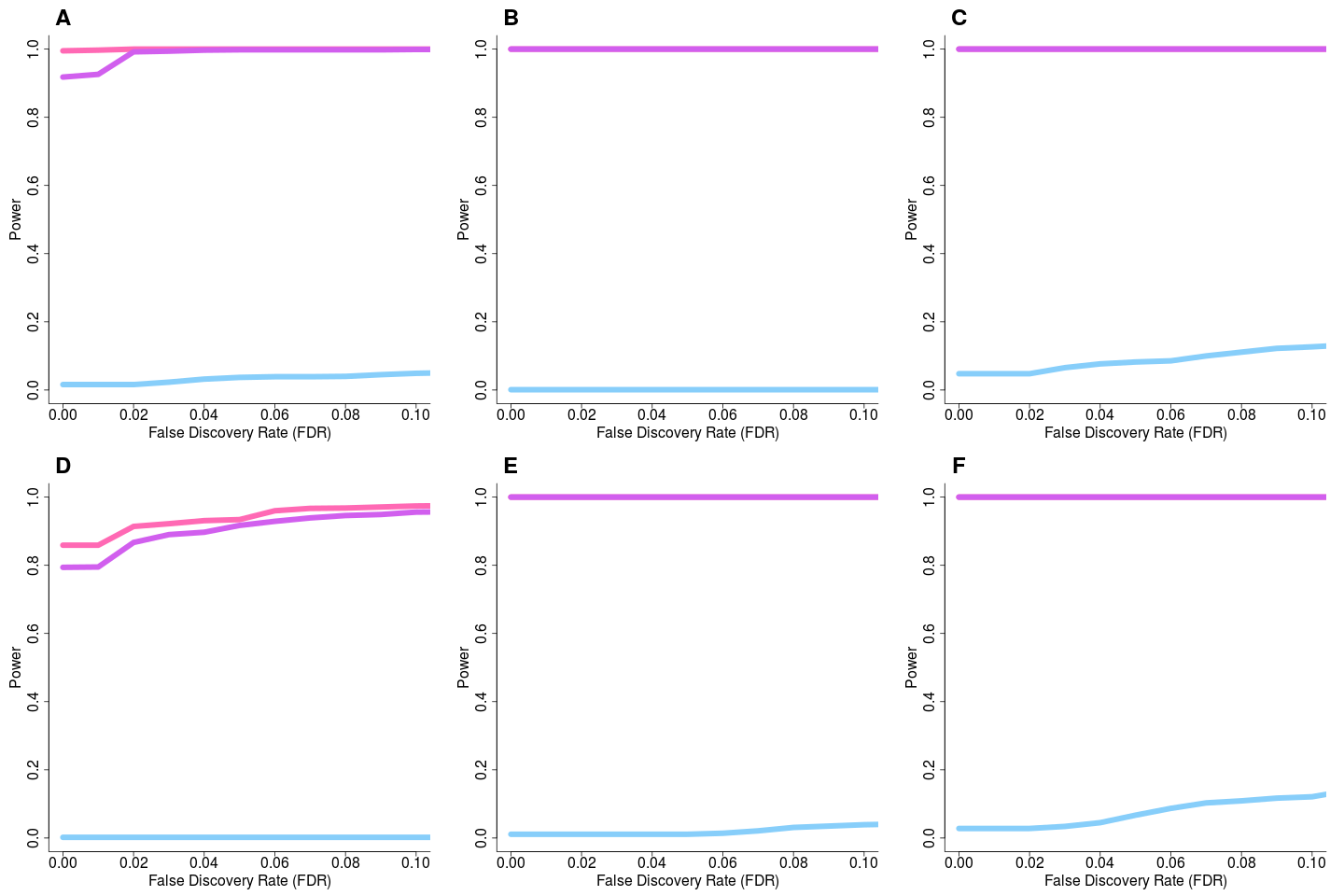

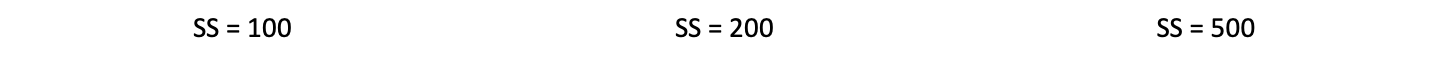

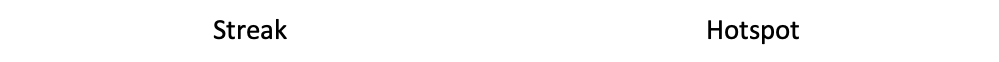

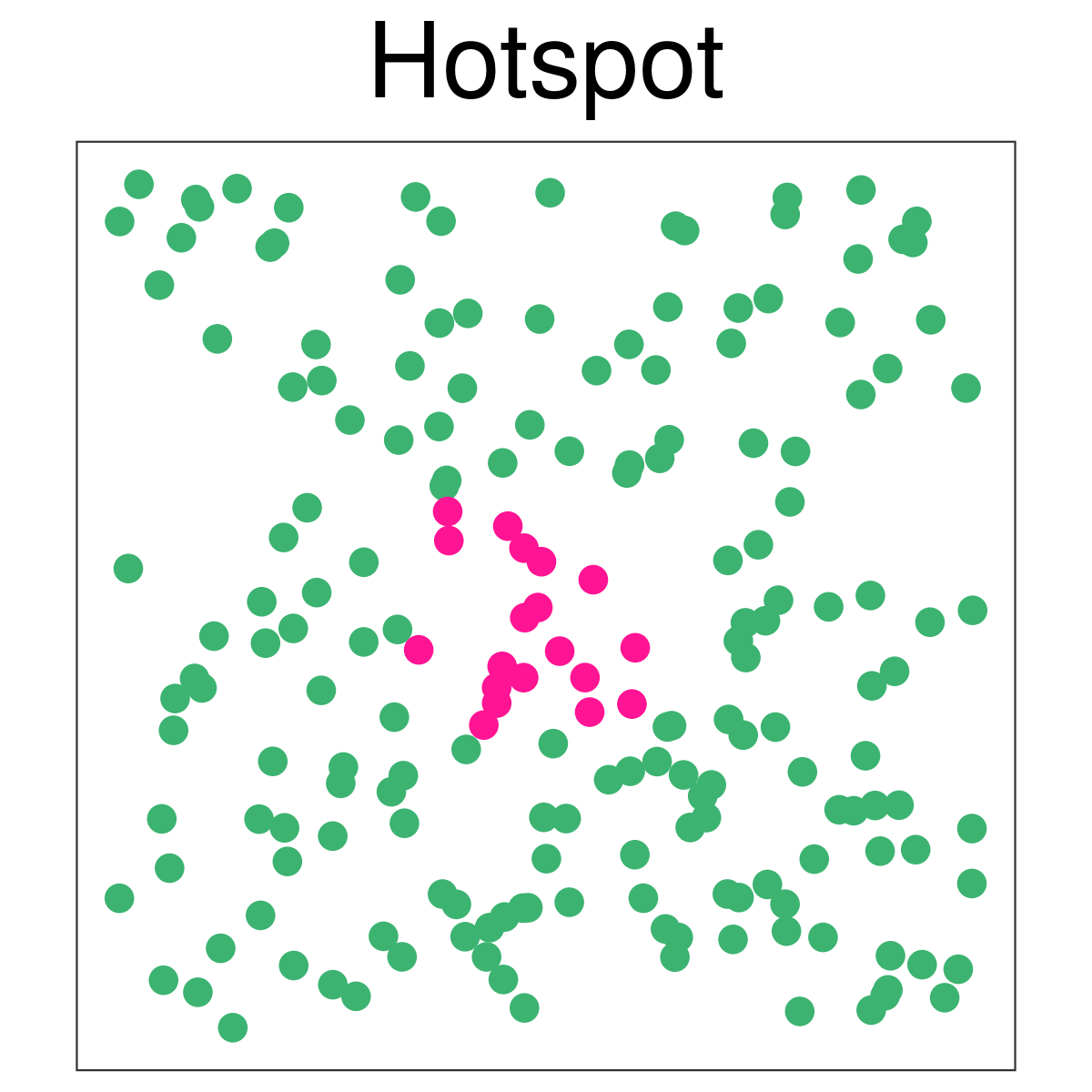

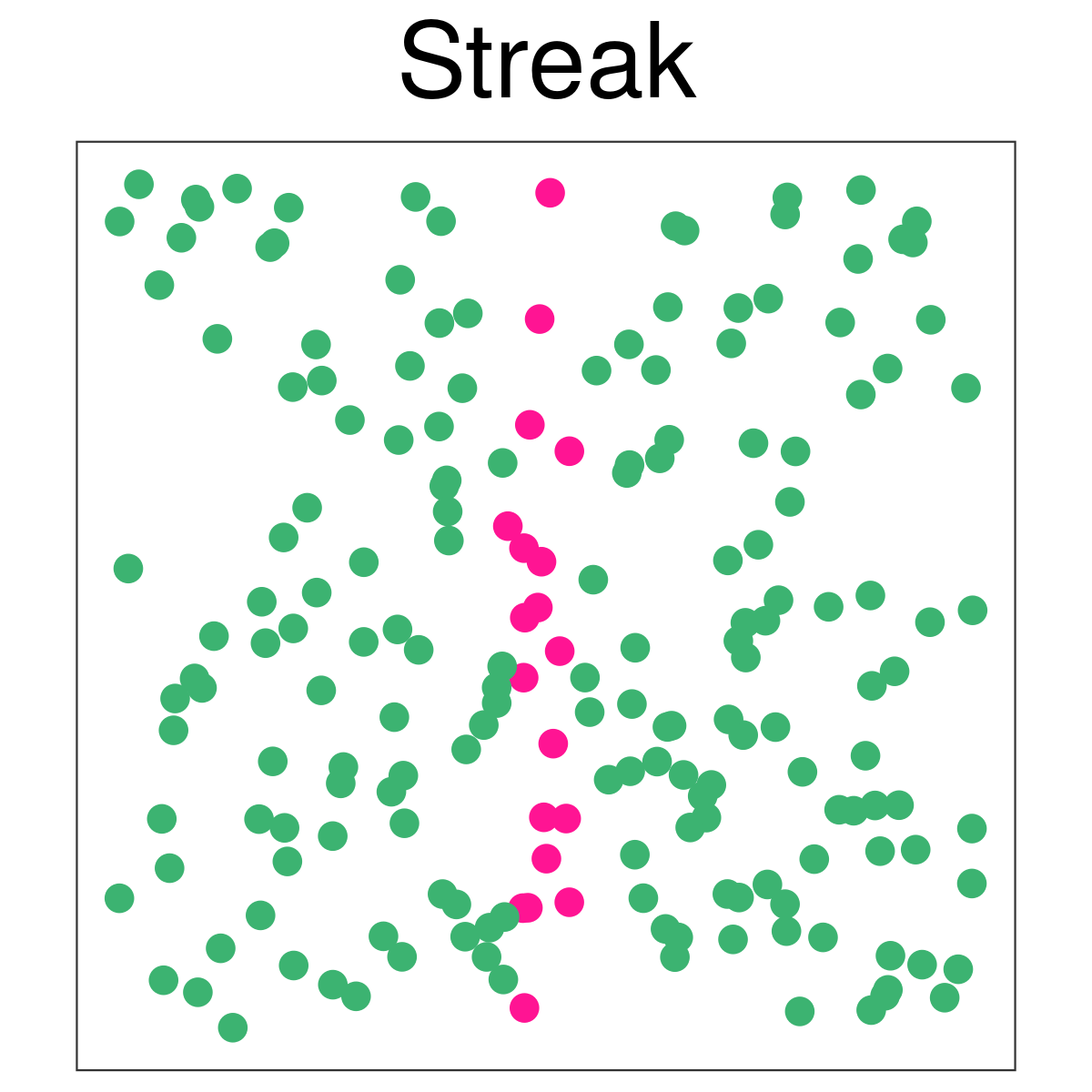

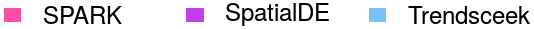

**Supplementary Figure 9: Quantile-quantile plot for the mouse olfactory bulb data.** Quantile-quantile plot of the observed -log10 *p*-values from different methods against the expected -log10 *p*-values under the null in the real data analysis. *p*-value distribution suggests that SPARK detected more genes with spatial expression pattern than SpatialDE. In contrast, *p*-values from various Trendsceek tests all behave like null and suggest a lack of power by Trendsceek tests.

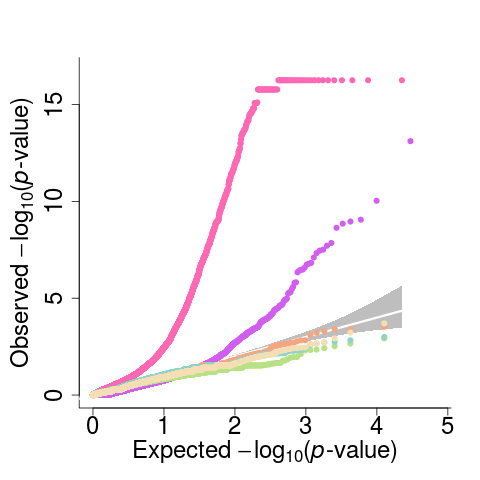

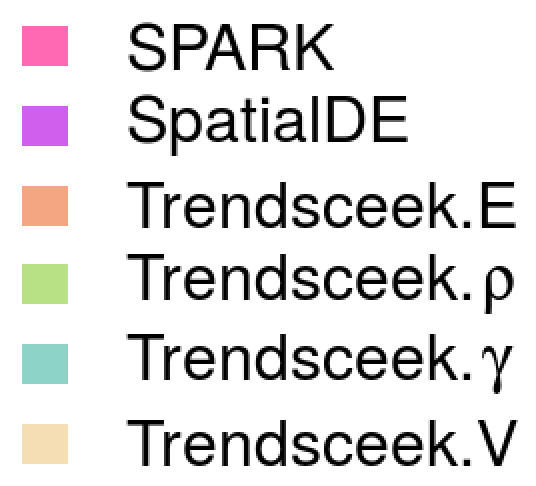

**Supplementary Figure 10: Boxplot displays expression level for the significant genes identified by different methods in the mouse olfactory bulb data.** Results are shown for genes that are detected by SPARK only (first column), genes that are detected by SpatialDE only (middle column), and genes that are detected by both methods. Gene expression level is computed on the log2 scale.

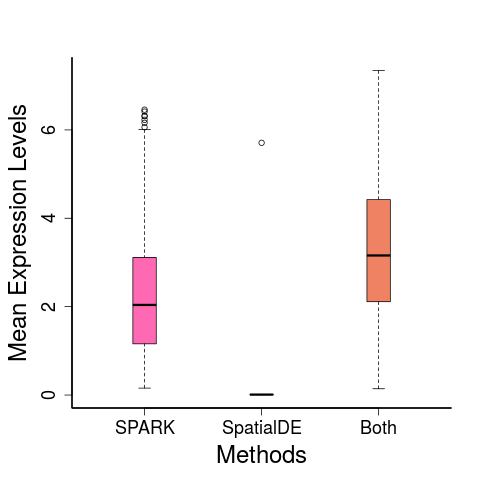

**Supplementary Figure 11: Spatial expression pattern for five genes that are only identified by SpatialDE in mouse olfactory bulb data.** Genes are identified based on an FDR cutoff of 0.05. Color represents relative gene expression level (purple: high; green: low). Most of these genes are only expressed in one or two spatial locations and may be false signals.

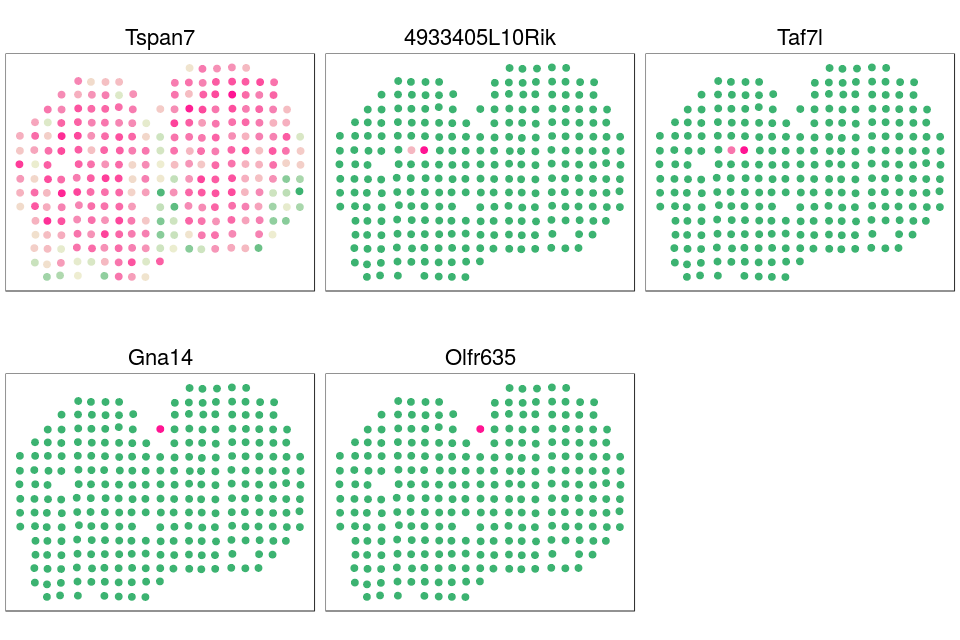

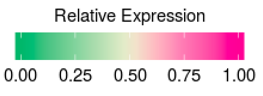

**Supplementary Figure 12: Visualization of significant SE genes identified by SPARK in the mouse olfactory data.** Scatter plot of 772 significant SE genes identified by SPARK. We first used the UMAP (*umap* R package) to reduce the dimension into two dimensions. Then, we used the cell labels obtained from hcluster to visualize the distribution of cells in each cluster. SE genes are detected based on an FDR cutoff of 0.05.

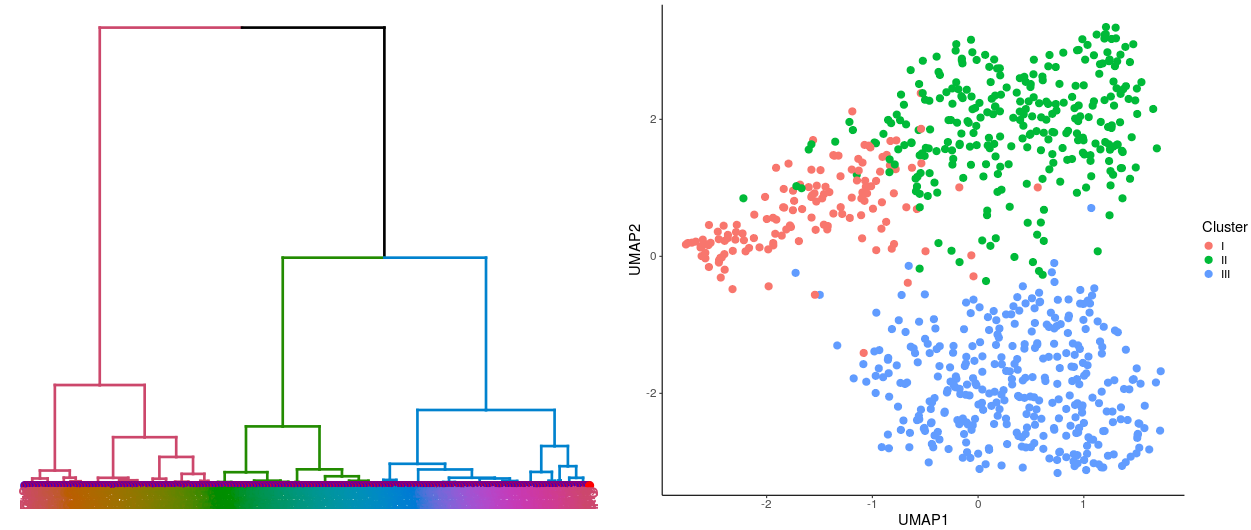

**Supplementary Figure 13: Spatial expression pattern for 20 pattern I genes identified only by SPARK.** Genes are identified based on an FDR cutoff of 0.05. Color represents relative gene expression level (purple: high; green: low). We ranked all 119 pattern I genes identified only by SPARK based on their p-values from the most significant to the least significant. We then obtained 20 genes with increasing *p*-values from ranked list and show these genes here to provide an overview on the spatial expression pattern for genes that are only detected by SPARK. Almost all these 20 genes show clear spatial expression pattern.

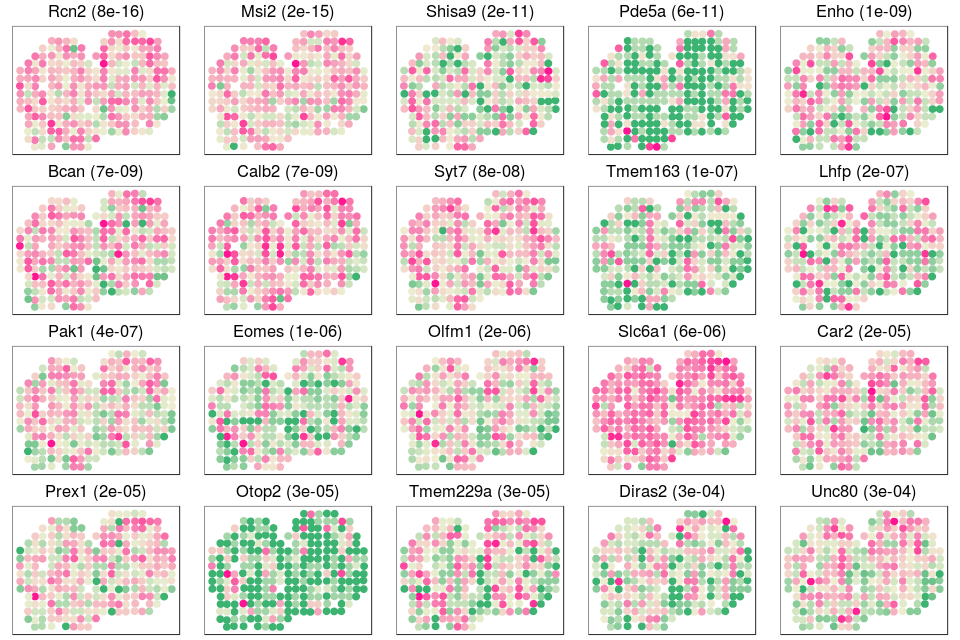

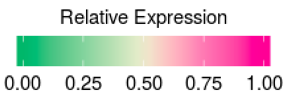

**Supplementary Figure 14: Spatial expression pattern for 20 pattern II genes identified only by SPARK.** Genes are identified based on an FDR cutoff of 0.05. Color represents relative gene expression level (purple: high; green: low). We ranked all 270 pattern II genes identified only by SPARK based on their p-values from the most significant to the least significant. We then obtained 20 genes with increasing *p*-values from ranked list and show these genes here to provide an overview on the spatial expression pattern for genes that are only detected by SPARK. Most of the 20 genes show clear spatial expression pattern.

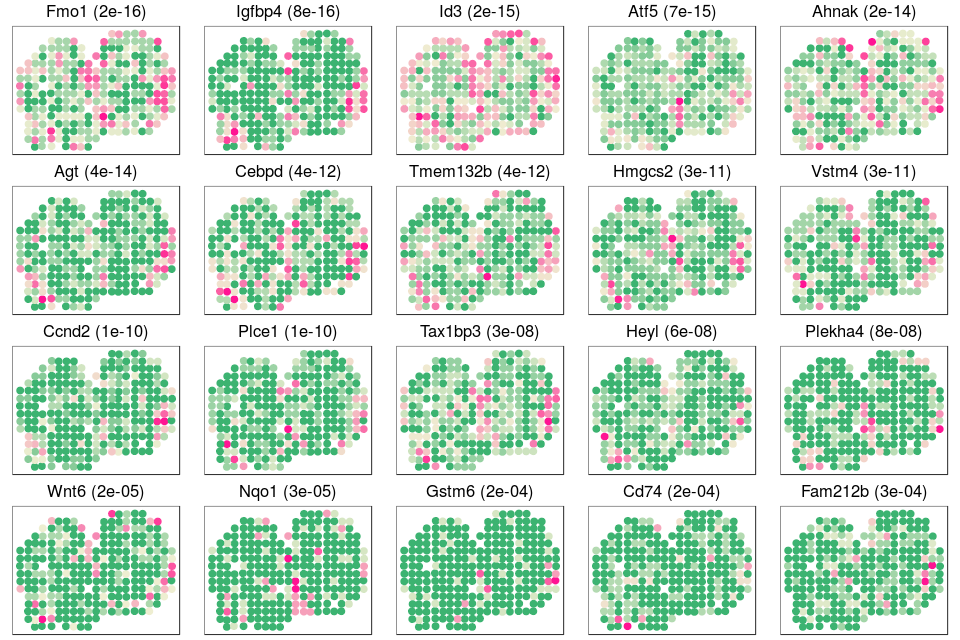

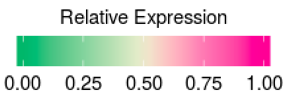

**Supplementary Figure 15: Spatial expression pattern for 20 pattern III genes identified only by SPARK.** Genes are identified based on an FDR cutoff of 0.05. Color represents relative gene expression level (purple: high; green: low). We ranked all 321 pattern III genes identified only by SPARK based on their p-values from the most significant to the least significant. We then obtained 20 genes with increasing *p*-values from ranked list and show these genes here to provide an overview on the spatial expression pattern for genes that are only detected by SPARK. Almost all these 20 genes show clear spatial expression pattern.

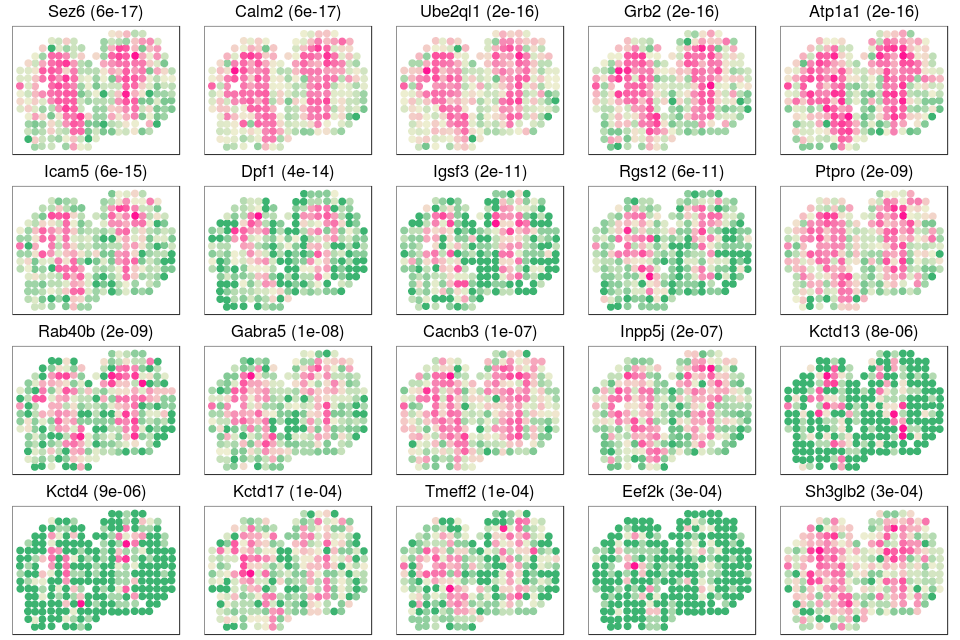

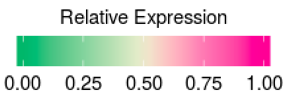

**Supplementary Figure 16: Spatial expression pattern for eight genes identified by SPARK in the mouse olfactory bulb.** Genes are identified based on an FDR cutoff of 0.05. Color represents relative gene expression level (purple: high; green: low). These eight genes are previously known to be spatially expressed in the mitral cell layer. SPARK p-values are shown in parenthesis. Five of these genes (*Nmb, Reln, Rcan2, Shisa3* and *Plcxd2*) were not identified by SpatialDE. None of the eight genes were identified by Trendsceek.

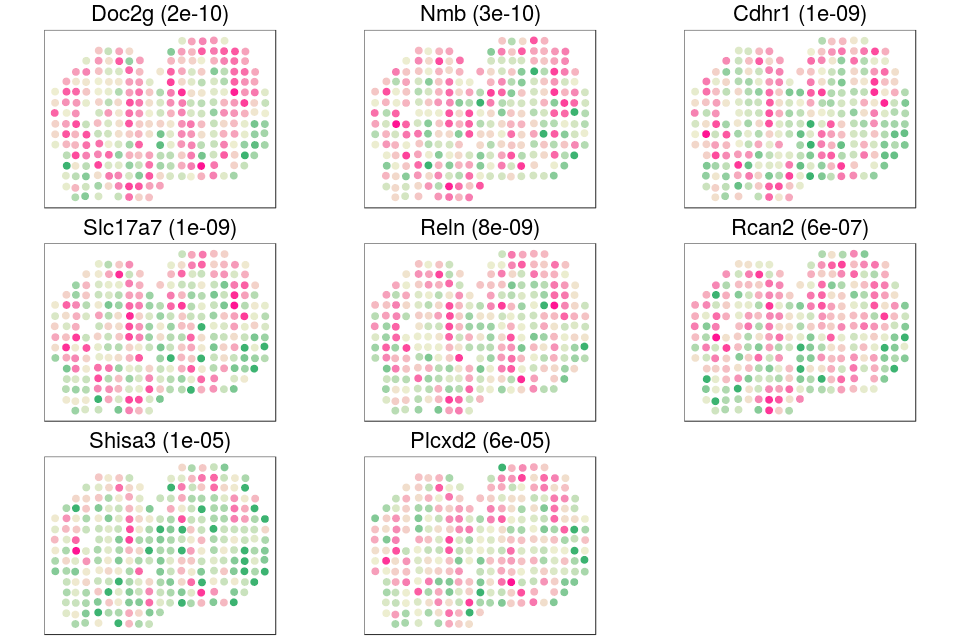

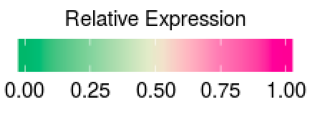

**Supplementary Figure 17: Venn Diagram and Bubble Plot of functional enrichment analyses of SE genes identified by SPARK and SpatialDE for mouse olfactory bulb data.** (**A**) Venn Diagram of enriched GO terms based on the SE genes identified by SPARK and SpatialDE. The SE genes identified by SPARK are enriched in 1023 GO terms, 64 of which are overlapped with SpatialDE. (**B**) Venn Diagram of enriched KEGG pathways based on the SE genes identified by SPARK and SpatialDE. The SE genes identified by SPARK are enriched in 79 KEGG pathways, which cover the two KEGG pathways enriched by SpatialDE identified SE genes. (**C**) Bubble plot shows –log10 *p*-values for gene sets uniquely identified by SPARK (959). Gene sets are colored by three categories: GO biological process (blue), GO molecular function (purple), GO cellular component (yellow).

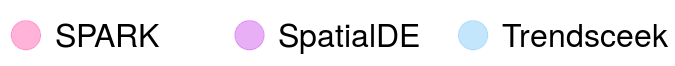

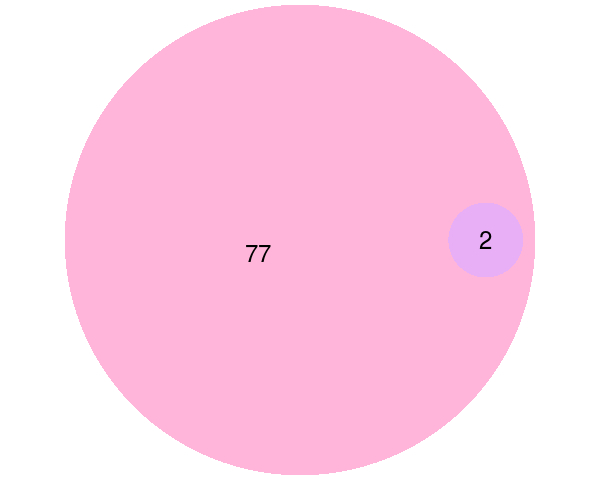

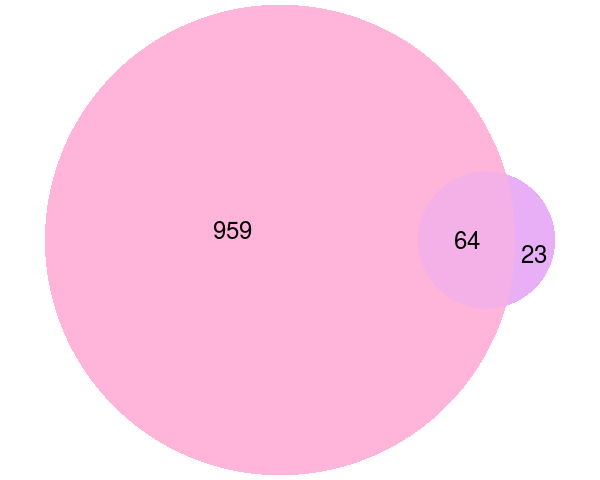

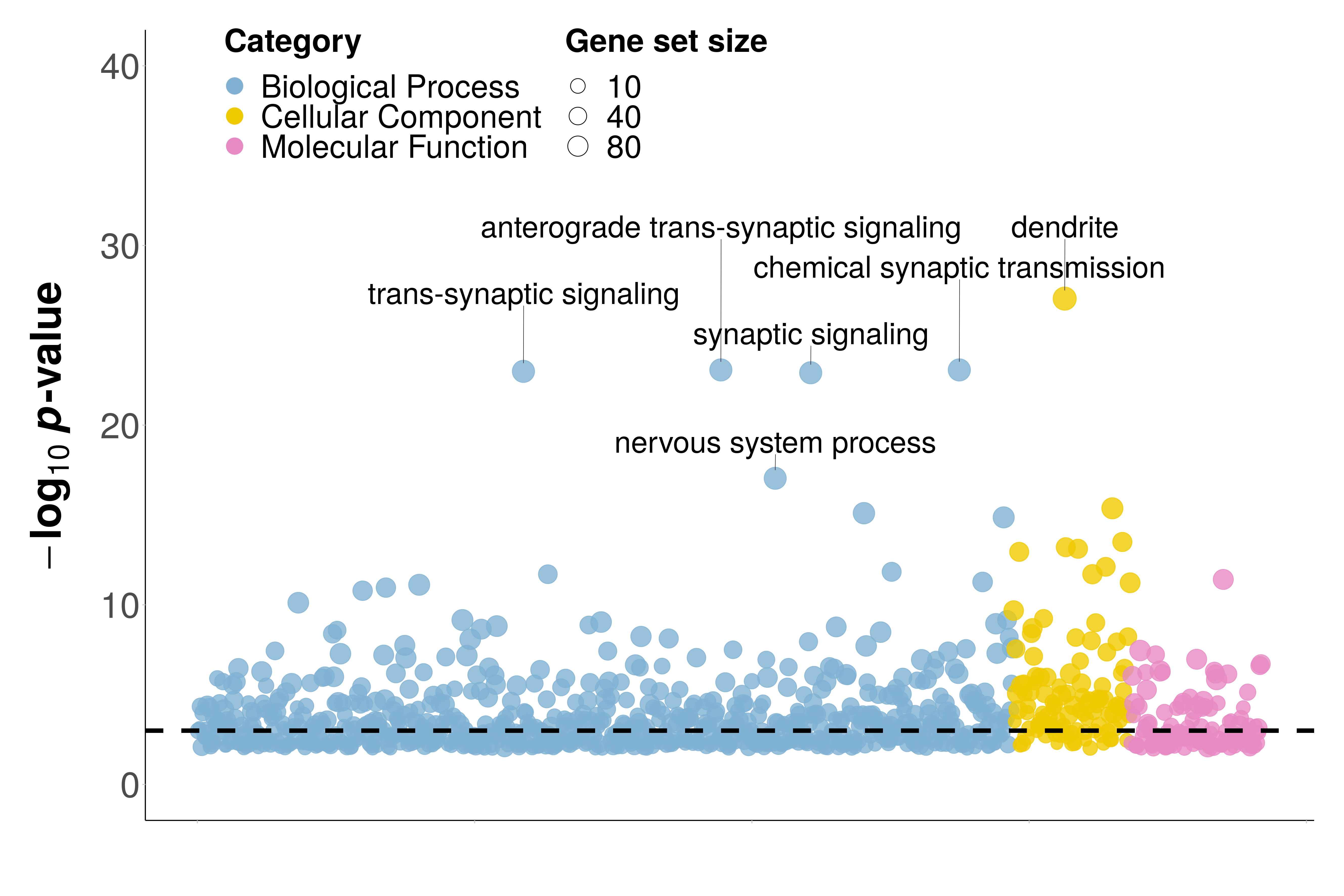

**A**

**B**

**C**

**Supplementary Figure 18: Venn Diagram of functional enrichment analyses of SE genes identified by SPARK and SpatialDE for three patterns in the mouse olfactory bulb data.** (**A**) Venn Diagram of enriched GO terms based on the SE genes identified by SPARK and SpatialDE for pattern I. The SE genes identified by SPARK in pattern I are enriched in 489 GO terms, 96 of which are overlapped with SpatialDE. (**B**) Venn Diagram of enriched GO terms based on the SE genes identified by SPARK and SpatialDE for pattern II. The SE genes identified by SPARK in pattern II are enriched in 714 GO terms, 117 of which are overlapped with SpatialDE. (**C**) Venn Diagram of enriched GO terms based on the SE genes identified by SPARK and SpatialDE for pattern II. The SE genes identified by SPARK in pattern II are enriched in 684 GO terms, 22 of which are overlapped with SpatialDE.

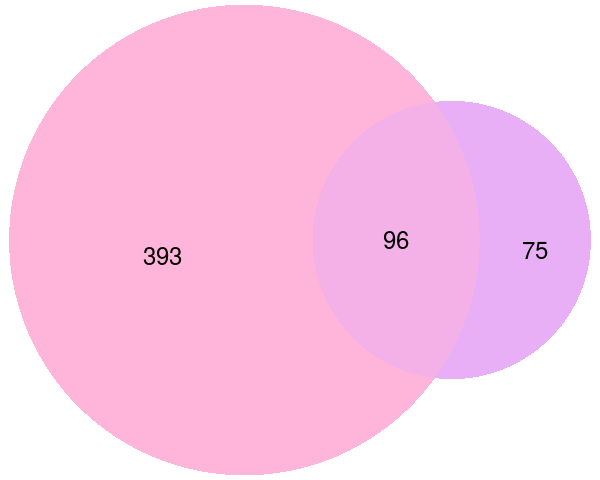

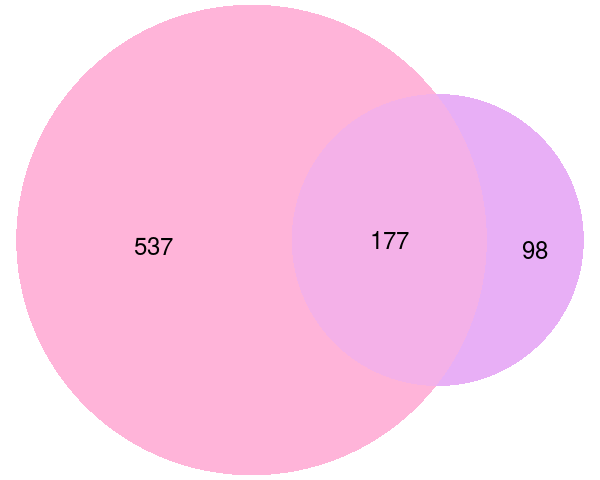

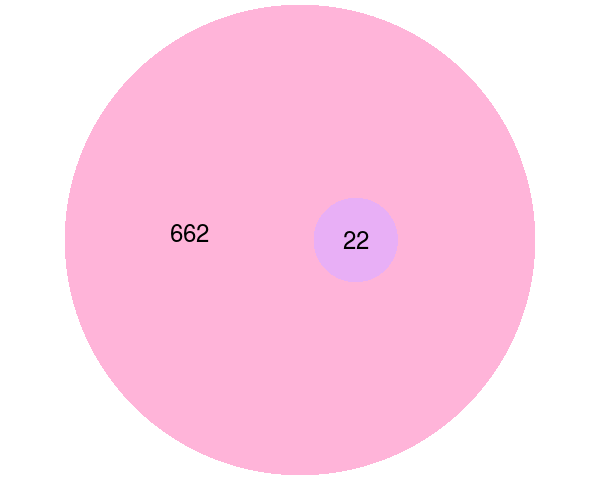

**A**

**B**

**C**

**Supplementary Figure 19: Bubble plot shows –log10 *p*-values for gene sets in pattern I uniquely identified by SPARK (393) in the mouse olfactory bulb data.** Gene sets are colored by three categories: GO biological process (blue), GO molecular function (purple), GO cellular component (yellow).

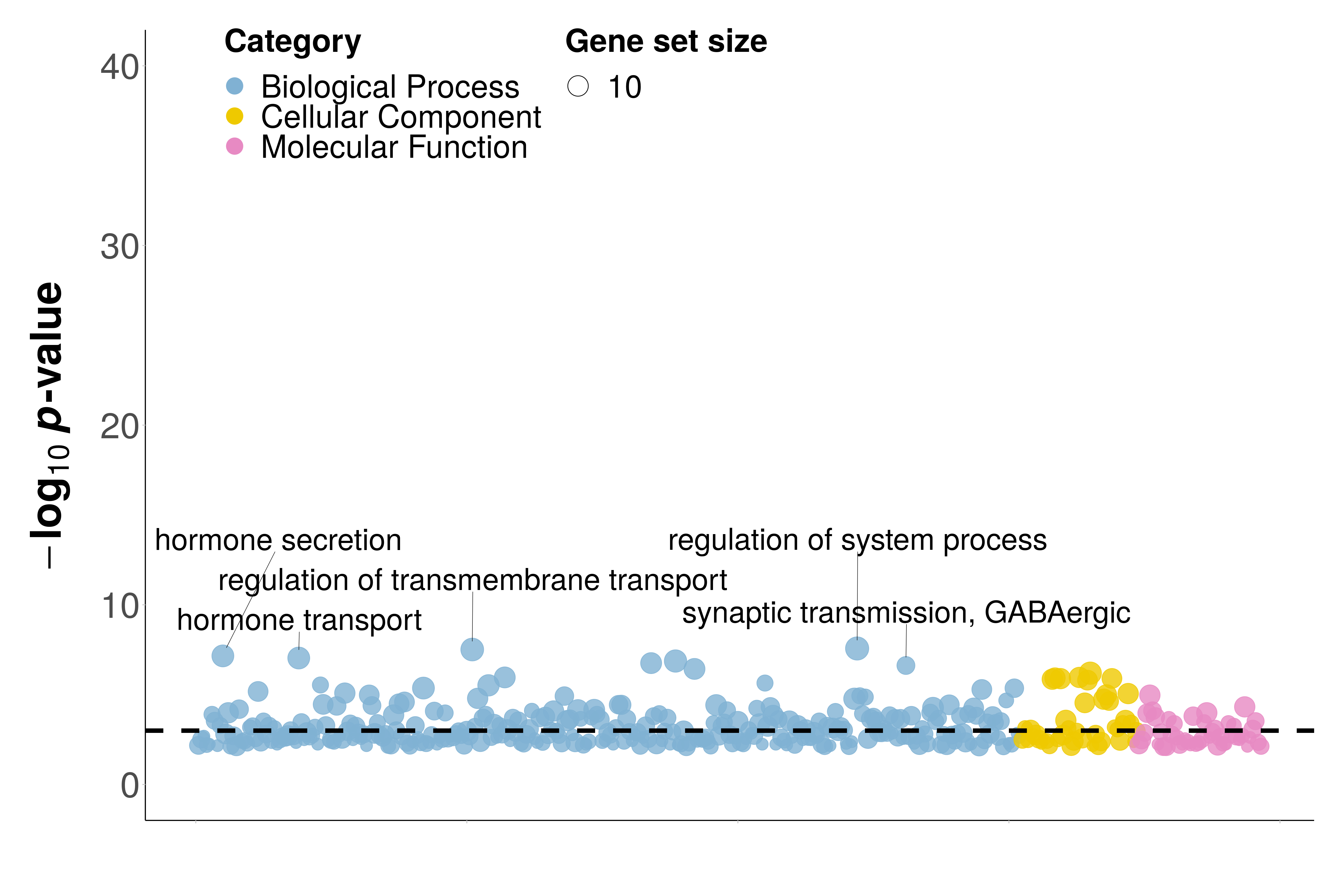

**Supplementary Figure 20: Bubble plot shows –log10 *p*-values for gene sets in pattern II uniquely identified by SPARK (537) in the mouse olfactory bulb data.** Gene sets are colored by three categories: GO biological process (blue), GO molecular function (purple), GO cellular component (yellow).

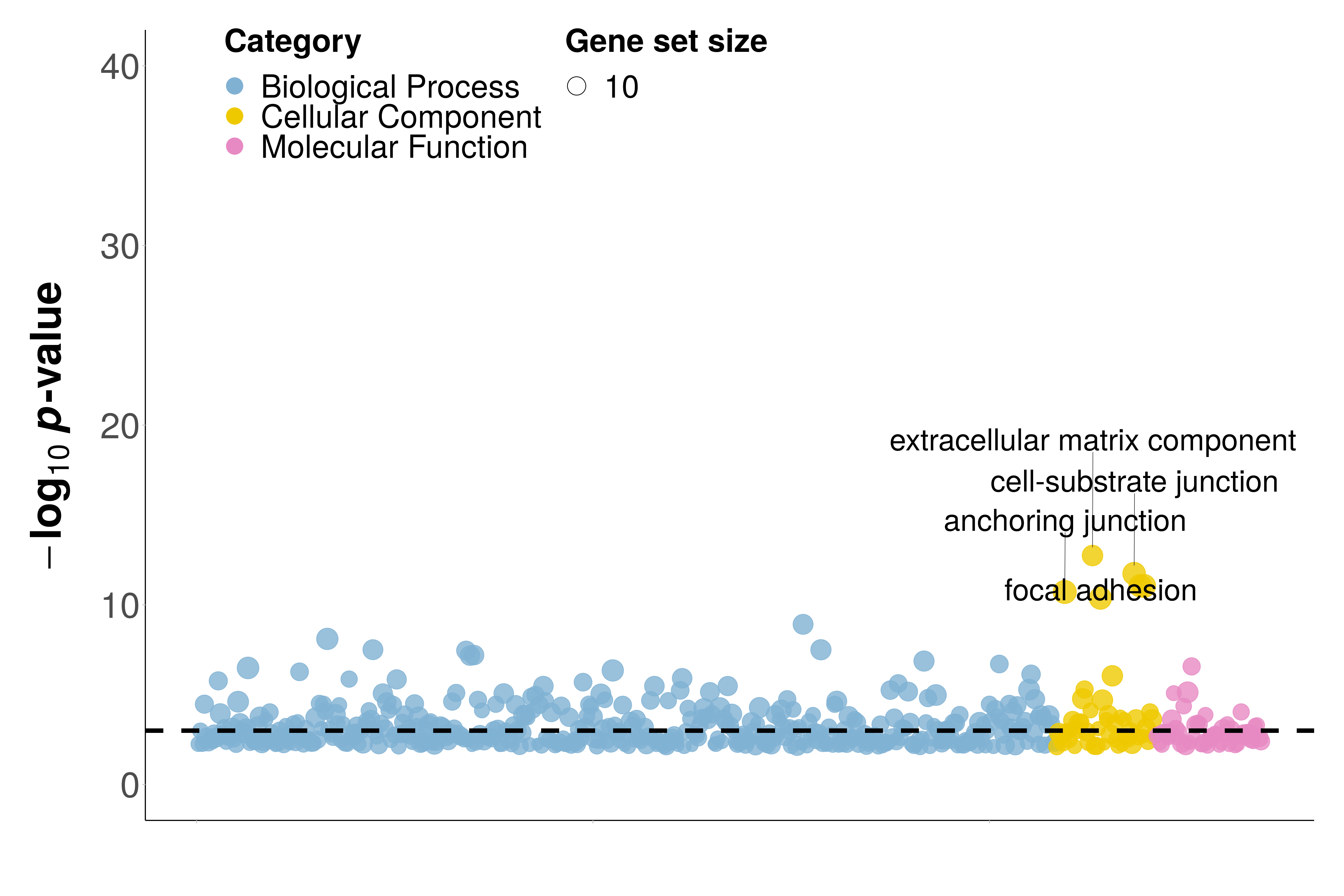

**Supplementary Figure 21: Bubble plot shows –log10 *p*-values for gene sets in pattern III uniquely identified by SPARK (662) in the mouse olfactory bulb data.** Gene sets are colored by three categories: GO biological process (blue), GO molecular function (purple), GO cellular component (yellow).

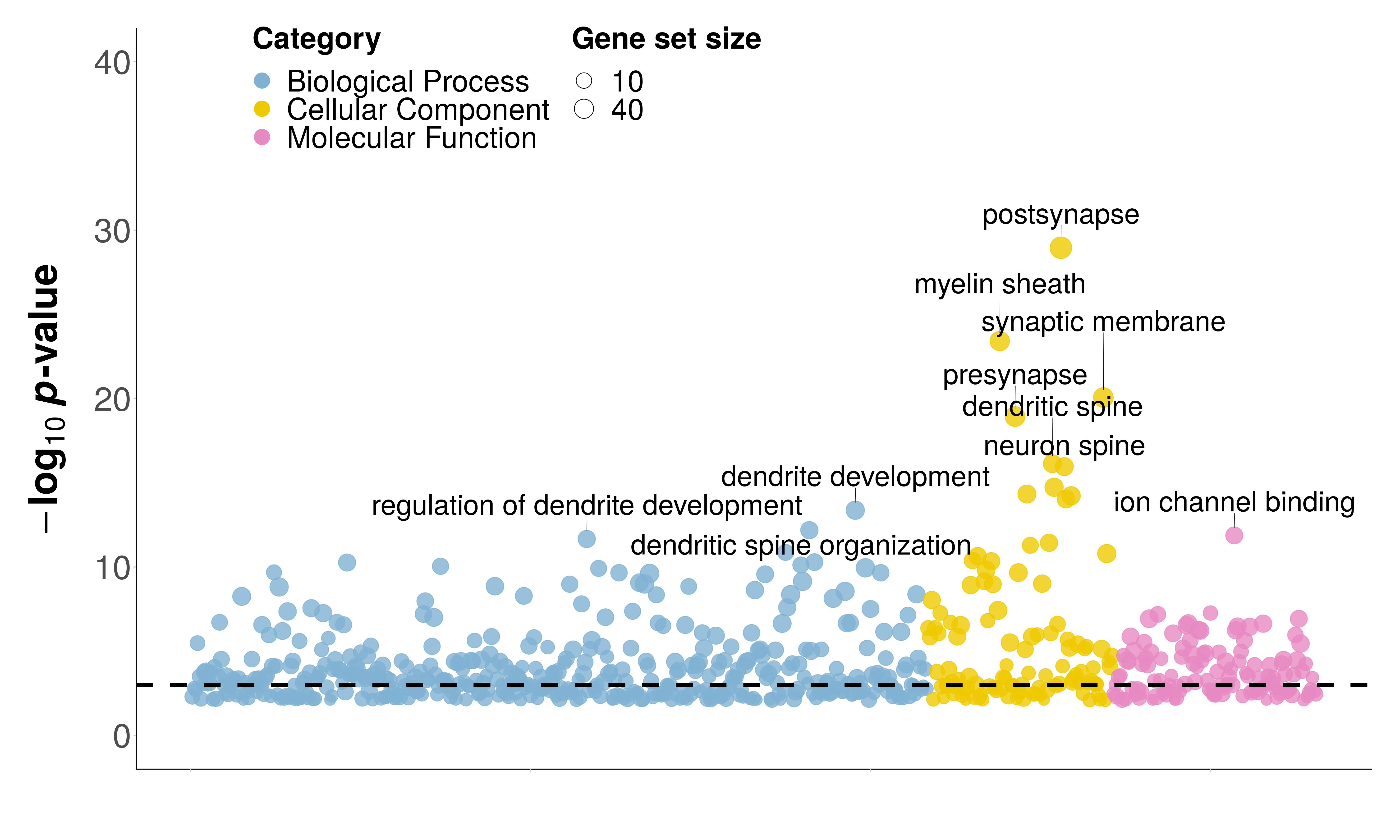

**Supplementary Figure 22: Quantile-Quantile Plot for human breast cancer data.** Quantile-quantile plot of the observed -log10 *p*-values from different methods against the expected -log10 *p*-values under the null in the real data analysis. *p*-value distribution suggests that SPARK detected more genes with spatial expression pattern than SpatialDE. In contrast, *p*-values from various Trendsceek tests all behave approximately like null and suggest a lack of power by Trendsceek tests

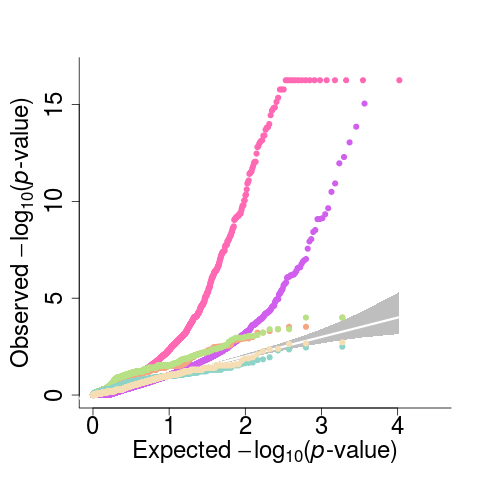

**Supplementary Figure 23: Boxplot displays expression level for the significant genes identified by different methods in the human breast cancer data.** Results are shown for genes that are detected by SPARK only (first column), genes that are detected by SpatialDE only (second column), genes that are detected by Trendsceek only (third column), and genes that are detected by all methods. Gene expression level is computed on the log2 scale.

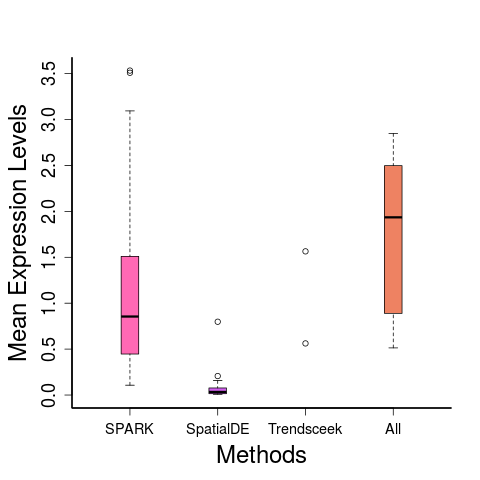

**Supplementary Figure 24: Spatial expression pattern for 20 SE genes identified only by SPARK in the human breast cancer data.** Genes are identified based on an FDR cutoff of 0.05. Color represents relative gene expression level (purple: high; green: low). We ranked all 202 SE genes identified only by SPARK based on their *p*-values from the most significant to the least significant. We then obtained 20 genes with increasing *p*-values from ranked list and show these genes here to provide an overview on the spatial expression pattern for genes that are only detected by SPARK. Almost all these 20 genes show clear spatial expression pattern.

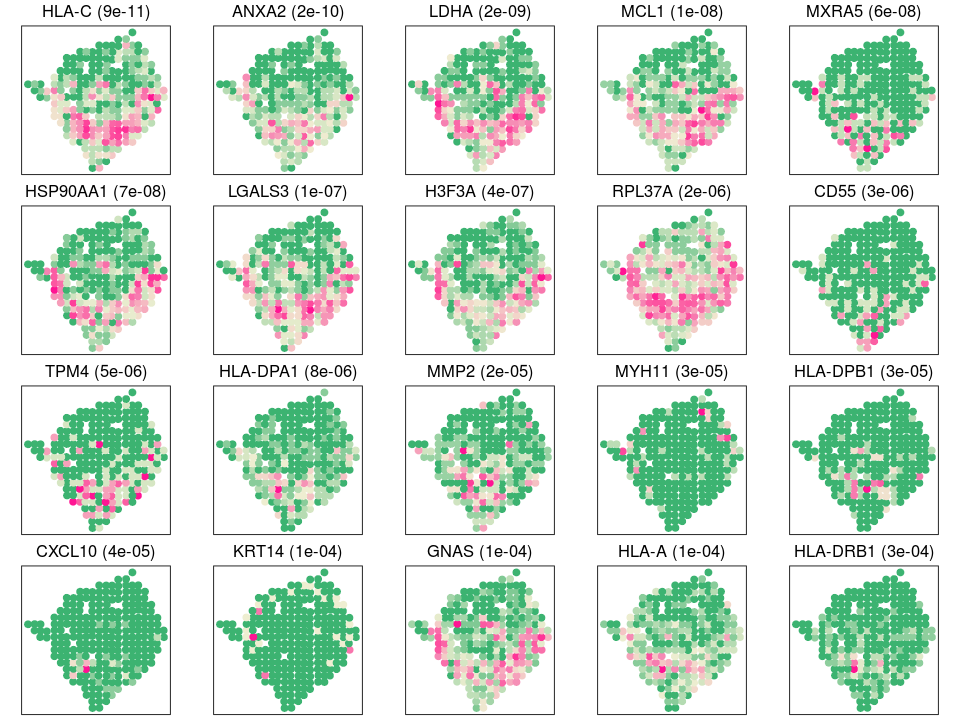

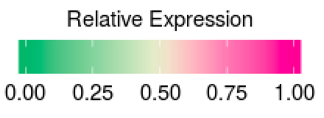

**Supplementary Figure 25: Spatial expression pattern for ten SE genes identified by SPARK in the human breast cancer data.** Genes are identified based on an FDR cutoff of 0.05. Color represents relative gene expression level (purple: high; green: low). These ten genes are cancer relevant genes with known spatial localization highlighted in the original study. SPARK *p*-values are shown in parenthesis. *SCGB2A2*, *KRT17* and *MMP14* were not identified by both SpatialDE and Trendsceek.

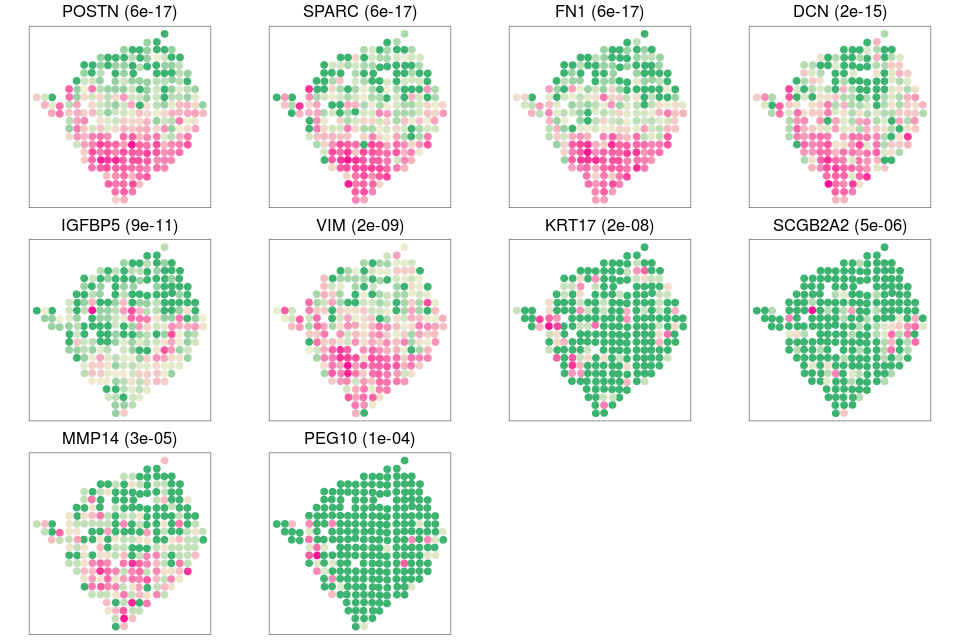

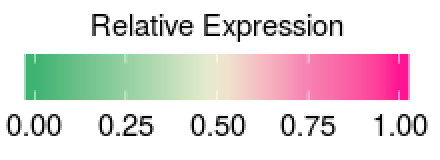

**Supplementary Figure 26: Venn Diagram and Bubble Plot of functional enrichment analyses of SE genes identified by SPARK and SpatialDE in the human breast cancer data.** (**A**) Venn Diagram of enriched GO terms based on the SE genes identified by SPARK and SpatialDE. The SE genes identified by SPARK are enriched in 542 GO terms, 266 of which are overlapped with SpatialDE. (**B**) Venn Diagram of enriched KEGG pathways based on the SE genes identified by SPARK and SpatialDE. The SE genes identified by SPARK are enriched in 20 KEGG pathways, which cover the three KEGG pathways enriched by SpatialDE identified SE genes. (**C**) Bubble plot shows –log10 *p*-values for gene sets uniquely identified by SPARK (351). Gene sets are colored by three categories: GO biological process (blue), GO molecular function (purple), GO cellular component (yellow).

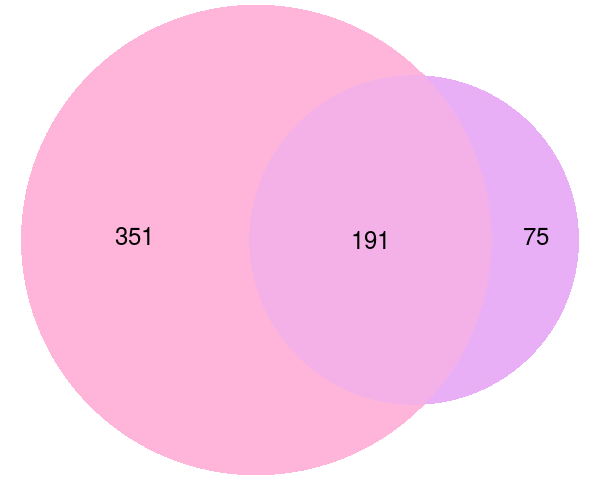

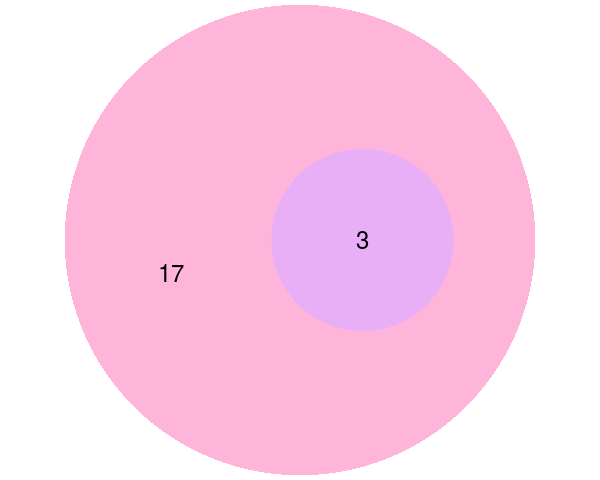

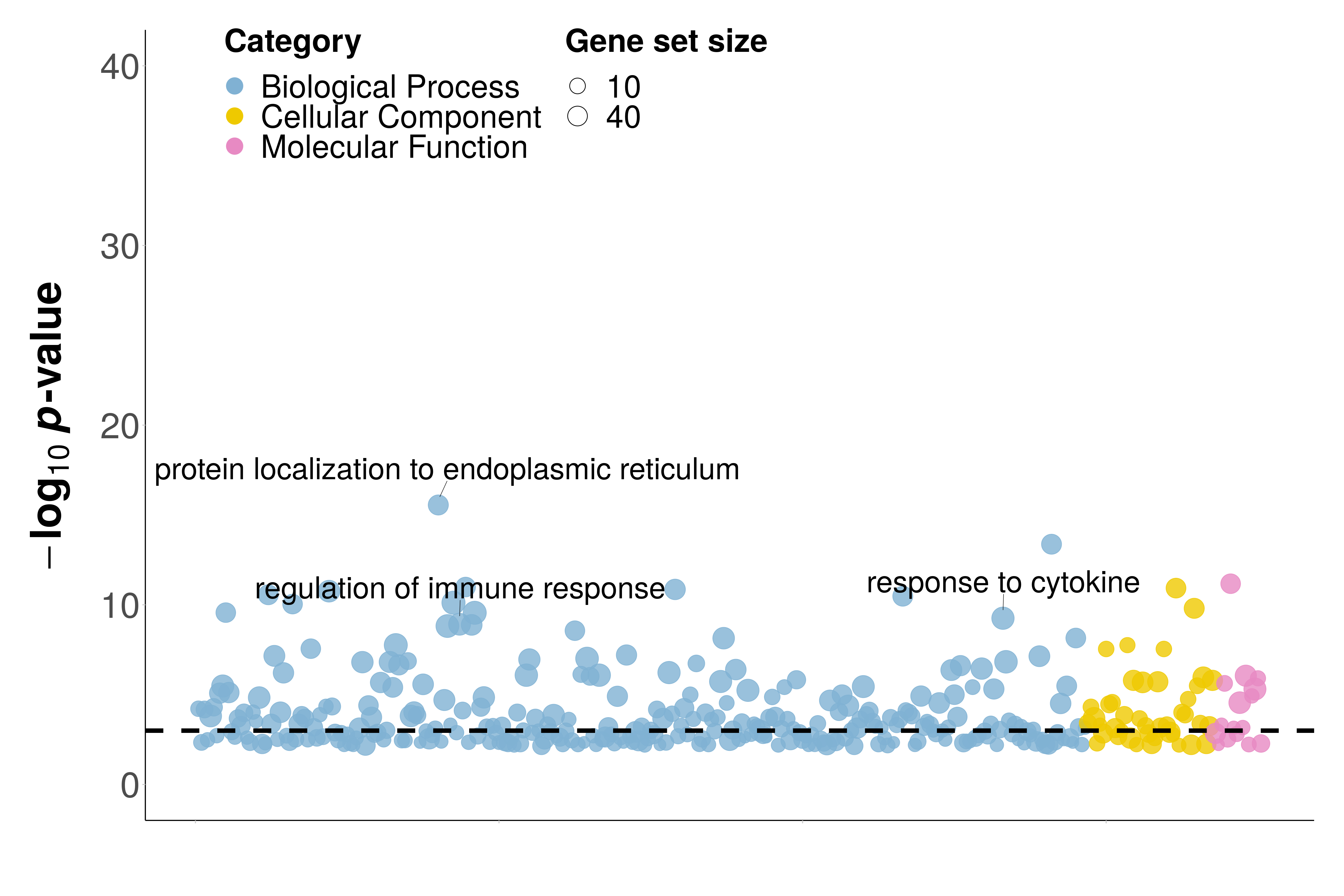

**A**

**B**

**C**

**Supplementary Figure 27: QQ Plot and Venn Diagram for mouse hypothalamus data. (A)** Quantile-quantile plot of the observed -log10 *p*-values from different methods against the expected -log10 *p*-values under the null in the real data analysis. Compared methods include SPARK (pink), SpatialDE (purple), Trendsceek.E (light salmon) which is the Emark test of Trendsceek, Trendsceek.$\rho$ (yellow-green) which is the Markcorr test of Trendsceek, Trendsceek.$\gamma$ (light green) which is the Markvario test of Trendsceek, and Trendsceek.V (wheat) which is the Vmark test of Trendsceek. *p*-value distribution suggests both SPARK and SpatialDE are more powerful than various Trendsceek tests in detecting genes with spatial expression pattern. **(B)** Venn diagram shows the overlap of the significant genes detected by SPARK, SpatialDE and Trendsceek the mouse hypothalamus data. SE genes are determined based on the p-value threshold that only one blank control gene is also detected.

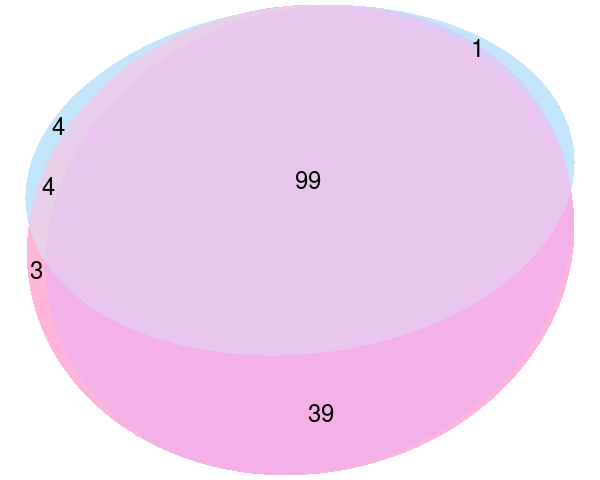

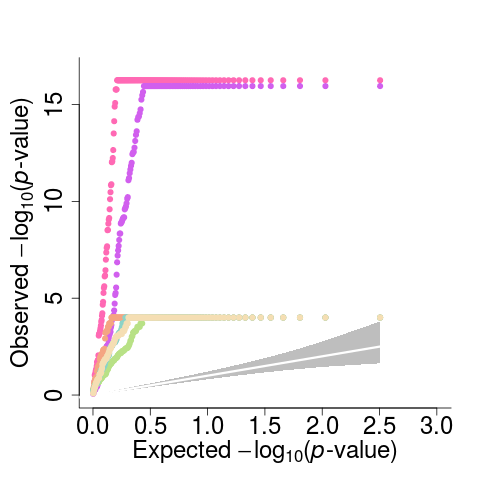

**A**

**B**

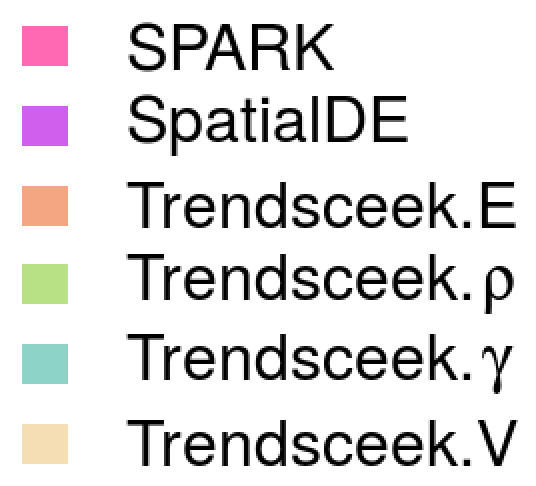

**Supplementary Figure 28: Spatial distributions of five individual cell classes with corresponding representative genes highlighted in the original study.** (**A**) Cells in each of the five cell classes are shown as colored dots (legends below), while the remaining cells are shown as gray dots. (**B**) Spatial expression pattern of five representative genes (*Slc17a6*, *Pdgfra*, *Fn1,* *Selplg, Aqp4*) for each of the five cells classes are shown. These five genes are identified as SE by both SPARK and SpatialDE. The *p*-values for the five genes from SPARK are shown inside parenthesis. Color represents relative gene expression level (purple: high; green: low).

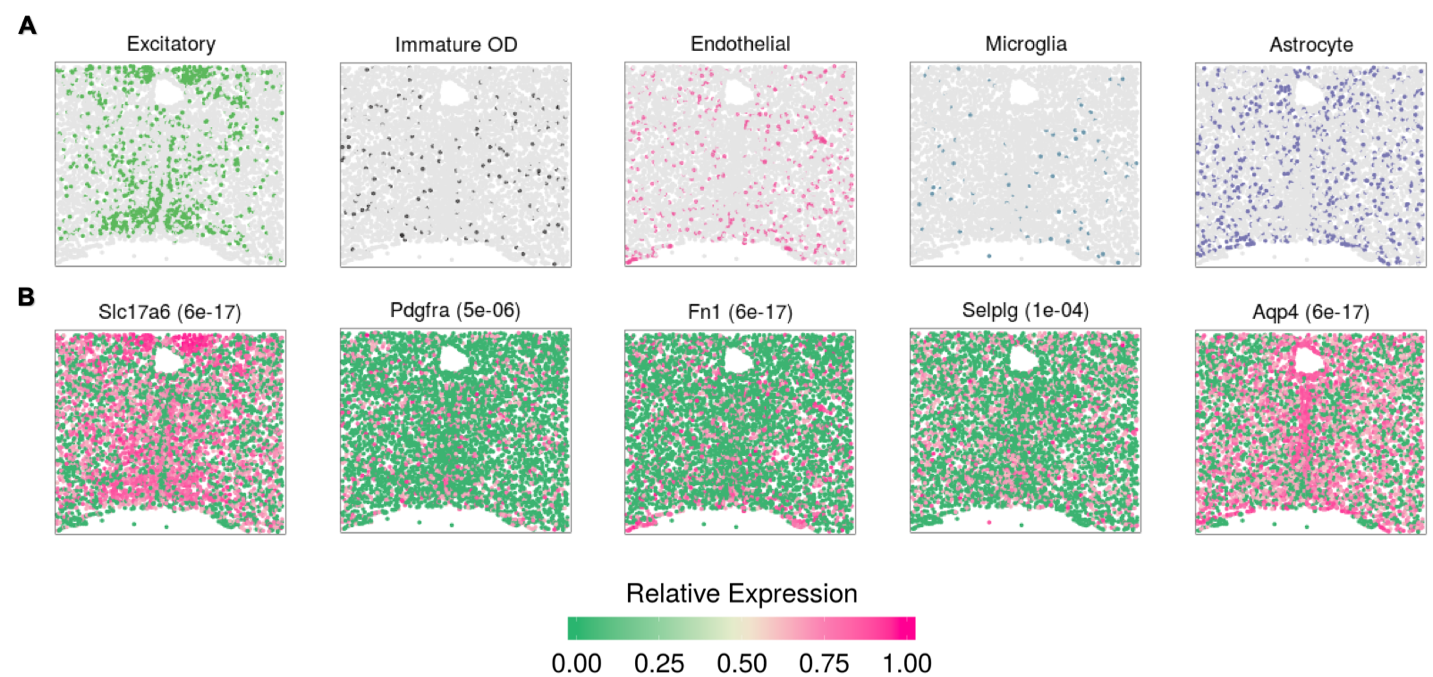

**Supplementary Figure 29: Analyzing the mouse hippocampus data**. (**A**) Voronoi representative of the tissue structure in the mouse hippocampus data. Red dashed rectangle represents the area that is adjusted for border artifacts. (**B**) Quantile-quantile plot of the observed -log10 *p*-values from different methods are plotted against the expected -log10 *p*-values under the null in the permuted data. *p*-values are combined across ten permutation replicates. Compared methods include SPARK (pink), SpatialDE (purple), Trendsceek.E (light salmon) which is the Emark test of Trendsceek, Trendsceek.$\rho$ (yellow-green) which is the Markcorr test of Trendsceek, Trendsceek.$\gamma$ (light green) which is the Markvario test of Trendsceek, and Trendsceek.V (wheat) which is the Vmark test of Trendsceek. Both SPARK Trendsceek test statistics produce reasonably calibrated *p*-values while SpatialDE yield slightly conservative *p*-values. (**C**) Quantile-quantile plot of the observed -log10 *p*-values from different methods against the expected -log10 *p*-values under the null in the real data analysis. *p*-value distribution suggests that SPARK detected more genes with spatial expression pattern than SpatialDE. In contrast, *p*-values from various Trendsceek tests all behave approximately like null and suggest a lack of power by Trendsceek tests. (**D**) Power plot shows the number of genes with spatial expression pattern (y-axis) identified by different methods at a range of false discovery rates (FDRs; x-axis). Across a range of FDRs, SPARK detected more genes with spatial expression pattern than SpatialDE and Trendsceek (sky-blue, which is combined test of Trendsceek).

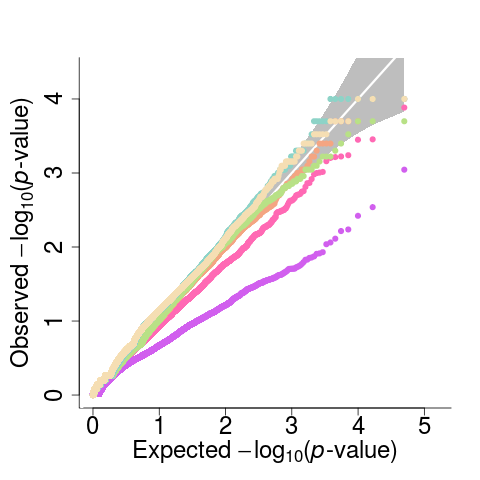

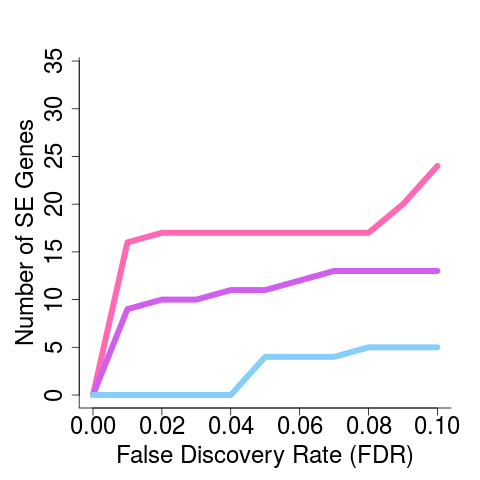

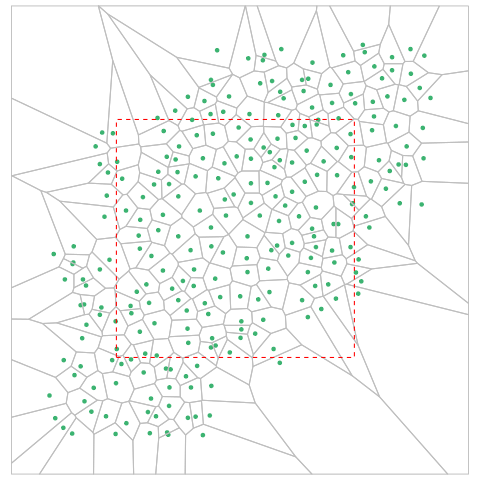

**A**

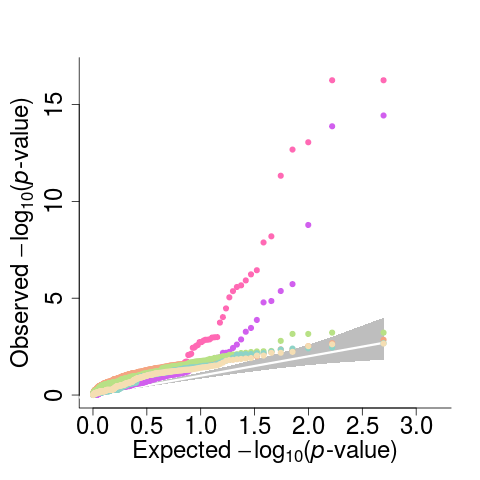

**B**

**C**

**D**

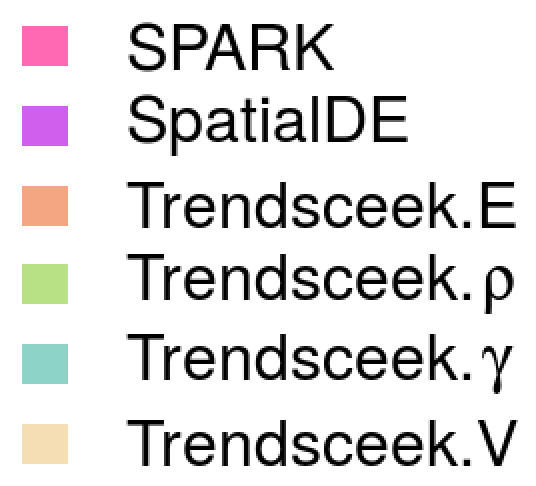

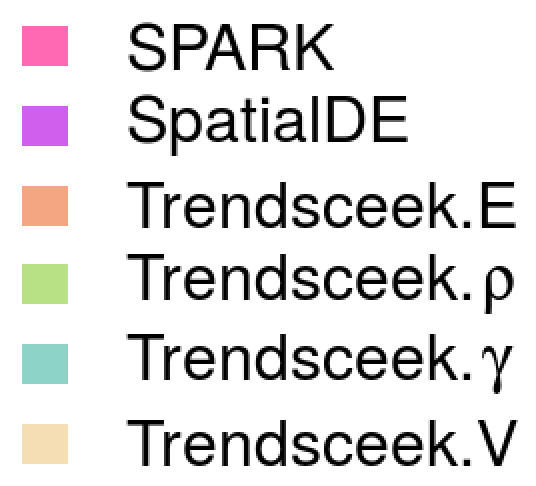

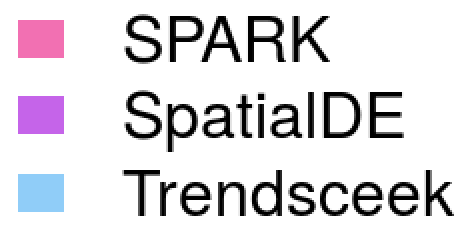

**Supplementary Figure 30: Venn diagram shows the overlap of significant genes detected by SPARK, SpatialDE and Trendsceek in the mouse hippocampus data.** Genes are detected based on an FDR cutoff of 0.05.

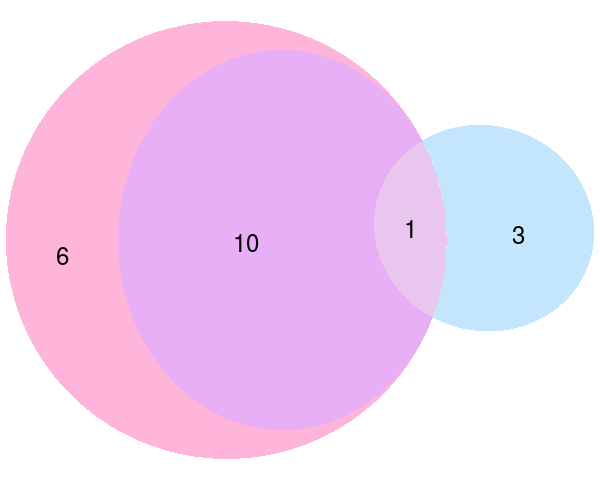

**Supplementary Figure 31: Spatial expression patterns for eleven SE genes identified by both SPARK and SpatialDE in mouse hippocampus data.** Genes are identified based on an FDR cutoff of 0.05. Color represents relative gene expression level (purple: high; green: low). SPARK *p*-values are shown in parenthesis. Among these genes, only *Ctss* gene was also identified by Trendsceek.

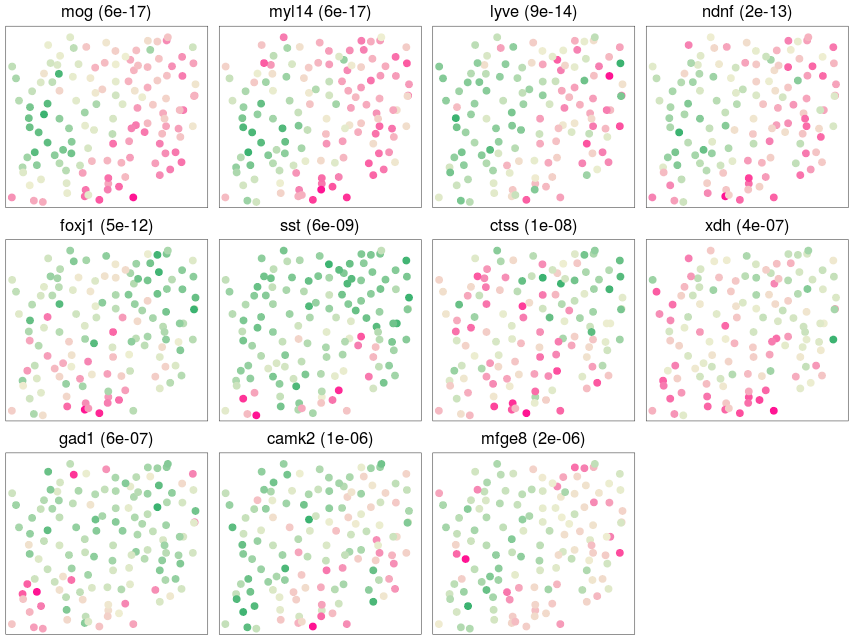

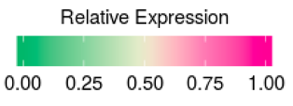

**Supplementary Figure 32: Spatial expression patterns for six SE genes identified only by SPARK in mouse hippocampus data.** Genes are identified based on an FDR cutoff of 0.05. Color represents relative gene expression level (purple: high; green: low). SPARK *p*-values are shown in parenthesis.

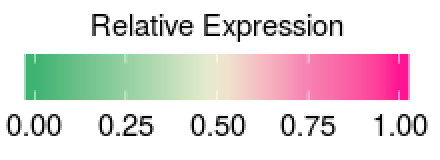

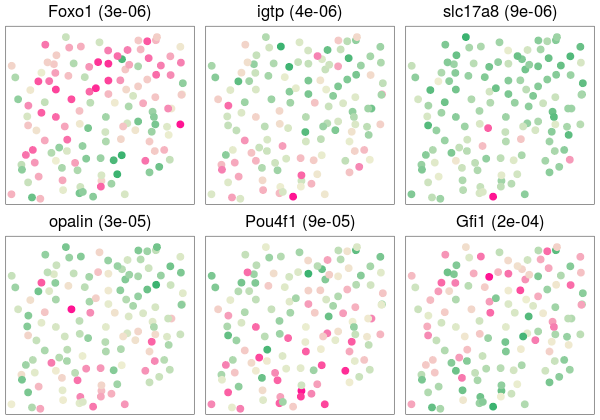

**Supplementary Figure 33: Spatial expression patterns for three SE genes identified only by Trendsceek in mouse hippocampus data.** Genes are identified based on an FDR cutoff of 0.05. Color represents relative gene expression level (purple: high; green: low).

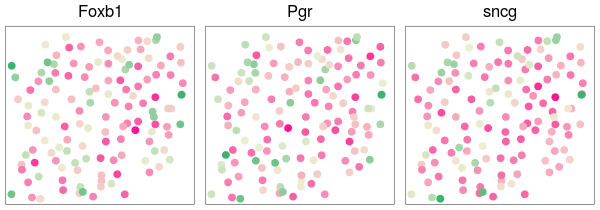

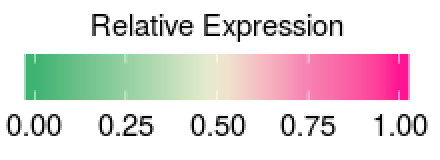

**Supplementary Figure 34: Quantile-Quantile plot of Moran’s I test on the permuted mouse olfactory data.** Quantile-quantile plot of the observed -log10 *p*-values from different methods against the expected -log10 *p*-values under the null in the permuted mouse olfactory bulb data. *p*-values are combined across ten permutation replicates. The Moran’s I test was performed using the *moran.test* from R package *spdep*.

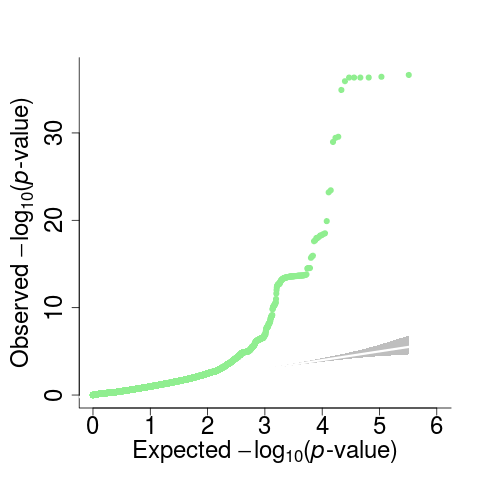

**Supplementary Figure 35: Venn diagrams show the overlap of the significant genes detected by SPARK and Moran’s I test.** Results are shown for (**A**) the mouse olfactory bulb data; (**B**) the mouse Hypothalamus data; and (**C**) the mouse hippocampus data. Moran’s I test did not detect any significant genes in the human breast cancer data, which is thus not shown here. Genes are detected based on an FDR cutoff of 0.05. The Moran’s I test was performed using the *moran.test* from R package *spdep*.

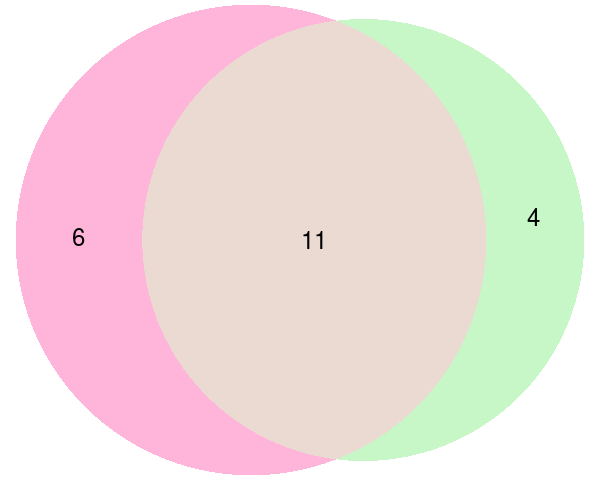

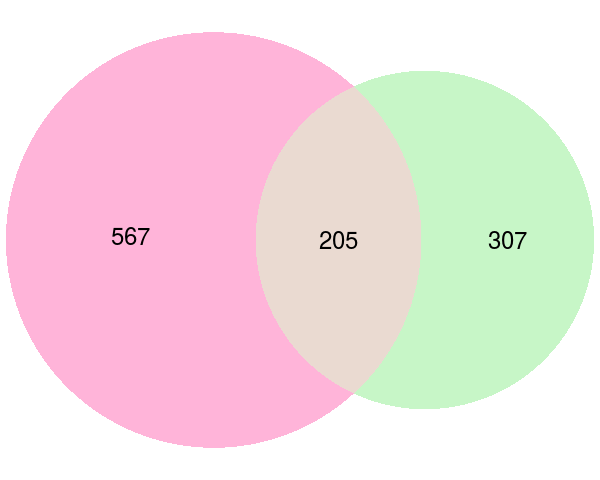

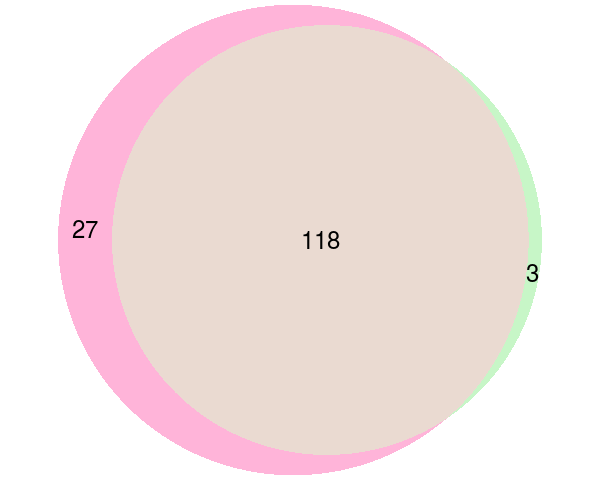

**A**

**B**

**C**

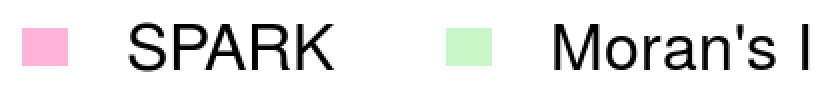

**Supplementary Figure 36: Mean and variance plots for four real datasets**. Each dot represents a gene, whose mean (x-axis) and variance (y-axis) were calculated. Loess curve is fitted for each dataset (red line). A clear mean-variance dependency pattern, with variance being larger than the mean, is clearly visualizable for all four data sets, regardless whether they are sequencing based or smFISH based. In each panel, the diagonal dashed line is where the variance equals the mean.

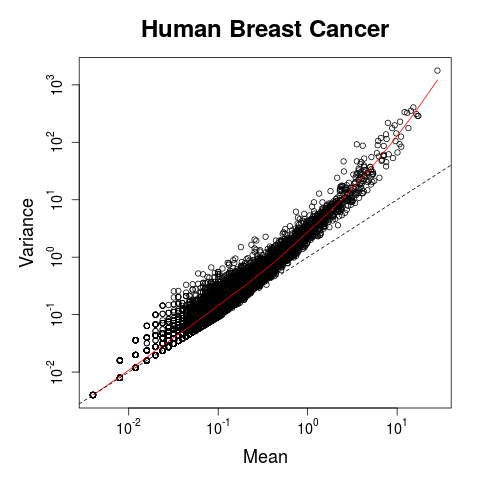

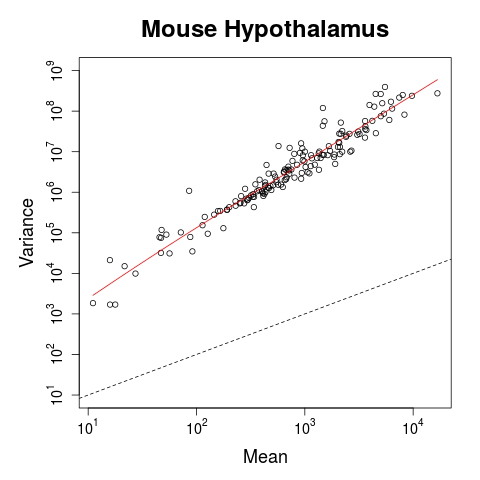

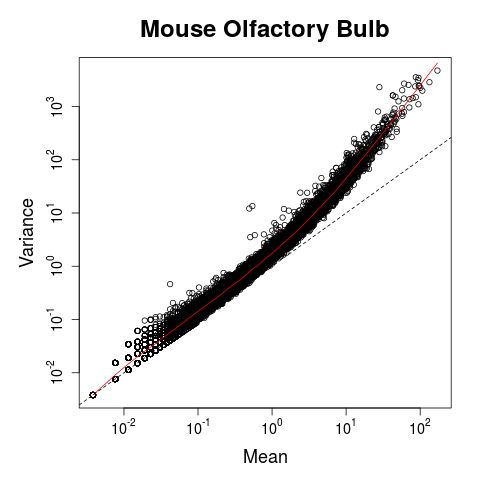

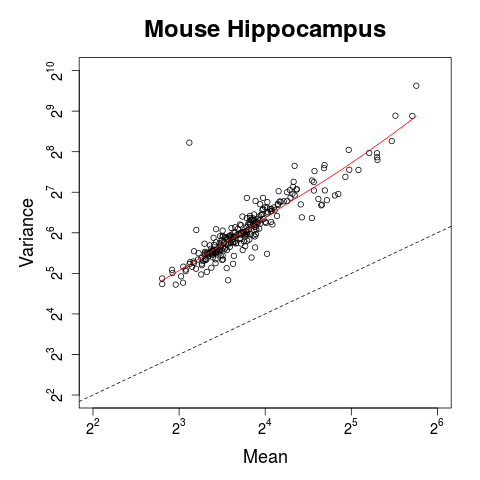

**D**

**A**

**C**

**B**

**Supplementary Figure 37: Quantile-quantile plot for *p*-values obtained from the Gaussian version of SPARK.** Quantile-quantile plot of the observed -log10 *p*-values from gaussian version of SPARK are plotted against the expected -log10 *p*-values under the null either for the real data (steel blue) or for the permuted data (light pink). Results are shown for (**A**) the mouse olfactory bulb data; (**B**) the human breast cancer data; (**C**) the mouse hypothalamus data; and (**D**) the mouse hippocampus data.

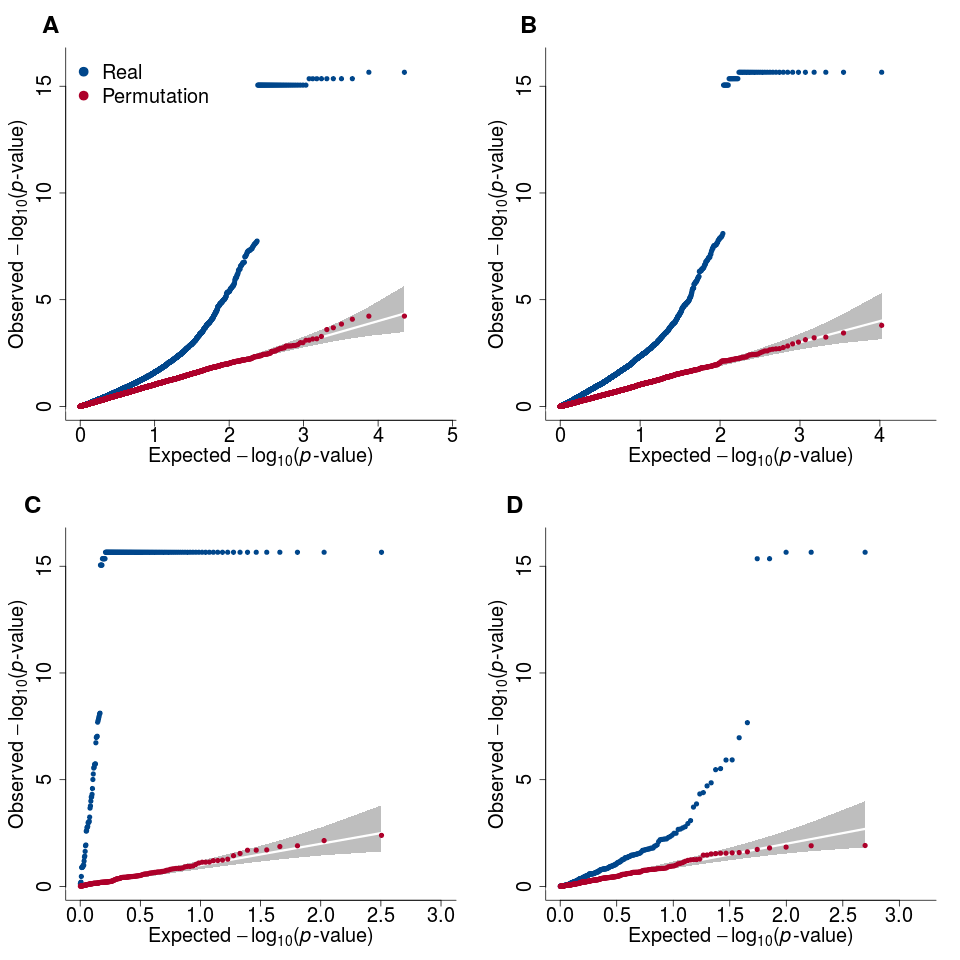

**Supplementary Figure 38: Venn diagram shows the overlap of significant genes detected by (the Poisson version of) SPARK and the Gaussian version of SPARK in all four real data sets.** Genes are detected based on an FDR cutoff of 0.05. Results are shown for **(A)** the mouse olfactory bulb data; **(B)** the human breast cancer data; **(C)** the mouse hypothalamus data; and **(D)** the mouse hippocampus data.

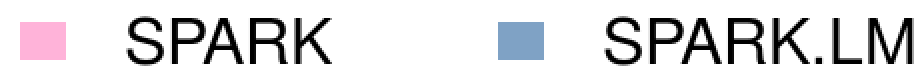

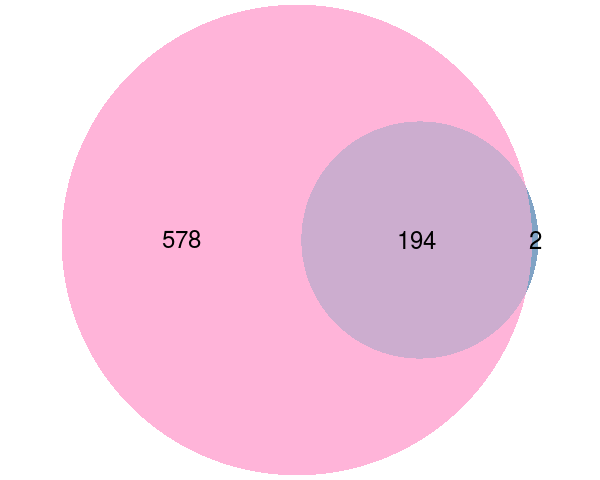

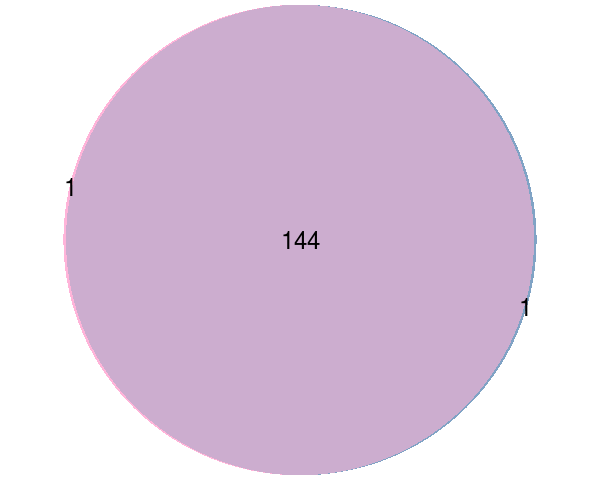

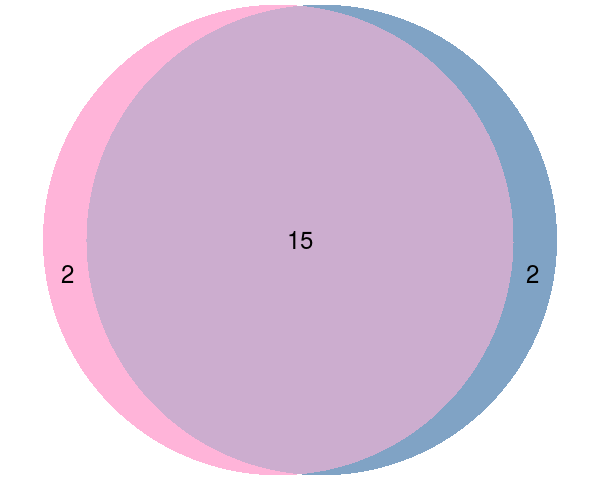

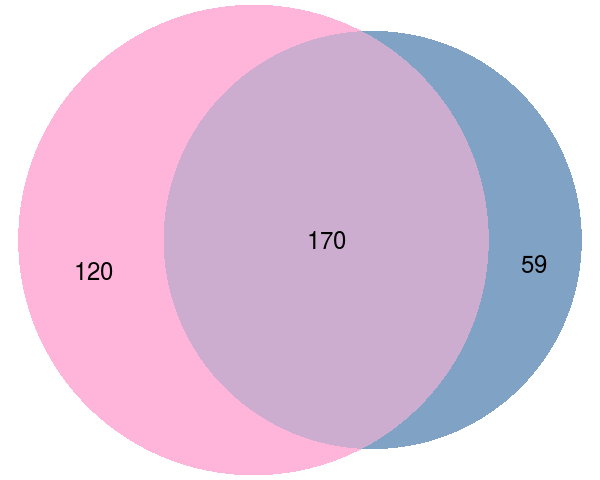

**B**

**A**

**D**

**C**

**Supplementary Figure 39: Quantile-quantile plot for *p*-values obtained separately from each of the ten kernels of SPARK.** The observed -log10 *p*-values from each kernel of SPARK are plotted against the expected -log10 *p*-values under the null for the real data sets. Results are for **(A)** the mouse olfactory bulb data; **(B)** the human breast cancer data; **(C)** the mouse hypothalamus data; **(D)** the mouse hippocampus data. The performance of SPARK is either identical to (**A-C**) or close to (**D**) the best kernel one could select.

**Supplementary Figure 40: Computational time of different methods for analyzing data with different sample sizes in the simulations**. Plot shows computational time in minutes (y-axis; on log10-scale) for analyzing 100 genes with different sample sizes (x-axis) for different methods. Compared methods include SpatialDE (purple), Trendsceek (blue), SPARK (pink), SPARK with 10 threads (dotted pink line). Computation are carried out using Intel Xeon E5-2683 2.00GHz processors. Both SPARK and SpatialDE are much more computationally efficient than Trendsceek.

**Supplementary Figure 41: Heatmaps illustrates five Gaussian kernels and five cosine kernels used in the mouse olfactory bulb data.** Five Gaussian kernels (GSP1-GSP5) are used to capture a range of possible focal expression correlation patterns while five cosine kernels (COS1-COS5) are used to capture a range of periodic expression patterns.

**Supplementary Figure 42: Spatial expression patterns summarized based on the 772 SE genes that are identified by SPARK.** (**A**) Clustering patterns obtained through hierarchical agglomerative clustering when the number of clusters is set to be five. (**B**) Clustering patterns obtained through hierarchical agglomerative clustering when the number of clusters is set to be three. The three main patterns summarized in (**B**) appear to be a subset of the patterns summarized in (**A**).

**A**

**B**

**Supplementary Table 3: Computation time in minutes for analyzing the four real datasets using different methods.** Computation are carried out using Intel Xeon E5-2683 2.00GHz processors. Computation time listed is for analyzing all genes for SPARK or SpatialDE but is for analyzing up to 500 most highly variable genes for Trendsceek.

| **Data Set** | **# of Samples/Genes** | **SPARK** | **SPARK*** | **SpatialDE** | **Trendsceek** |
| --- | --- | --- | --- | --- | --- |
| **Mouse Olfactory Bulb** | 260/11,274 | 46.53 | 5.62 | 6.98 | 80.88 |
| **Human Breast Cancer** | 250/5,262 | 25.21 | 3.74 | 5.91 | 426.50 |
| **Mouse Hypothalamus** | 4,975/160 | 62.80 | 15.8 | 16.7 | >7 days |
| **Mouse Hippocampus** | 131/249 | 0.69 | 0.28 | 0.10 | 990.98 |

*: SPARK with 10 threads; the other three columns list time when only 1 thread is used.

**Supplementary Text**

1.1 **Covariance Kernel Functions**

Following the previous literatures^1^^,^ ^2^, we considered two types of spatial kernel functions in SPARK:

(1) Gaussian kernel function (a.k.a. squared exponential kernel function or radial basis kernel function) in the form of

$$K_{G}(\boldsymbol{s}_{i}\boldsymbol{,}\boldsymbol{s}_{j})=\exp\left( -\frac{{\boldsymbol{\|}\boldsymbol{s}_{i}\boldsymbol{-}\boldsymbol{s}_{j}\boldsymbol{\|}}^{2}}{2\sigma^{2}} \right),$$

where $\left\| \boldsymbol{s}_{i}\boldsymbol{-}\boldsymbol{s}_{j} \right\|\boldsymbol{=}\sqrt{\left( \boldsymbol{s}_{i1}\boldsymbol{-}\boldsymbol{s}_{j1} \right)^{\boldsymbol{2}}\boldsymbol{+}\left( \boldsymbol{s}_{i2}\boldsymbol{-}\boldsymbol{s}_{j2} \right)^{\boldsymbol{2}}}$ is the Euclidean distance; and $\sigma$ is the length scale parameter that effectively characterizes the size of the focal expression patterns. The Gaussian kernel function is infinitely differentiable.

(2) Cosine kernel function which is a periodic kernel function in the form of

$$K_{C}\left( \boldsymbol{s}_{i}\boldsymbol{,}\boldsymbol{s}_{j} \right)=\cos\left( 2\pi\frac{\left\| \boldsymbol{s}_{i}\boldsymbol{-}\boldsymbol{s}_{j} \right\|}{\phi} \right),$$

where $\phi$ is the periodicity parameter that effectively characterizes the frequency of the periodic expression pattern.

The choice of the length scale parameter ($\sigma$) in the Gaussian kernel and the choice of the periodicity parameter ($\phi$) in the cosine kernel are important for constructing powerful hypothesis test. As default, SPARK uses five Gaussian kernels with pre-determined $\sigma$ values together with five cosine kernels with pre-determined $\phi$ values. The choice of $\sigma$ and $\phi$ for these kernels is pre-determined based on the pair-wise Euclidean distances among spatial locations in the data to ensure scale-invariance. In particular, we first obtained the minimum ($m_{1}$) and the maximum ($m_{2}$) value of the non-zero Euclidean distances across all pairs of spatial locations. We then extracted ten equally spaced values ranging from $log10(m_{1}/2 )$ to $log10\left( m_{2}*2 \right)$ and kept the five values in the middle of the list (i.e. the 3^rd^ to 7^th^ values). Afterwards, we converted the obtained five values back to the original Euclidean distance scale by taking power of ten and used the converted values to serve as the five $\sigma$’s and the five $\phi$’s.

1.2 **Parameter Estimation in the Null Model**

Here, we describe our inference algorithm for parameter estimation in the null GSLM model. These parameter estimates from the null model are used to construct score test statistics. For simplicity of the presentation, we will ignore the spatial location notation for all variables in the model in the following text (e.g. replacing $y_{i}(\boldsymbol{s}_{\boldsymbol{i}})$ with $y_{i}$). Our inference algorithm is based on the penalized quasi-likelihood (PQL) algorithm^3^ and employs an iterative numerical optimization procedure. In each iteration, we introduce a set of continuous pseudo-data $\tilde{\boldsymbol{y}}$ to replace the originally observed count data $\boldsymbol{y}$. The pseudo-data $\tilde{\boldsymbol{y}}$ is obtained based on a second order Taylor expansion using the conditional distribution $P\left( \left. y_{i} \right|\boldsymbol{b, \epsilon} \right)$ with the first and second order moments $E\left( \left. y_{i} \right|\boldsymbol{b, \epsilon} \right)$ and $V\left( \left. y_{i} \right|\boldsymbol{b, \epsilon} \right)$, both evaluated at the current estimates of the fixed coefficients $\boldsymbol{\beta}$ as well as the current estimates of the spatial effects $\boldsymbol{b}$ and the nugget effects $\boldsymbol{\epsilon}$. With the pseudo-data, the complex GLSM likelihood function for the original count data $\boldsymbol{y}$ is replaced by a much simpler spatial linear mixed model likelihood function for the continuous pseudo-data$\tilde{\boldsymbol{y}}$, thereby alleviating much of the computational burden associated with GLSM. With pseudo-data $\tilde{\boldsymbol{y}}$, we can perform estimation and update parameters using the standard average information (AI) algorithm commonly used in linear mixed models^4^^,^ ^5^. By iterating between the approximation step of obtaining the pseudo-data $\tilde{\boldsymbol{y}}$ and the inference step of updating the parameter estimates via the AI algorithm, SPARK allows us to perform inference in a computationally efficient fashion.

We focus on the null model under the null hypothesis ${H_{0}:\tau}_{1}=0$. In this case, the SPARK model is reduced to:

$y_{i}\sim Poi\left( {N_{i}\lambda}_{\boldsymbol{i}} \right),$

$$\log\left( \lambda_{i} \right)=\eta_{i}=\boldsymbol{x}_{i}^{T}\boldsymbol{\beta}+\epsilon_{i}, \epsilon_{i}\sim N\left( 0, \tau_{2} \right).$$

Our goal is to estimate $\boldsymbol{\beta}$ and $\tau_{2}$. To do so, we note that the observations $y_{i}$ are independent conditional on the unobserved nugget effects $\boldsymbol{\epsilon}$ and the fixed effects $\boldsymbol{x}_{i}\boldsymbol{\beta}$, with conditional mean $E\left( \left. y_{i} \right|\boldsymbol{\beta,\epsilon} \right)=\mu_{i}=g^{-1}(\boldsymbol{x}_{i}\boldsymbol{\beta}+\epsilon_{i})$ and conditional variance $V\left( y_{i}\boldsymbol{|\beta},\boldsymbol{\epsilon} \right)=\mu_{i}$, where $g\left( \cdot\right)$ is the log link function. We use these two conditional moments to obtain the quasi-likelihood for$i$-th sample, ${ql}_{i}\left( \left. \boldsymbol{\beta} \right|\tilde{\boldsymbol{\epsilon}} \right)=\int_{y_{i}}^{\mu_{i}} \frac{y_{i}-t}{t}dt$, which serves as an approximation for the conditional likelihood. The joint likelihood function can thus be approximated by the joint quasi-likelihood function

$$ql\left( \boldsymbol{\beta},\tau_{2} \right)=log\int\left( \prod_{i=1}^{n} ql_{i}\left( \boldsymbol{\epsilon,\beta} \right) \right)P\left( \boldsymbol{\epsilon|}\tau_{2} \right)d\boldsymbol{\epsilon}.$$

We use Laplace approximation on $\boldsymbol{\epsilon}$ to further approximate the above function and obtain

$\tilde{ql}\left( \boldsymbol{\beta},\tau_{2} \right)=\frac{1}{2}\log\left| \mathbf{VD}+\mathbf{I} \right|+\sum_{i=1}^{n} {ql}_{i}\left( \tilde{\boldsymbol{\epsilon}},\boldsymbol{\beta} \right)-\frac{1}{2}{\tilde{\boldsymbol{\epsilon}}}^{T}\mathbf{V}^{-1}\tilde{\boldsymbol{\epsilon}},$	(1)

where$\mathbf{V}=\tau_{2}\mathbf{I}$,$\tilde{\boldsymbol{\epsilon}}=\underset{\boldsymbol{\epsilon}}{\mathrm{argmax}} \left( \sum_{i=1}^{n} {ql}_{i}\left( \left. \boldsymbol{\beta} \right|\boldsymbol{\epsilon} \right)-\frac{1}{2}\boldsymbol{\epsilon}^{T}\mathbf{V}^{-1}\boldsymbol{\epsilon} \right)$ and $\mathbf{D}=diag\left( 1/{g'\left( \mu_{i} \right)} \right)$ is a diagonal weight matrix with diagonal elements being the inverse of the first order derivative of $g\left( \mu_{i} \right)$.

We treat the approximated quasi-likelihood function $\tilde{ql}\left( \boldsymbol{\beta},\tau_{2} \right)$ in equation (1) as the target function. And we obtain estimates for $\boldsymbol{\beta}$ and $\tau_{2}$ alternately from equation (1). Specifically, we first obtain estimates for $\boldsymbol{\beta}$conditional on the current estimates of $\tau_{2}$. To do so, following^3^^,^ ^6^, we assume that the iterative weights vary slowly with respect to the conditional mean; that is

$$\frac{\partial\mathbf{D}}{\partial\mu_{i}}\approx0.$$

We then obtain the first order derivatives with respect to either $\boldsymbol{\beta}$ or $\boldsymbol{\epsilon}$, and set the two first order derivatives to zero; that is

$\mathbf{X}^{T}\boldsymbol{D\Delta}\left( \boldsymbol{y-\mu} \right)\boldsymbol{=0,}$	(2)

$\boldsymbol{D\Delta}\left( \boldsymbol{y-\mu} \right)\boldsymbol{-}\mathbf{V}^{\boldsymbol{-1}}\boldsymbol{\epsilon=0,}$	(3)

where $\mathbf{X}$ is an *n* by *k* covariate matrix obtained by stacking $\boldsymbol{x}_{i}$; $\boldsymbol{\mu}=\left( \mu_{1}, \cdots,\mu_{n} \right)^{T}$ and $\boldsymbol{\Delta=}\mathrm{diag}\left\{ g'\left( \mu_{i} \right) \right\}$.

We now define the pseudo-data

$\tilde{y_{i}}=\eta_{i}+g^{'}\left( \mu_{i} \right)\left( y_{i}-\mu_{i} \right),$	(4)

and our equation (2) becomes

$\boldsymbol{\epsilon=V}\mathbf{H}^{-1}\left[ \tilde{\mathbf{y}}\boldsymbol{-}\mathbf{X}\boldsymbol{\beta} \right]\boldsymbol{,}$	(5)

where $\mathbf{H=}\mathbf{D}^{-1}\mathbf{+V}$ is a diagonal matrix. Substituting equation (5) into equation (3), we can obtain the estimates

 $\hat{\boldsymbol{\beta}}\boldsymbol{=}\left[ \mathbf{X}^{\mathbf{T}}\mathbf{H}^{-1}\mathbf{X} \right]^{-1}\mathbf{X}^{T}\mathbf{H}^{-1}\tilde{\mathbf{y}}\boldsymbol{,}$	(6)

and

$\hat{\boldsymbol{\epsilon}}\mathbf{=V}\mathbf{H}^{-1}\left[ \tilde{\mathbf{y}}\boldsymbol{-}\mathbf{X}\hat{\boldsymbol{\beta}} \right]$**.** (7)

Both of the above two estimates are conditional on the nugget estimate $\tau_{2}$.

Next, we obtain estimates for the variance component $\tau_{2}$conditional on the current estimates of $(\boldsymbol{\epsilon}, \boldsymbol{\beta)}$. To do so, we integrate out the fix effects $\boldsymbol{\beta}$ in equation (1) to obtain the restricted likelihood function as

$$\tilde{ql}_{R}\left( \tau_{2} \right)=c_{R}-\frac{1}{2}\log\left| \mathbf{H} \right|-\frac{1}{2}\log\left| \mathbf{X}^{T}\mathbf{H}^{-1}\mathbf{X} \right|-\frac{1}{2}{\tilde{\mathbf{y}}}^{T}\mathbf{P}\tilde{\mathbf{y}},$$

where $\mathbf{P}= \mathbf{H}^{-1}-\mathbf{H}^{-1}\mathbf{X}^{T}\left( \mathbf{X}^{T}\mathbf{H}^{-1}\mathbf{X} \right)^{-1}\mathbf{X}\mathbf{H}^{-1}$, and $c_{R}$ is a constant. We use the AI algorithm to obtain variance component estimate. In particular, we obtain the first derivatives as

$$\frac{\partial\tilde{ql}_{R}\left( \tau_{2} \right)}{\partial\tau_{2}}=\frac{1}{2}\left\{ {\tilde{\mathbf{y}}}^{T}\mathbf{PP}\tilde{\mathbf{y}}-tr\left( \mathbf{P} \right) \right\},$$

and the second derivatives as

$$\frac{\partial^{2}\tilde{ql}_{R}\left( \tau_{2} \right)}{\partial\tau_{2}^{2}}=\frac{1}{2}tr\left( \mathbf{PP} \right)-{\tilde{\mathbf{y}}}^{T}\mathbf{PPP}\tilde{\mathbf{y}}$$

Because the elements in the expected information matrix are

$$E\left[ \frac{\partial^{2}\tilde{ql}_{R}\left( \sigma^{2},h^{2} \right)}{\partial\tau_{2}\partial\tau_{2}} \right]=-\frac{1}{2}tr\left( \mathbf{PP} \right),$$

we can obtain the average information (AI) as an average of the above quantity; that is

$$\mathrm{AI}\mathbf{=-}{\tilde{\mathbf{y}}}^{T}\mathbf{PPP}\tilde{\mathbf{y}}.$$

With the first and second order derivatives, we can perform Newton-Raphson update with the AI algorithm and obtain estimates for $\tau_{2}$.

As a summary, SPARK implements the PQL algorithm that consists of the following steps:

1. Initialize the parameters, $\boldsymbol{\beta}^{\left( 0 \right)}\boldsymbol{,} \tau_{2}^{\left( 0 \right)},$ and obtain the pseudo-data ${\tilde{\mathbf{y}}}^{\left( 0 \right)}$ as in equation (4). Set $t=1$.
2. Update $\tau_{2}^{\left( t \right)}\boldsymbol{=}\tau_{2}^{\left( t-1 \right)}\boldsymbol{+}{AI}^{-1}\left( \frac{\partial{ql}_{R}\left( \tau_{2} \right)}{\partial\tau_{2}} \right)$;
3. Update $\boldsymbol{\beta}^{\left( t \right)}$and $\boldsymbol{\epsilon}^{\left( t \right)}$ with $\tau_{2}^{\left( t \right)}$ and ${\tilde{\mathbf{y}}}^{\left( t-1 \right)}$ as in equations (6) and (7);
4. Update ${\tilde{\mathbf{y}}}^{\left( t \right)}$ using the $\boldsymbol{\beta}^{\left( t \right)}$and $\boldsymbol{\epsilon}^{\left( t \right)}$ as in equation (4);
5. Set $t=t+1$, and repeat steps 2-4 until convergence.
   1.3 **Gaussian Version of SPARK**

Besides the count-based model, we also developed a Gaussian version of SPARK for modeling normalized spatial data. We again examine one gene at a time. The observed expression measurements for the *i*th cell, $y_{i}\left( \boldsymbol{s}_{i} \right)$, is no longer count data but normalized data. We can obtain such normalized data through, for example, log transformation of the original count. We assume the normalized expression data follows a normal distribution, i.e.,

$y_{i}\left( \boldsymbol{s}_{i} \right) =\boldsymbol{x}_{i}\left( \boldsymbol{s}_{i} \right)^{\boldsymbol{T}}\boldsymbol{\beta}+b_{i}\left( \boldsymbol{s}_{i} \right)+\epsilon_{i}$, $i=1,2,\cdots,n$

where $\boldsymbol{\beta}$ again is a *k*-vector of coefficients that include an intercept representing the mean log-expression of the gene across spatial locations together with *k*-1 coefficients for the corresponding explanatory variables; $\epsilon_{i}$ is the residual error that is independently and identically distributed from $N(0,\tau_{2} )$ with variance $\tau_{2}$; and $b_{i}(\boldsymbol{s}_{i})$ is a zero-mean, stationary Gaussian process modeling the spatial correlation pattern among spatial locations

$$\boldsymbol{b}\left( \boldsymbol{s}_{i} \right)={(b_{1}(\boldsymbol{s}_{1}),b_{2}(\boldsymbol{s}_{2}),\cdots,b_{n}(\boldsymbol{s}_{n}))}^{T}\sim MVN\left( \mathbf{0},\tau_{1}\boldsymbol{K}(\boldsymbol{s}) \right),$$

where the covariance $\boldsymbol{K}(\boldsymbol{s})$ is a kernel function of the spatial locations$\boldsymbol{s}=\left( \boldsymbol{s}_{1},\cdots,\boldsymbol{s}_{n} \right)^{T}$, with *ij*’th element being$\boldsymbol{K}(\boldsymbol{s}_{i},\boldsymbol{s}_{j})$; $\tau_{1}$ is a scaling factor of the covariance kernel; and MVN denotes a multivariate normal distribution. In the above model, the covariance for the normalized gene expression $y_{i}\left( \boldsymbol{s}_{i} \right)$ is$\boldsymbol{\Sigma}=\tau_{1}\boldsymbol{K}\left( \boldsymbol{s} \right)+\tau_{2}\boldsymbol{I}$, where $\boldsymbol{I}$ is an *n*-dimensional identity matrix. We are again interested in testing the null hypothesis ${\boldsymbol{H}_{\mathbf{0}}:\tau}_{1}=0$ and we do so by using the score statistics which is computed in the following form

$$\boldsymbol{Q}\left( \boldsymbol{y} \right)={\tilde{\boldsymbol{y}}\left( \boldsymbol{s} \right)}^{\boldsymbol{T}}\boldsymbol{SKS}\tilde{\boldsymbol{y}}\left( \boldsymbol{s} \right),$$

where $\boldsymbol{S=I-X}\left( \boldsymbol{X}^{\boldsymbol{T}}\boldsymbol{X} \right)^{\boldsymbol{-1}}\boldsymbol{X}$ is the projection matrix to the subspace orthogonal to the covariates $\boldsymbol{X}$**.** The score statistic follows a mixture of chi-square distributions under the null^7^^,^ ^8^

$${\boldsymbol{Q}\left( \boldsymbol{y} \right)}/{\tau_{2}} \sim\sum_{i=1}^{t} \phi_{i}\chi_{1i}^{2},$$

where $\left\{ \phi_{i} \right\}$ is the eigenvalues of $\boldsymbol{SKS}$; $\chi_{1i}^{2}$ is a random variable independently and identically distributed from a chi-square distribution with the degree of freedom being one; and $t$ is the number of non-zero eigen values. After obtaining the p-value for each kernel, we again combined them together though the Cauchy combination rule.

Compared with the Poisson version of SPARK, the Gaussian version of SPARK is computationally much more efficient. In addition, the Gaussian version of SPARK may be more robust to model misspecification than the Poisson version of SPARK, thus could be more effective in certain data applications. We have implemented both the Poisson version and Gaussian version of SPARK in the software package.

1.4 **Score Test and the Cauchy Combination Rule**

For each of the ten kernel functions described in section 1.1, we obtained parameters using the PQL algorithm described in section 1.2. Afterwards, we construct a score statistics

$$S_{0}\equiv\frac{1}{2} {\tilde{\boldsymbol{y}}}^{T}\boldsymbol{PKP}\tilde{\boldsymbol{y}}.$$

Because each $\tilde{y}_{i}$ follows a normal distribution $N(x_{i}^{T}\beta, g^{'}(\mu_{i})+\tau_{2})$ under the null, the score statistic $S_{0}$ follows a mixture of $\chi_{1}^{2}$ distributions $\sum_{i=1}^{n} \psi_{i}\chi_{1i}^{2}$ under the null, where $\psi_{i}$ are the eigenvalues of the matrix $\boldsymbol{PK}/2$ and are served as weights for the mixture distribution; while $\chi_{1i}^{2}$ denotes a random variable from the chi-square distribution with one degree of freedom. We compute *p*-value based on the above score test statistic using the Satterthwaite Method^9^. Specifically, we approximate the mixture chi-square distribution of $S_{0}$ by a scaled chi-square distribution ${\kappa\chi}_{\nu}^{2}$ through matching moments, where $\kappa$ is the scaled parameter and $\nu$ is the degrees of freedom. By matching the mean of the two distributions and matching the variance of the two distribution, we obtain

$$\hat{\kappa}=\frac{I_{\tau_{1}\tau_{1}}}{2tr\left( PK \right)}, \hat{\nu}=\frac{4e_{s}^{2}}{tr\left( PKPK \right)}.$$

With $\hat{\kappa}$ and $\hat{\nu}$, we obtained *p*-value for the score statistics based on *p*-value definition.

Through the above procedure, we obtained ten different *p*-values, one for each spatial kernel. Afterwards, we combined these ten *p*-values into a single *p*-value through the recently developed Cauchy combination rule^10^. Specifically, we converted each of the ten *p*-values into a Cauchy statistic, averaged the ten Cauchy statistics, and then converted the average back to a single *p*-value using the Cauchy distribution. The Cauchy rule takes advantage of the fact that summation of Cauchy random variables also follows a Cauchy distribution regardless whether these random variables are correlated or not^10^. Therefore, the Cauchy rule allows us to combine multiple potentially correlated *p*-values into a single *p*-value without loss of type I error control. Due to numerical precision of the Cauchy cumulative density function, the combined *p*-value is output as zero when it is below the threshold of 5.55e-17; in this case, we simply set the output *p*-value as this threshold. Finally, after obtaining *m* *p*-values across *m* genes, we controlled for false discovery rate (FDR) using the *Benjamini–Yekutieli* (BY) procedure, which is effective under arbitrary dependence across genes^11^.

1.5 **Compared Methods Overview**

Here, we provide a brief description of the two existing methods for identifying genes with spatial expression patterns. We use the same set of notations as we used for SPARK to describe these methods.

*Trendsceek* examines one gene at a time. For each gene, Trendsceek first obtains the normalized gene expression $y_{i}$ for *i*-th cell through log10 transformation. It then models the expression level $y_{i}$ and the spatial location of *i*-th cell $s_{i}$ jointly through a marked point process. The marked point process relies on the probability density $f(s_{i},y_{i})$ to describe the probability of observing a point $s_{i}$ with mark $y_{i}(s_{i})$. In addition, it relies on the joint probability density $f(\left( s_{i},y_{i} \right),\left( s_{j},y_{j} \right))$ to describe the joint probability of observing two points $s_{i}$ and $s_{j}$ with marks $y_{i}$ and $y_{j}$. Trendsceek computes the Euclidean distance between the two points as $r=|s_{j}-s_{i}|$ and obtains a conditional probability $M\left( y_{i},y_{j}|r \right)=f(\left( s_{i},y_{i} \right),\left( s_{j},y_{j} \right))/f(r)$. Trendsceek then examines the joint distribution of all pairs of points at a particular radius *r* and defines the concept of mark segregation when the distribution is dependent on *r*, such that it deviates from what would be expected if the marks were randomly distributed over the spatial locations of the points. Therefore, the null hypothesis of Trendsceek is the absence of mark segregation defined as $H_{0}:M\left( y_{i},y_{j}|r \right)=M\left( y_{i} \right)M(y_{i})$. To test for the absence of mark segregation, Trendsceek computes four different non-parametric test statistics that include Stoyan's mark-correlation function, the mean-mark function, the variance-mark function, and the mark-variogram of a marked point process. Trendsceek relies on permutations to obtain a p-value for each test statistics. In the permutations, it samples the mark distribution without replacement and randomly reassigned expression levels to all cells, keeping their positions fixed and in effect conditioning on the given spatial locations. Finally, Trendsceek obtains four different p-values from the four non-parametric statistics for each gene. In addition, genes with Benjamini–Hochberg adjusted P ≤ 0.05 for at least one of the four statistics tests are considered as significant.

*SpatialDE* also examines one gene at a time. For each gene, SpatialDE first normalizes the raw count data using a variance stabilizing transformation. It treats the transformed values as the outcome and the log total count values as the covariate and performs a linear regression to remove the effects of read depth. It obtains the residuals from the linear regression as the final outcome $y_{i}$ for spatial modeling. Afterwards, SpatialDE models the gene expression profiles across spatial coordinates using linear mixed models with Gaussian kernels. For each of the ten Gaussian kernels, SpatialDE obtains a likelihood ratio test statistics and relies on the asymptotic chi-square distribution with one degree of freedom to further obtain an approximate p-value. Among the resulting ten p-values from ten Gaussian kernels, SpatialDE selects the minimal p-value as the final output. SpatialDE determines an initial set of significant genes based on these minimal p-values. For each of these initial significant genes, SpatialDE performs an additional step of model selection by using various other covariance kernels for further modelling of periodic patterns and linear trends. The p-value from the kernel with the lowest Bayesian information criterion (BIC) is reported as the final p-value for the given gene. Finally, SpatialDE uses q-value method to adjust for multiple testing.

Trendsceek relies on non-parametric test statistics and thus does not have an explicit underlying data generative model. In contrast, both SpatialDE and SPARK have an underlying data generative model and relies on parametric test statistics to perform inference. However, there are several key differences between SpatialDE and SPARK: **(1)** SPARK directly models count data, while SpatialDE models normalized expression data; **(2)** SPARK relies on a set of Gaussian and periodic kernels for analysis, while SpatialDE only uses Gaussian kernels for the initial screening; **(3)** SPARK calculates an exact p-value based on the score test statistics using a mixture chi-square distributions, while SpatialDE computes an approximate p-value based on the likelihood test statistics using asymptotic normal/chi-square approximation; **(4)** SPARK combines p-values across different kernels using a novel Cauchy combination rule that guarantees the calibration of the resulting p-value, while SpatialDE obtains the minimal p-value among these p-values as the final output; **(5)** SpatialDE also performs an additional analysis on the initial set of significant genes obtain through screening and relies on additional kernels to obtain the final p-value for these genes.

**REFERENCE**

1.	Svensson, V., Teichmann, S.A. & Stegle, O. SpatialDE: identification of spatially variable genes. *Nat Methods* **15**, 343-346 (2018).

2.	Rasmussen, C.E. & Williams, C.K.I. Gaussian Processes for Machine Learning. (MIT, 2006).

3.	Breslow, N.E. & Clayton, D.G. Approximate Inference In Generalized Linear Mixed Models. *J Am Stat Assoc* **88**, 9-25 (1993).

4.	Chen, H. et al. Control for Population Structure and Relatedness for Binary Traits in Genetic Association Studies via Logistic Mixed Models. *The American Journal of Human Genetics* **98**, 653--666 (2016).

5.	Yang, J.A., Lee, S.H., Goddard, M.E. & Visscher, P.M. GCTA: A Tool for Genome-wide Complex Trait Analysis. *Am J Hum Genet* **88**, 76-82 (2011).

6.	Gilmour, A.R., Thompson, R. & Cullis, B.R. Average information REML: An efficient algorithm for variance parameter estimation in linear mixed models. *Biometrics* **51**, 1440-1450 (1995).

7.	Schweiger, R. et al. RL-SKAT: An Exact and Efficient Score Test for Heritability and Set Tests. *Genetics* **207**, 1275-1283 (2017).

8.	Wu, M.C. et al. Rare-Variant Association Testing for Sequencing Data with the Sequence Kernel Association Test. *Am J Hum Genet* **89**, 82-93 (2011).

9.	Satterthwaite, F.E. An Approximate Distribution Of Estimates Of Variance Components. *Biometrics Bull* **2**, 110-114 (1946).

10.	Liu, Y.W. et al. ACAT: A Fast and Powerful p Value Combination Method for Rare-Variant Analysis in Sequencing Studies. *Am J Hum Genet* **104**, 410-421 (2019).

11.	Benjamini, Y. & Yekutieli, D. The control of the false discovery rate in multiple testing under dependency. *Ann Stat* **29**, 1165-1188 (2001).
